# Genetic population structure across Brittany and the downstream Loire basin provides new insights on the demographic history of Western Europe

**DOI:** 10.1101/2022.02.03.478491

**Authors:** Isabel Alves, Joanna Giemza, Michael Blum, Carolina Bernhardsson, Stéphanie Chatel, Matilde Karakachoff, Aude Saint Pierre, Anthony F. Herzig, Robert Olaso, Martial Monteil, Véronique Gallien, Elodie Cabot, Emma Svensson, Delphine Bacq-Daian, Estelle Baron, Charlotte Berthelier, Céline Besse, Hélène Blanché, Ozvan Bocher, Anne Boland, Stéphanie Bonnaud, Eric Charpentier, Claire Dandine-Roulland, Claude Férec, Christine Fruchet, Simon Lecointe, Edith Le Floch, Thomas Ludwig, Gaëlle Marenne, Vincent Meyer, Elisabeth Quellery, Fernando Racimo, Karen Rouault, Florian Sandron, Jean-Jacques Schott, Lourdes Velo Suarez, Jade Violleau, Eske Willerslev, Yves Coativy, Mael Jézéquel, Daniel Le Bris, Clément Nicolas, Yvan Pailler, Marcel Goldberg, Marie Zins, Hervé Le Marec, Mattias Jakobsson, Pierre Darlu, Emmanuelle Génin, Jean-François Deleuze, Richard Redon, Christian Dina

## Abstract

European genetic ancestry originates from three main ancestral populations - Western hunter-gatherers, early European farmers and Yamnaya Eurasian herders - whose edges geographically met in present-day France. Despite its central role to our understanding of how the ancestral populations interacted and gave rise to modern population structure, the population history of France has remained largely understudied. Here, we analysed the high-coverage whole-genome sequences and genome-wide genotype profiles of respectively 856 and 3,234 present-day individuals from the northern half of France, and merged them with publicly available present-day and ancient Europe-wide genotype datasets. We also explored, for the first time, the whole-genome sequences of six mediaeval individuals (300-1100 CE) from Western France to gain insights into the genetic impact of what is commonly known as the Migration Period in Europe. We found extensive fine-scale population structure across Brittany and the downstream Loire basin, emphasising the need for investigating local populations to better understand the distribution of rare and putatively deleterious variants across space. Overall, we observed an increased population differentiation between the northern and southern sides of the river Loire, which are characterised by different proportions of steppe vs. Neolithic-related ancestry. Samples from Western Brittany carry the largest levels of steppe ancestry and show high levels of allele sharing with individuals associated with the Bell Beaker complex, levels that are only comparable with those found in populations lying on the northwestern edges of Europe. Together, our results imply that present-day individuals from Western Brittany retain substantial legacy of the genetic changes that occurred in Northwestern Europe following the arrival of the Bell Beaker people c. 2500 BCE. Such genetic legacy may explain the sharing of disease-related alleles with other present-day populations from Western Britain and Ireland.

## Introduction

Understanding how genetic diversity is distributed within and between human populations (*i.e.* population structure) can shed light on the history of our species, and inform studies that search for genetic associations with traits and diseases (1). The increasing interest in elucidating the role of rare variation in complex traits, the consequent demand for surveying multiple local populations and the need for a detailed understanding of their evolutionary history has motivated multiple population genomic studies at a country-wide scale (2–5).

Located on one of the edges of the European landmass, Northwestern France includes the peninsula of Brittany, whose westernmost tip is named *Finistère* (*end of land*), and the surrounding region of *Pays-de-la-Loire*. The region is bordered by the English Channel to the north and the Bay of Biscay to the south, and is crossed by the Loire River -the largest river in France. From a linguistic point of view, this region has been the stage of long-term contacts between Celtic and Gallo-Romance spoken forms. The first evidence of modern human occupation of Northwestern France is associated with the Early Aurignacian culture, dating between about 43,000-37,000 years ago (6). However, population occupation in the region may have been scarce up to the Middle/Late Neolithic. It is during this period that megalithic construction appears in the archaeological record (6). The peninsula of Brittany hosts the oldest (7) and some of the biggest megalithic monuments (stone rows, long barrow and passage tombs) and one of the highest concentrations thereof. The Neolithic culture in Northwestern France largely derives from the Danubian Wave, also known as the Linear Pottery culture (8,9), although the presence of pottery with shell impressions in regions south of the Loire river raises the possibility of Mediterranean influence in the neolithization of the region (9). Genetic evidence suggests that the introduction of the France Neolithic lifestyle occurred through a complex interaction between the two Neolithic waves - the Danubian and the Mediterranean wave - and local, likely genetically structured, populations of hunter-gatherers (10–12). Both archaeological and genetic evidence suggest strong connectivity among the populations along the Atlantic façade from 4500 BC onwards, from Northern Iberia to Ireland and Western Britain (13). Although Northwestern France was likely part of the Atlantic façade with respect to its genetic landscape, there is no ancient DNA data currently available from the Neolithic period.

The connectivity along the western fringes of the European continent continued during the Early Bronze Age as suggested by the presence of a common Bell Beaker pottery style, usually known as the “Maritime” style (14,15), and further related artefacts (16). In addition, the Bell Beaker complex in Northwestern Europe is closely associated with the appearance of copper metallurgy (17). In Britain, the arrival of the Bell Beaker complex has been recently associated with a major genetic ancestry shift disrupting the preceding genetic homogeneity of the Atlantic façade and introducing genetic ancestry related to the Yamnaya migration from the Eurasian Steppes (18). These findings contrast with what has been found in Iberia, where Beaker- complex-associated individuals show low genetic affinity with those from central Europe suggesting considerable heterogeneity underlying the mode of transmission of the Bell Beaker complex. Within France, Yamnaya-related ancestry has been primarily found among Bell- Beaker associated individuals from the Northeastern and Southern parts of the country, although ancestry proportions widely vary across samples (10,11). However, the lack of samples from Northern and Northwestern France before the Iron Age hinders the full understanding of the introduction of Steppe ancestry along the westernmost part of the North Sea and the English Channel. In Northern France, the Iron Age is genetically characterised by high levels of Steppe-ancestry and a homogenization of it, suggesting continuous cultural change instead of a massive migration underlying the arrival of iron technology (19). In the Iron Age, Brittany was called *Aremorica* and was part of the Gaul. It was home to multiple Celtic-speaking tribes, like the Veneti and Osismii in the west of the peninsula (20).

During the Roman Empire, Roman influence reached Brittany, as attested by the presence of Roman-style villas and sanctuaries. However, such influence was far smaller in Northwestern France than in other parts of the Gaul (21). With the decline of the Roman Empire (around the 3rd century CE), Northern France became progressively ruled by the early mediaeval kingdoms (22). However, the history of Brittany remains largely unknown from the late Antiquity (4th- 5th century CE) up to the 9th century CE, when it was conquered by Frankish Carolingian Emperors and put under native rulership (23). Interestingly, it was during this period that the peninsula acquired the name of “Britannia”. Linguistically, places’ names and the still-surviving language closely related to Cornish and Welsh imply that extensive connections took place with Western Britain. Nevertheless, whether such a process involved small or large settlements from the British Isles, as often suggested, or is the basis of the Breton language is still under debate (24–27). Archaeological evidence displays changes in architecture and funerary rituals during this time. However, they reflect a mixture of influences, rather than a single major contribution, from the British Isles and French regions along the British Channel as well as from Germany (28).

Northwestern France has been the end point of multiple continental prehistoric migrations and occupied a central place in trading routes along the Atlantic façade (29). Understanding the genetic makeup of Northwestern France can inform on the complex interaction between the European-wide demographic events that have culminated in the present-day genetic landscape. However, a systematic assessment of patterns of population structure in Northwestern France and an identification of the past demographic events that have shaped them is still lacking. In this study, we analysed 3,234 genome-wide genotyped samples together with 620 high-coverage whole-genome sequences (WGS) from Brittany and *Pays-de-la-Loire*. Also, to put the genetics of Northwestern France into context we used an additional set of 236 WGS from other regions in France and merged the WGS with available datasets encompassing a geographically diverse set of present-day and ancient European samples. Last but not least, we analysed, for the first time, a set of western French mediaeval samples dated from c. 400 to 1100 CE in order to fill in the gap with respect to the lack of publicly available genome-wide data from the last two millennia.

## Results

### Haplotype-based population structure in Northwestern France

To explore patterns of genetic structure among the population in Northwestern France, we first applied the haplotype-based approach implemented in fineSTRUCTURE (vs4.1) (30) on 210,171 SNPs genotypes from 3,234 individuals from Brittany and *Pays-de-la-Loire* (see Methods). Genome-wide data revealed extensive population structure in Northwestern France with 154 clusters inferred, of which 78 contain more than 10 individuals (Fig. S1.1). At this finest scale, levels of clustering were similar to those previously found for Spain (3), and larger than those previously reported for France (31) and Great Britain (5). We investigated whether recent ancestry due to the sampling scheme carried out in this study impacts the clustering patterns and we found no evidence of such an effect (see Methods and Fig S1.2-S1.4). Pairs of individuals within clusters are not more closely related than pairs of individuals across the whole dataset.

At the coarser level k=3 (Fig. 1a) of the fineSTRUCTURE tree (FS tree), we found that the distribution of one of the clusters, hereafter referred to as the “Western Brittany” (WBR) cluster, broadly overlaps with the linguistic distribution of the Breton language. The cluster’s border falls between the easternmost (Loth line 1) and the westernmost (Loth line 2) estimated historical bounds of the Breton language (Fig. 1b). A second cluster, hereafter referred as “Eastern Brittany and Pays-de-la-Loire” (EBP), encompasses individuals from eastern Brittany (*département* Ille-et-Vilaine) and the area of *Pays-de-la-Loire* situated north of the river Loire. A third cluster, hereafter referred to as “South Loire” (SLO), covers the part of the *Pays-de-la- Loire* situated south of the river Loire.

**Figure 1.**
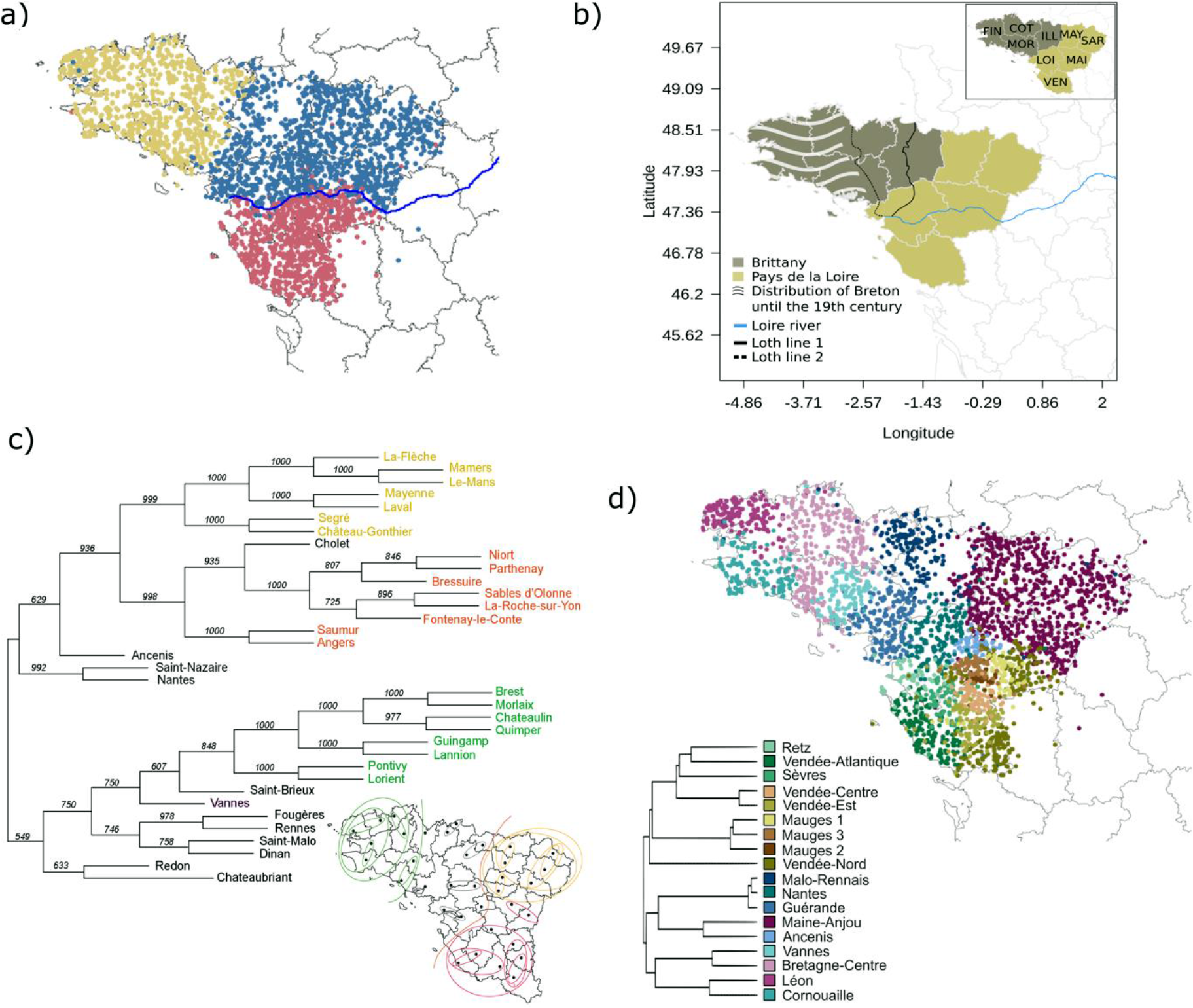
Population structure, linguistics, and patronym distribution in Northwestern France **(a)** fineSTRUCTURE clustering of 3,234 samples from Brittany and the downstream Loire basin at k=3. **(b)** Distribution of the Breton language over time (adapted from the *Celtic languages Donald MacAulay Cambridge University Press page 373*). Loth line 1, attested by the border between Celtic place names in –ac (on the west side) and Romance place names in –é (on the east side), represents the westernmost boundary between Armorican Gaulish and Gallo-Romance spoken varieties (25). Loth line 2 represents the Celto-Romance linguistic border in the 16th century CE. Grey lines encircle political territorial divisions called *départements*. Only *départements* belonging to the regions of Brittany and *Pays-de-la-Loire* are coloured. In the inset, the *départements* are abbreviated as FIN (Finistère), COT (Côtes- d’Armor), MOR (Morbihan), ILL (Ille-et-Vilaine), MAY (Mayenne), SAR (Sarthe), MAI (Maine-et-Loire), LOI (Loire-Atlantique), VEN (Vendée). **(c)** Neighbour Joining tree from pairwise Arccos distances (68) computed on the distribution of surnames between pairs of *arrondissements* from Northwestern France. The support of the tree nodes was obtained by bootstrapping 1,000 times the distance matrices. **(d)** 18 genetic clusters identified using total variation distances (TVD) from the fineSTRUCTURE results (see Material and Methods for details). The TVD-based tree was cut at k=39 and to ease visualisation we merged clusters with <5 individuals into the closest cluster (>=31 individuals) resulting in a tree with 18 clusters. All the clusters have at least 31 individuals. The TVD-based tree is sample size independent and is more informative on cluster genetic affinities (31). In (a) and (c), individuals are coloured according to their cluster assignment. Individuals’ coordinates correspond to the most common grandparents’ birthplace.

The relative distributions of these clusters mimic those of surnames across the entire region, as inferred through a Neighbour Joining Tree (Fig. 1c). The three surname-based clusters are highly supported by the bootstrap analysis (> 75%). Interestingly, the surnames from the northern part of *Pays-de-la-Loire* are more closely related to those from south Loire than to those from Brittany. Furthermore, while the highest level of differentiation is observed between WBR and SLO (F_ST_=0.00107), F_ST_ values indicate larger population differentiation between EBP and SLO than between EBP and WBR (respectively 0.00042 vs. 0.00028, Fig. S1.5). Together we found evidence that the river Loire influences population differentiation, likely acting as a barrier to gene flow.

For higher levels of clustering resolution (k > 3), because the FS tree depends on the sample sizes (30) and might not reflect haplotype sharing differences between pairs of clusters (32), we computed the clustering tree based on the total variation distances (TVD tree, (5,32)). Indeed, when assessing the performance of the two tree-building algorithms (see SOM for details) by computing cluster assignment confidence (5), we found that for the same k the FS- tree shows lower cluster assignment confidence than the TVD-based tree (Fig. S1.6). We chose k=39 and to ease visualisation we merged small-sized clusters (1-5 individuals) with the closest large cluster (>=31 individuals) resulting in 18 clusters (>31 individuals each). As a result, the north Loire region splits into nine clusters, four of which in the westernmost part of Brittany, while individuals from south Loire split also into nine clusters (Fig. 1d). Overall, we retrieved a TVD-tree largely consistent with the fineStructure tree at k=3, with the exception of samples located at the borders between the three main clusters (e.g., individuals assigned to the Malo- Rennais cluster of the TVD-tree)”. In the resulting 18 clusters, individuals from the westernmost part of Brittany are grouped into four clusters: “Cornouaille”, “Lèon”, “Vannes” and “Breatagne-Centre”, while individuals mostly from south of the Loire river cluster into nine clusters (Fig. 1d). Samples geographically located between these regions are grouped into five clusters: “Malo-Rennais”, “Nantes”, “Gérande”, “Maine-Anjou” and “Ancenis”. The coancestry matrix for the 18 clusters (Fig. S1.6) reveals a global pattern of increased coancestry within clusters, as expected after a history of shared drift between individuals in the same group, with the exceptions of the clusters ‘Sèvre’ and ‘Vendée-Est’.

The cluster “Maine-Anjou’’ encompasses the largest number of individuals (n=799) and covers by far the largest geographical area, revealing broader genetic homogeneity relative to the other regions. Most of the other clusters appear to have contributed to the coancestry of this large cluster, indicating that cluster haplotype homogeneity may result from repeated admixture into the region (Fig. S1.7).

The second largest cluster (n=344), “Bretagne-Centre”, stretches from the northern to the southern coast of Brittany indicating the existence of a corridor of relative genetic homogeneity connecting the two sides of the peninsula (see SOM for details). While the neighbouring clusters “Leon”, “Cornouaille’’ and “Vannes” display large within-cluster but low between-cluster coancestry (Fig. S1.7), despite their close geographic proximity, each of them appears to have contributed to the coancestry of the “Bretagne-Centre” cluster. When considering the area located south to the river Loire, most clusters– and particularly “Mauges 1” and “Mauges 2” - appear to have largely contributed to the coancestry of the “Sèvres” cluster in eastern Vendée. In contrast, the “Mauges [1-3]” and “North-Vendée” clusters show low coancestry from most of the other clusters and large within-cluster coancestry, likely indicating stronger genetic drift within these groups. Finally, we observed a low contribution of the clusters south of the river Loire to the coancestry of those from the westernmost part of Brittany, indicating that gene flow between these two areas has been relatively limited, in agreement with the F_ST_ values obtained between the 18 clusters (Fig S1.8).

We then explored identity-by-descent (IBD) sharing between pairs of individuals belonging to the 18 clusters (Fig S1.9), and measured average length of runs of homozygosity (ROH) across individuals within administrative circumscriptions in Northwestern France (Fig. S1.10). Higher levels of identity-IBD sharing for chromosomal segments of any size as well as increased length of ROH were observed between the clusters “Mauges [1-3]” and “Vendée-Nord”, indicating that low effective population sizes may explain the particular population structure found in this restricted area. Similar patterns were observed within Brittany. However, considering that increased IBD sharing within the westernmost clusters of Brittany is only observed for chromosome segments under 7cM (Fig. S1.9), lower effective population sizes within this cluster are likely more ancient than for “Mauges [1-3]” and “Vendée-Nord”. Overall, these results fit with those based on the coancestry matrices and F_ST_ values (Fig S1.8).

### Relationship between fine genetic structure, linguistics and geography

In order to elucidate whether particular cultural features can explain the observed genetic structure, we explored spatial language and surname distribution potentially coinciding with cluster borders. First, we found that the cluster “Bretagne-Centre” overlaps largely with the dialectal area featured by the use of two initial consonants - the aspirated [h] instead of an unaspirated, and the alveolar fricative [z] instead of [s] (Fig. S1.11 and Supplementary Data). Similarly, we observed substantial overlap between the “Cornouaille” cluster and the area with palatalization of -h- [h] into -y- [j] (Fig. S1.12; see Supplementary data). Second, we observed high correlation between pairwise F_ST_ values and surname-based distances (Fig. S1.13), even after correcting for geographical distances (Partial Spearman correlation = 0.68). To illustrate this finding, the “Leon” and “Malo-Rennais” cluster locations match those of the surname clusters Brest/Morlaix and Saint-Malo/Dinan, respectively (Fig. 1c). Overall, these observations support the hypothesis that language has played a role in shaping the genetic structure currently observed in Northwestern France.

In parallel, we checked whether rivers other than the Loire could have driven population differentiation at a finer scale, and observed repeated overlap between river courses and cluster borders (Fig S1.14). For instance, the clusters “Lèon” and “Cornouaille’’ are respectively delimited to the east by the Morlaix River and Laïta-Ellé rivers, and separated from each other by the river Aulne. “Vannes” is enclosed to the west by the Blavet river and to the north by the Oust. The “Nantes” cluster is bordered by the rivers Semnon and Vilaine to the north and the west, respectively, while the “Malo-Rennais” cluster’s borders co-localize with the Gouessant and the Yvel - Hivet to the west. In South Loire, the cluster “Vendée-Atlantique” is bordered to the southwest by the river Lay (Fig. S1.14). Finally, when inspecting clusters at k=78 within the area of “Maine-Loire”, we could observe recurrent proximity between cluster borders and the waterways of Mayenne, Vègre and Loir (data not shown). These recurring coincidences of watercourses with genetic cluster borders, while not formally tested, may reflect the impact of rivers on local demographics.

### Evolution over time of effective population sizes

We estimated the historical dynamics of effective population size (Ne) using IBDNe (33), and observed 12-, 5- and 5.2-fold increases in Ne, respectively for WBR, EBP and SLO, in the last 1,450 years (assuming a generation time of 29 years; Fig. S1.15a). Such expansions have led to present-day population sizes of around 10^5.5^ to 10^5.7^ individuals for the three clusters, which is broadly consistent with the explosive population growth estimated for other European populations (34–36). However, the Ne trajectories differ between clusters (Fig. S1.15a). Indeed, while WBR shows the smallest Ne in the Early Mediaeval Period (∼5th century CE), its effective population size surpassed that of SLO by the High Mediaeval Period (∼1000 CE). This suggests that population expansion started earlier in WBR than in SLO, which kept a stable Ne until the 10 last generations (i.e., 17th century CE). Reduced Ne at recent times explain the stronger IBD sharing observed between the clusters “Mauges [1-3]” and “Vendée-Nord” for chromosome segments > 7cM. Finally, the Ne trajectories in EBP and SLO show a slight and short population decline, starting around 1230-1350 CE and lasting for almost ∼300 years (Fig. S1.15a). Similar patterns have been previously described in French and other European populations and putatively associated with the Black Death (31,34). However, we caution that the Ne profiles suggesting a bottleneck are not consistently observed across different thresholds for the minimum chromosome length (Fig S1.15b-d). In addition, our computer simulations suggest that multiple demographic scenarios (e.g., population structure within the EBP and SLO samples) generate similar Ne profiles (see SOM for details, Fig. S6). Historical evidence that the Black Death had a lesser demographic impact in Western Brittany is lacking, although recent studies refining its impact through pollen record support a lighter effect of the epidemics in Brittany (37).

### Fine-scale population structure based on rare variants

Population stratification based on rare variation is typically stronger than with common polymorphism (1,38) and rare variants are particularly informative to infer recent fine-scale population structure. Here, we computed allele sharing between pairs of individuals originating from Brittany and *Pays-de-la-Loire* using genotypes from 620 high-coverage whole-genome sequences (see Methods). We computed independently two matrices, reporting genotypes with minor allele counts equal to two (MAC 2) and ranging from 3 to 10 (MAC 3-10), respectively. We applied hierarchical clustering to both matrices to cluster individuals according to patterns of allele sharing. When assuming three population groups (*k*=3), most individuals from the three westernmost *départements* of Brittany are clustered into a single group, both for MAC 2 and MAC 3-10 (Fig. 2). The proportion of individuals assigned to this cluster decreases as one moves away from Western Brittany, regardless of the allele count category, and the two alternative clusters become more prevalent. Similarly, to the fineSTRUCTURE findings, these results provide evidence for relative differentiation between traditionally Breton-speaking populations in Western Brittany and their Gallo-speaking neighbours.

**Figure 2.**
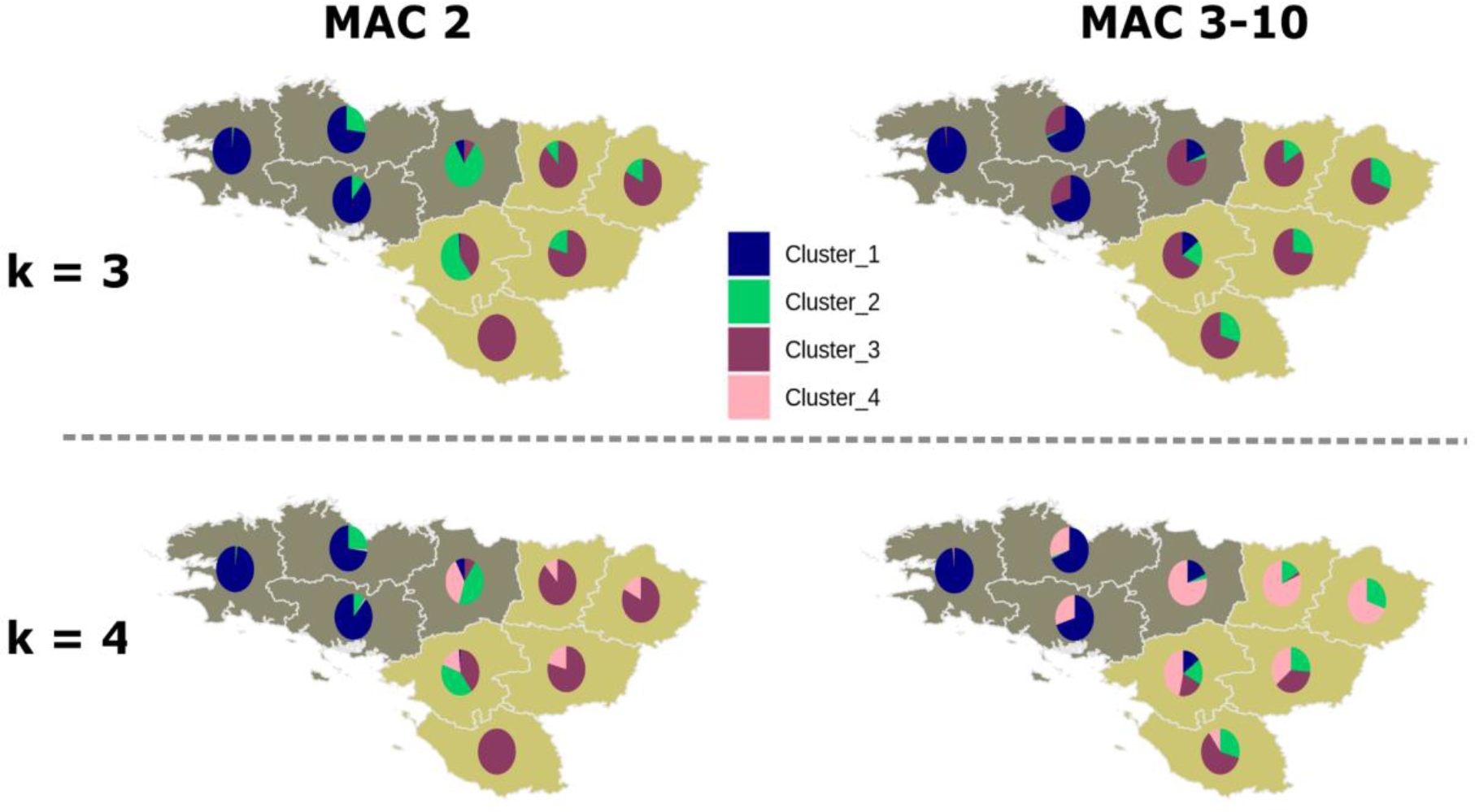
Allele sharing within and between populations. Proportions of individuals assigned to different clusters for each *département* in Northwestern France (see Fig. 1b), using hierarchical clustering on levels of rare allele sharing between pairs of individuals. Allele-sharing matrices were computed for alleles present in two chromosomes (MAC 2) and three to 10 chromosome copies (MAC 3-10) across individuals from Northwestern France and from one million randomly selected sites.

Interestingly, we identified a genetic component restricted to the *départements* located south to the river Loire for *k*=4 (cluster_2) for MAC 3-10 (Fig. 2, Fig. S2.1). For MAC 2 alleles we found this cluster only at *k*=9 (cluster_2). Assuming that alleles with MAC 2 (minor allele frequency ≈ 0.0016) tend to be more recent, these results suggest that population structure does not result from reduced gene flow between the northern and southern shores of the river in the very near past (average doubleton age ∼500 years (39)). Consistently, we found no significant differences in surname distributions between the riversides across *arrondissements* crossing the Loire (data not shown). Clustering patterns within Brittany are, on the other hand, consistent across the full range of allele counts, indicating that population differentiation associated with traditionally Breton-speaking groups has persisted to modern times.

With MAC 2 alleles, increasing *k* from 3 to 10 assign individuals from neighbouring *départements* into 7 additional geographically restricted clusters (Fig. S2.1), suggesting similar patterns of population structure as found with fineSTRUCTURE (Fig. 1d). Although an exhaustive comparison with fineSTRUCTURE results is beyond the scope of this study, this general concordance in clustering patterns emphasises the power of rare variants to infer fine-scale population structure. With MAC 3-10 alleles, increasing *k* tends to generate smaller clusters with relatively large geographical distribution, likely reflecting a relative lack of resolution to detect population structure (Fig. S2.1).

### Brittany in the context of France

To investigate population structure on a larger geographical scale, we enriched our dataset with additional WGS-based genotypes from 233 individuals originating from neighbouring French areas (see Methods). We first performed principal component analysis (PCAs) based on common and low-frequency variants (Fig. S2.2). While population differentiation based on common variants separates mostly Brittany from the remaining regions, low-frequency variants disclose a more subtle population structure, as previously reported (38,40), with additional separation between Northeastern and Southwestern France (for further details, refer to Génin *et al.,* manuscript in preparation). Pairwise F_ST_ values also support stronger differentiation between WBR (Fig. 1d) and the remaining populations (Fig. S1.5). In subsequent analyses, we will consider individuals from Brittany and *Pays-de-la-Loire* according to the three clusters shown in Fig. 1a (WBR, EBP and SLO), unless stated otherwise.

### Brittany in the European context

To further investigate the genetic history of people from Northwestern France, we first performed a PCA on our entire WGS dataset merged with genotype data from 20 diverse Northern and Western European populations (41,42). In agreement with the previously reported isolation-by-distance pattern in Europe (43), we found that French samples are continuously distributed along the axis connecting Southwestern (i.e., Spain) and Northwestern (i.e., Ireland/UK) Europe (Fig. 3a). Individuals from Central/Southwestern France appear closer to samples from Spain whereas individuals from Brittany appear at the other extreme and closer to samples from Ireland/UK. While individuals from Brittany fall onto the axis and overlap with the Irish, Welsh and Cornish samples, samples from Eastern Great Britain show a slight shift towards Central Europe. These results support the idea of increased genetic proximity between Brittany and Ireland, as previously suggested (44). In contrast with Brittany and similarly to what we observe for samples from Eastern Great Britain, individuals from Northern and Eastern France show a slight shift towards Central Europe (represented here mainly by Germany).

**Figure 3.**
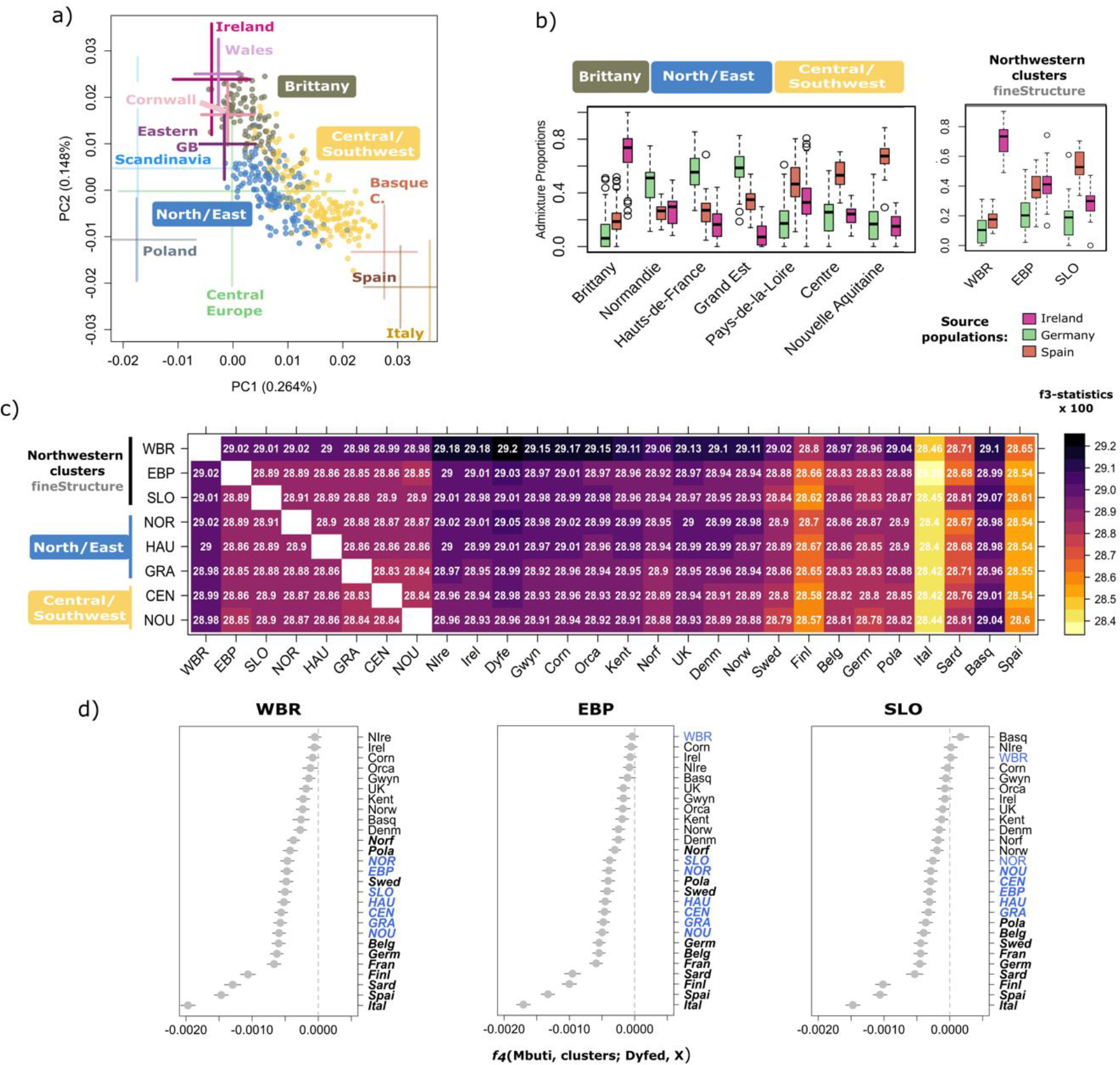
Genetic affinities between Brittany and Ireland/West British Isles **(a)** The two first principal components of genetic variation in the “modern merged dataset”, which includes 843 French WGSs for which the four grand-parent origin was concordant, and genome-wide data from 20 central and western European populations (population acronyms below, see Methods for further details). To avoid bias due to sample size differences among regions all locations are represented by a maximum of 100 individuals resulting in a total of 2,070 samples and 201,999 independent SNPs. For simpler visualisation samples from Norway, Sweden and Denmark were labelled as “Scandinavia”, samples from Belgium and Germany were labelled as “Central Europe”. FranceGenRef samples are represented by dots and labelled with boxes while non-French samples are represented by 2sd of the two PCs. UK samples other than those from the POBI dataset, whose origin of the grandparents is known, are not shown. Similarly, French samples from public datasets are also not shown. To better visualise the distribution of our samples the plot is a zoom of the full PCA (Fig. S3.1). **(b)** Ancestry proportions retrieved from a Supervised Admixture analysis considering Ireland, Spain and Germany as proxy source populations on the basis of their polarised positions relative to the distribution of the French samples on panel (a). **(c)** Heatmap reporting outgroup *f3-statistics* in the form f3(Mbuti; French population/cluster, X), where French population/cluster is represented on the y-axis and X on the x-axis. **(d)** *f4*-statistics of the form *f4*(Mbuti, French cluster; Dyfed, X), where X are the populations on the right side of the plot. Due to disparities in samples sizes, both the outgroup *f3*- and *f4*-statistics were computed on a maximum of 25 samples per population. In bold are represented *f4-statistics* values with a corresponding |Z| > 3. Cluster acronyms: western Brittany (WBR), eastern Brittany and *Pays-de-la-Loire* (EBP), south Loire (SLO). Region/country acronyms: Brittany (BRI); *Pays-de-la-Loire* (PAY); *Normandie* (NOR); *Hauts-de-France* (HAU); *Grand Est* (GRA); *Centre-Val de Loire* (CEN); *Nouvelle-Aquitaine* (NOU), Basque Country (Basq), Belgium (Belg), Cornwall (Corn), Denmark (Denm), Dyfed (Dyfe), Finland (Finl), Germany (Germ), Gwynedd (Gwyn), Ireland (Irel), Italy (Ital), Kent (Kent), Norfolk (Norf), Northern Ireland (NIrel), Norway (Norw), Orkney Islands (Orca), Poland (Pola), Sardinia (Sard), Spain (Spai), Sweden (Swed), United Kingdom (UK).

To further investigate the contribution of other European populations to the genetic makeup of Northwestern France we used the regression-based statistical approach implemented in GLOBETROTTER software (45). We found that all of the seven French populations showed evidence of admixture (*P* < 0.0001). In agreement with the PCA results, three main sources of ancestry – North/Central, Northwestern and Southwestern Europe - were consistently found across these seven populations. The North/Central component derives mainly from Denmark and/or Belgium, the Northwestern component from Ireland and the Southwestern one from Spain and/or Italy (Fig. S3.2). While *Hauts-de-France* (HAU) and *Grand Est* (GRA) ancestries show evidence of single admixture events, with predominant and similar contributions from the North/Central and the Southwestern components, the remaining populations (BRE, NOR, PAY and NOU) show evidence of multiple waves of admixture (the best-guess model could not be inferred for CEN). PAY (*Pays-de-la-Loire*) and NOU (*Nouvelle-Aquitaine*) show evidence of mixed ancestries of predominant Southwestern European origins and the population of Brittany (BRI) was found to trace back ∼23.5% of its DNA to Ireland vs. 14% or less found in any other French population. Brittany is also the region where Cornwall has contributed the most to ancestry (∼9% vs. 3.1-3.4% across the remaining populations).

To confirm the proportions of ancestry in relation with these external groups, we performed supervised clustering considering Ireland, Germany and Spain as putative sources of ancestry for the French samples. We found that the Irish component accounts for >75% of the ancestry in ∼50% of the individuals from Brittany (Fig. 3b), while it contributes to a much lesser extent to the genetic composition of the other regions (8-33%). On the other side of the spectrum, a Spanish-like component accounts for the largest proportion (∼70%) in *Nouvelle-Aquitaine* (NOU), while German-like ancestry is predominant among individuals from Northern and Eastern France (*Normandie,* NOR; *Hauts-de-France,* HAU; *Grand Est*, GRA), with its proportion increasing with the geographical proximity to the German border.

When considering the three clusters inferred by fineSTRUCTURE in Northwestern France, we observe the same ancestry pattern for WBR as for Brittany, with a major Irish component. Consistently with this result, the smallest F_ST_ values between the WBR group and other non-French populations were retrieved with the Irish and Northern Irish populations (0.00057 and 0.00062, respectively, Fig. S3.3). The only French population showing lower pairwise F_ST_ with WBR is the neighbouring EBP (F_ST_=0.00028, Fig. S1.5). Finally, among the three clusters, SLO carries the largest Spanish-related ancestry while EBP carries similar proportions of the Irish- and Spanish-related components, reflecting its intermediary position between WBR and SLO. In summary, our results indicate that people from Brittany show strong genetic affinities with populations from Western Britain and Ireland, although separated by the Celtic sea.

To measure the relationship between the genetic clusters found in Northwestern France and other European populations, we used the *outgroup f3*-statistics to assess the genetic drift shared by pairs of populations relative to the outgroup population (Mbuti)(46,47). Given that this statistic reflects the length of the branch from the internal node to the outgroup (connecting the pair of populations being tested), it is not affected by lineage-specific genetic drift, contrarily to F_ST_. We found that most French clusters located north of the river Loire share the largest drift with Southwestern Welsh populations (i.e., from Dyfed), whereas those located south to the river Loire share the largest drift with the Basques (Fig. 3c, Table S1.1). The *f4-statistics* of the form *f4*(Mbuti, French subgroup; Dyfed, X), which should produce significantly positive values when the tested population shares more alleles with X than Dyfed, show that the French subgroups from areas south to the river Loire consistently share more alleles with the Basques than with the Southwestern Welsh. Conversely, those located north to the river Loire share the largest amount of alleles with Southwestern Welsh and other populations from Great Britain and Scandinavia (Fig. 3d, Fig. S3.4).

### Genetic continuity since mediaeval times

To disentangle the sources of ancestry contributing to the modern French genetic makeup, we merged our WGS data with the largest available compilation of ancient (>3000 samples including ∼400 ancient Vikings) and modern samples (>5000)(48,49). In addition, we sequenced six individuals with dates ranging from the 4th to the 12th century CE, from *Pays-de-la-Loire* (Fig. 4a) to increase our resolution in detecting changes in ancestry during the Mediaeval Period. PCA resulting from projecting the ancient individuals onto the principal component space of modern variation shows that most of the samples fall well within the distribution of present-day French (Fig. S4.1). Out of the six individuals, one (fra009, Table S2.1) likely represents a migrant with genetic affinities to present-day North Africans. This individual, dated from the 5th-6th century CE, was found in an archaeological site located in an ancient town likely built during the Roman period (see SOM, Supplementary archaeological details). Trading networks involving this town may explain the presence of North African migrants so far north. To test whether French Mediaeval samples from the 3-4th century CE and samples from the 6-7th century CE significantly differ in their genetic affinities to other ancient European populations we computed the *f4*-statistics of the form *f4(Mbuti, ancient European sample; sLoire_France_3-4cCE, sLoire_France_6-7cCE).* We found no significant differences in allele sharing between individuals from early (300-550 CE, fra001 and fra004) and later Mediaeval Period (600-700 CE, fra016 and fra017, Table S2.2). Therefore, we considered individuals from both periods to represent the same population and refer to them as “Mediaeval French”.

**Figure 4.**
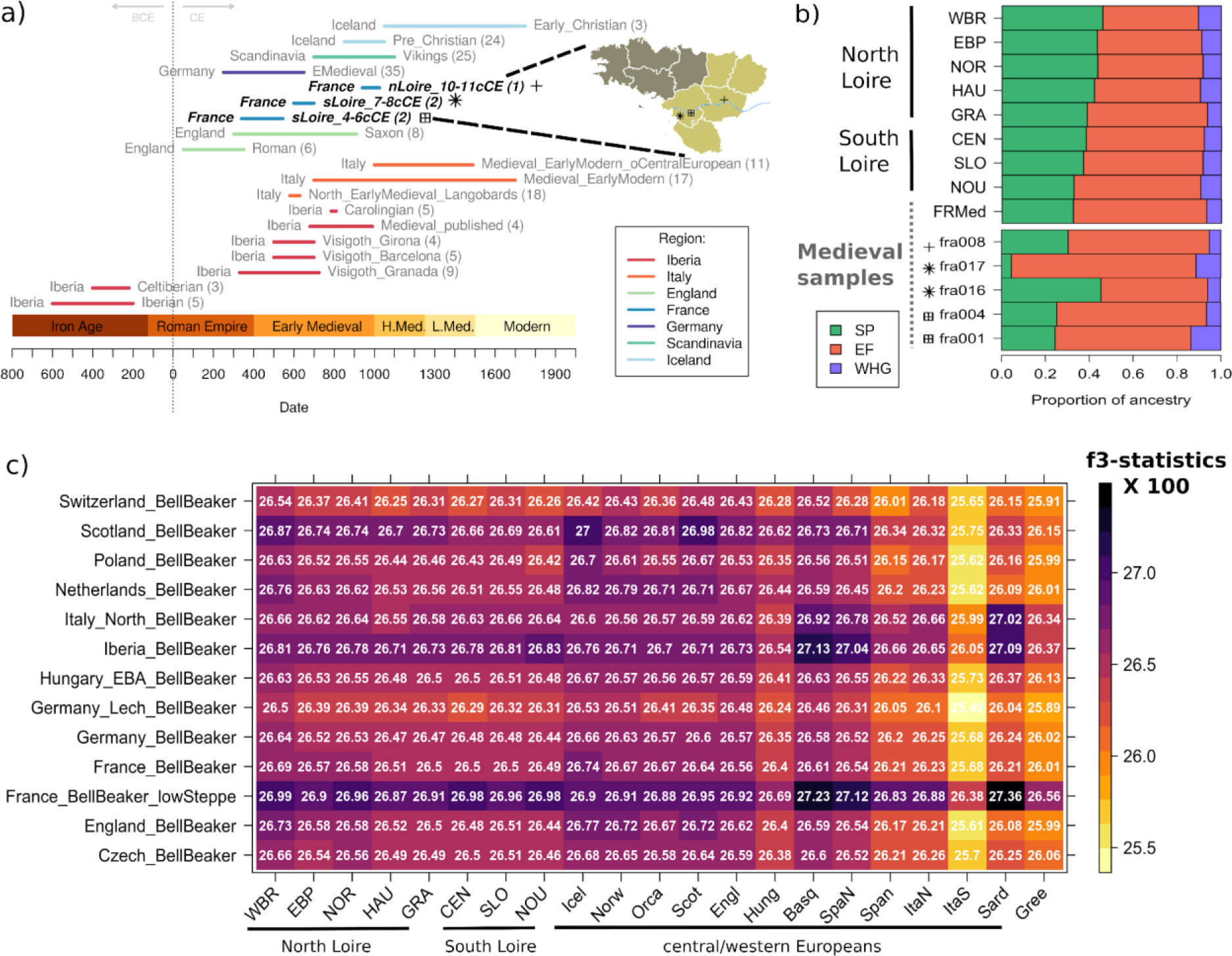
Genetic continuity, admixture and relationship between modern and ancient French populations **(a)** Timeline of the ancient individuals used to test one- and two-way models of admixture to explain the ancestry of present-day French using *qpAdm*. **(b)** Mixture proportions from the three major populations contributing to the European ancestry using three-way admixture model, where each population on the left (y-axis) is modelled as a mixture of Mesolithic Western Hunter-Gatherer- (WHG), Neolithic early farmers- (EF) and Early Bronze Age Steppe- (SP) derived ancestry. **(c)** Shared drift between Bell-Beaker-associated individuals and present-day Europeans by means of the outgroup *f3*-statistics of the form *f3*(Mbuti; Bell Beaker Pop, X), where X is the population on the x-axis. These analyses were performed on the “ancient merged dataset” (see Material and Methods for details).

To check for ancestry changes since the Mediaeval Period, we tested whether Mediaeval French individuals are a good proxy source for the ancestry of present-day Northwestern French. To do so we used the modelling approach implemented in *qpAdm* (50) and we tested one-way models (i.e., genetic continuity) using the five Mediaeval French individuals as surrogate source population for the present-day French. Our results support genetic continuity between the eight present-day French populations (WBR, EBP, SLO, NOR, HAU, GRA, CEN and NOU) and Mediaeval French (*P* > 0.05, Table 1). *qpAdm* modelling results are consistent with the PCA showing that Mediaeval French from *Pays-de-la-Loire* fall within the distribution of modern individuals from the same region. Due to constraints in sample overlap between our SNP-array and WGS datasets, the randomly selected individuals within WBR do not cover the full geographic distribution of the initial cluster (*départments:* Morbihan, Finistère and Côtes- d’Armor) with the *départment* of Finistère being mostly underrepresented. Hence, we tested a one-way model using only individuals from the westernmost *département* (*Finistère*) of Brittany and we found evidence (*P* = 0.0169) for no continuity between Mediaeval samples and present-day individuals from Finistère. Due to the fact that model fit *p-values* can be affected by factors such as sample size and coverage, *p-values* comparison should be avoided. Nevertheless, given that we kept the same sample sizes (n=25 with the exception of NOR with n=19) and differences in coverage should be minimal due to the quality of our sequencing, it is tempting to argue that the lack of continuity when using only samples from Finistère might reflect ancestry variation already present during Mediaeval times. An alternative scenario could, for instance, include later migrations from a non-French source likely through the sea. We also found that one-way models using ancient Northwestern Europeans, such as those dating of the Roman period in Great Britain, fit the ancestry of French populations north of the Loire (0.074 < *P* < 0.936; WBR, EBP, NOR, HAU and GRA; Table S2.3) but do not fit the ancestry of those south of the river (*P* < 0.05, CEN, SLO and NOU). Furthermore, Italians dating from the early modern period with a central European ancestry fit modern French from south of the river Loire (*P* > 0.05). Interestingly, ancient Spanish reported as Celtiberians or from the Mediaeval period do not generally fit contemporary French (with the exception of NOU) and only Spanish individuals associated with the Germanic invasions, especially individuals associated with Visigoth or Carolingian archaeological remains, are suitable proxies for contemporary French ancestry (Table S2.3).

**Table 1.** *qpAdm* results for one-way and two-way models used to estimate the ancestry of Mediaeval and present-day individuals from France. Only models with p-value > 0.05 are reported.

| Target | Source 1 | Source 2 | p-value | Prop 1 | Prop 2 | std error |
| --- | --- | --- | --- | --- | --- | --- |
| WBR | FRMedieval | - | 0.0679 | - | - |  |
| WBR | FRMedieval | English | 0.5338 | 0.009 | 0.991 | 0.262 |
| WBR | FRMedieval | England_Roman.SG | 0.9026 | 0.061 | 0.939 | 0.230 |
| WBR | FRMedieval | Norwegian | 0.9645 | 0.333 | 0.667 | 0.080 |
| WBR | FRMedieval | Orcadian | 0.6743 | 0.337 | 0.663 | 0.100 |
| WBR | FRMedieval | Vikings | 0.4740 | 0.338 | 0.662 | 0.386 |
| WBR | FRMedieval | Scottish | 0.9438 | 0.386 | 0.614 | 0.090 |
| WBR | FRMedieval | Icelandic | 0.7598 | 0.414 | 0.586 | 0.082 |
| WBR | FRMedieval | England_Saxon.SG | 0.4363 | 0.468 | 0.532 | 0.125 |
| WBR | FRMedieval | Iceland_Pre_Christian.SG | 0.2734 | 0.551 | 0.449 | 0.114 |
| WBR | FRMedieval | Ireland_BA.SG | 0.6912 | 0.569 | 0.431 | 0.103 |
| EBP | FRMedieval | - | 0.3551 | - | - |  |
| EBP | FRMedieval | Hungarian | 0.7005 | 0.369 | 0.631 | 0.169 |
| EBP | FRMedieval | Germany_EMedieval.SG | 0.7579 | 0.392 | 0.608 | 0.171 |
| EBP | FRMedieval | England_Roman.SG | 0.7827 | 0.397 | 0.603 | 0.303 |
| EBP | FRMedieval | English | 0.5775 | 0.437 | 0.563 | 0.216 |
| EBP | FRMedieval | Vikings | 0.8665 | 0.480 | 0.520 | 0.115 |
| EBP | FRMedieval | Norwegian | 0.9295 | 0.514 | 0.486 | 0.100 |
| EBP | FRMedieval | Orcadian | 0.8039 | 0.531 | 0.469 | 0.114 |
| EBP | FRMedieval | Scottish | 0.9340 | 0.548 | 0.452 | 0.104 |
| EBP | FRMedieval | Icelandic | 0.8634 | 0.561 | 0.439 | 0.100 |
| EBP | FRMedieval | Iceland_Pre_Christian.SG | 0.6714 | 0.651 | 0.349 | 0.114 |
| EBP | FRMedieval | England_Saxon.SG | 0.6094 | 0.660 | 0.340 | 0.130 |
| EBP | FRMedieval | Ireland_BA.SG | 0.7875 | 0.695 | 0.305 | 0.106 |
| NOR | FRMedieval | - | 0.1628 | - | - |  |
| NOR | FRMedieval | England_Roman.SG | 0.6352 | 0.236 | 0.764 | 0.276 |
| NOR | FRMedieval | Hungarian | 0.2784 | 0.295 | 0.705 | 0.300 |
| NOR | FRMedieval | Germany_EMedieval.SG | 0.2726 | 0.404 | 0.596 | 0.295 |
| NOR | FRMedieval | Vikings | 0.4790 | 0.459 | 0.541 | 0.140 |
| NOR | FRMedieval | Norwegian | 0.4225 | 0.551 | 0.449 | 0.133 |
| NOR | FRMedieval | Scottish | 0.5091 | 0.563 | 0.437 | 0.130 |
| NOR | FRMedieval | Orcadian | 0.3055 | 0.575 | 0.425 | 0.160 |
| NOR | FRMedieval | Icelandic | 0.3413 | 0.612 | 0.388 | 0.130 |
| NOR | FRMedieval | Iceland_Pre_Christian.SG | 0.3397 | 0.640 | 0.360 | 0.130 |
| NOR | FRMedieval | Ireland_BA.SG | 0.4111 | 0.696 | 0.304 | 0.125 |
| NOR | FRMedieval | English | 0.1308 | 0.746 | 0.254 | 0.701 |
| NOR | FRMedieval | England_Saxon.SG | 0.1839 | 0.748 | 0.252 | 0.190 |
| HAU | FRMedieval | - | 0.5685 | - | - |  |
| HAU | FRMedieval | English | 0.8013 | 0.479 | 0.521 | 0.193 |
| HAU | FRMedieval | England_Roman.SG | 0.9115 | 0.486 | 0.514 | 0.204 |
| HAU | FRMedieval | Hungarian | 0.7682 | 0.515 | 0.485 | 0.169 |
| HAU | FRMedieval | Norwegian | 0.9407 | 0.589 | 0.411 | 0.107 |
| HAU | FRMedieval | Vikings | 0.8224 | 0.605 | 0.395 | 0.134 |
| HAU | FRMedieval | Germany_EMedieval.SG | 0.6659 | 0.616 | 0.384 | 0.247 |
| HAU | FRMedieval | Orcadian | 0.8163 | 0.630 | 0.370 | 0.131 |
| HAU | FRMedieval | Icelandic | 0.8971 | 0.631 | 0.369 | 0.108 |
| HAU | FRMedieval | Scottish | 0.8971 | 0.638 | 0.362 | 0.120 |
| HAU | FRMedieval | Iceland_Pre_Christian.SG | 0.8094 | 0.694 | 0.306 | 0.114 |
| HAU | FRMedieval | England_Saxon.SG | 0.7117 | 0.729 | 0.271 | 0.141 |
| HAU | FRMedieval | Ireland_BA.SG | 0.8836 | 0.734 | 0.266 | 0.104 |

**Table 1** (continued)
| Target | Source 1 | Source 2 | p-value | Prop 1 | Prop 2 | std error |
| --- | --- | --- | --- | --- | --- | --- |
| GRA | FRMedieval | - | 0.7582 | - | - |  |
| GRA | FRMedieval | Hungarian | 0.8134 | 0.680 | 0.320 | 0.188 |
| GRA | FRMedieval | Germany_EMedieval.SG | 0.7388 | 0.812 | 0.188 | 0.235 |
| GRA | FRMedieval | Vikings | 0.7452 | 0.831 | 0.169 | 0.175 |
| GRA | FRMedieval | Norwegian | 0.7569 | 0.846 | 0.154 | 0.150 |
| GRA | FRMedieval | Scottish | 0.7496 | 0.866 | 0.134 | 0.148 |
| GRA | FRMedieval | Icelandic | 0.7235 | 0.889 | 0.111 | 0.155 |
| GRA | FRMedieval | Orcadian | 0.7141 | 0.897 | 0.103 | 0.179 |
| GRA | FRMedieval | Ireland_BA.SG | 0.7502 | 0.901 | 0.099 | 0.116 |
| GRA | FRMedieval | England_Roman.SG | 0.7331 | 0.905 | 0.095 | 0.286 |
| GRA | FRMedieval | Iceland_Pre_Christian.SG | 0.7180 | 0.921 | 0.079 | 0.153 |
| GRA | FRMedieval | English | 0.6927 | 0.961 | 0.039 | 0.287 |
| GRA | FRMedieval | Iberia_Medieval_published | 0.7869 | 0.988 | 0.012 | 0.307 |
| GRA | FRMedieval | England_Saxon.SG | 0.6917 | 0.989 | 0.011 | 0.176 |
| CEN | FRMedieval | - | 0.5725 | - | - |  |
| CEN | FRMedieval | Hungarian | 0.5170 | 0.825 | 0.175 | 0.266 |
| CEN | FRMedieval | Vikings | 0.5265 | 0.855 | 0.145 | 0.199 |
| CEN | FRMedieval | Norwegian | 0.5286 | 0.884 | 0.116 | 0.163 |
| CEN | FRMedieval | Scottish | 0.5180 | 0.912 | 0.088 | 0.162 |
| CEN | FRMedieval | Germany_EMedieval.SG | 0.5012 | 0.917 | 0.083 | 0.334 |
| CEN | FRMedieval | Ireland_BA.SG | 0.5366 | 0.917 | 0.083 | 0.130 |
| CEN | FRMedieval | Icelandic | 0.4998 | 0.940 | 0.060 | 0.168 |
| CEN | FRMedieval | England_Roman.SG | 0.5335 | 0.941 | 0.059 | 0.359 |
| CEN | FRMedieval | Iceland_Pre_Christian.SG | 0.5143 | 0.943 | 0.057 | 0.160 |
| CEN | FRMedieval | Orcadian | 0.4926 | 0.973 | 0.027 | 0.200 |
| CEN | FRMedieval | Iberia_Medieval_published | 0.7578 | 0.997 | 0.003 | 0.296 |
| SLO | FRMedieval | - | 0.7921 | - | - |  |
| SLO | FRMedieval | Hungarian | 0.8100 | 0.734 | 0.266 | 0.201 |
| SLO | FRMedieval | Vikings | 0.7962 | 0.814 | 0.186 | 0.171 |
| SLO | FRMedieval | Germany_EMedieval.SG | 0.7660 | 0.834 | 0.166 | 0.240 |
| SLO | FRMedieval | Norwegian | 0.7981 | 0.841 | 0.159 | 0.147 |
| SLO | FRMedieval | Spanish_North | 0.7422 | 0.845 | 0.155 | 0.547 |
| SLO | FRMedieval | England_Roman.SG | 0.7864 | 0.857 | 0.143 | 0.268 |
| SLO | FRMedieval | Scottish | 0.7816 | 0.874 | 0.126 | 0.146 |
| SLO | FRMedieval | Icelandic | 0.7708 | 0.879 | 0.121 | 0.148 |
| SLO | FRMedieval | English | 0.7452 | 0.882 | 0.118 | 0.263 |
| SLO | FRMedieval | Orcadian | 0.7568 | 0.887 | 0.113 | 0.171 |
| SLO | FRMedieval | Ireland_BA.SG | 0.7769 | 0.910 | 0.090 | 0.119 |
| SLO | FRMedieval | Iceland_Pre_Christian.SG | 0.7520 | 0.922 | 0.078 | 0.146 |
| SLO | FRMedieval | Iberia_Celtiberian | 0.6461 | 0.958 | 0.042 | 0.287 |
| SLO | FRMedieval | England_Saxon.SG | 0.7308 | 0.974 | 0.026 | 0.168 |
| SLO | FRMedieval | Iberia_Medieval_published | 0.6612 | 0.986 | 0.014 | 0.355 |
| SLO | FRMedieval | Basque | 0.7237 | 0.993 | 0.007 | 0.355 |
| NOU | FRMedieval | - | 0.7250 | - | - |  |
| NOU | FRMedieval | Spanish_North | 0.9240 | 0.311 | 0.689 | 0.304 |
| NOU | FRMedieval | Basque | 0.7515 | 0.642 | 0.358 | 0.263 |
| NOU | FRMedieval | Iberia_IA | 0.6964 | 0.863 | 0.137 | 0.254 |
| NOU | FRMedieval | Iberia_Celtiberian | 0.6659 | 0.865 | 0.135 | 0.254 |
| NOU | FRMedieval | Iberia_Medieval_published | 0.7030 | 0.914 | 0.086 | 0.287 |
| NOU | FRMedieval | Sardinian | 0.6689 | 0.957 | 0.043 | 0.088 |

We also tested two-way models combining Mediaeval French with other ancient or present-day (non-French) populations (Table 1). We found that models fitting the WBR ancestry involve populations from the British Isles, Norway or Iceland (0.27< *P* <0.96). Indeed, the proportion of ancestry derived from present-day English or English Romans as proxy sources exceeds 90% (standard error of 0.26 and 0.23, respectively) versus less than 10% from Mediaeval French. Moreover, we observed that two-way models involving Northwestern Europeans/Scandinavians and Mediaeval French fit the ancestry of the other present-day French (EBP, NOR and HAU) on the English Channel coast (*P* > 0.13, Table 1). However, the Scandinavian or Northwestern European contribution in these groups is smaller than that found in WBR. For EBP, NOR, HAU, GRA, CEN and SLO, we noted that two-way models including a central European proxy source - such as early Mediaeval individuals from Germany or modern Hungarians - also fit the data with estimated contributions as large as 0.70 and 0.63 to NOR and EBP, respectively. However, they do not represent a good fit for the ancestry of WBR or NOU, in agreement with the low proportions of Germany-related ancestry previously observed with the supervised clustering analysis (Fig. 3b). In sum, *qpAdm* modelling shows relative population continuity between the Mediaeval period and present-day times in Northwestern France while emphasising a north/south Loire split with respect to the genetic contribution from Northwestern or Mediterranean Europe, respectively. Although genetic ancestry from countries in Northwestern European is found across the multiple populations dwelling north to the river Loire, WBR is the population in which such ancestry is found at its maximum.

### Large genetic affinities between Brittany and ancient Bell Beakers

Using a three-way admixture model, we then estimated the ancestry contributions from the three major ancient populations that spread across Europe - western hunter-gatherers (WHG), early farmers (EF) and steppe pastoralists (SP) – to present-day French and confirmed the aforementioned results (Fig. 4b and Table S2.4). SP ancestry proportion for regions bordering the Channel (HAU, NOR, EBP and WBR) varies from 42% to 46% versus less than 40% among CEN, SLO, NOU and GRA, and reaches its maximum in WBR where it is similar, if not larger than EF ancestry (46% +- 2.1% and 43.6% +- 2%, respectively, Table S2.4). Consistently, significant negative values (Z-score < -4, Table S2.5) for *f4* statistics of the form *f4(Mbuti, Russia_EBA_Yamnaya_Samara; WBR, other modern French)* revealed increased allele sharing between Early Bronze Age Yamnaya pastoralists from the Eurasian steppe and WBR relative to other present-day French samples. In agreement with these elevated levels of steppe-like ancestry found in WBR, the results of the outgroup *f3*-statistics of the form *f3*(Mbuti; Bell Beaker, present-day Europeans) and the *f4*-statistics of the form *f4*(*Mbuti, Bell Beaker; WBR, other modern French*) show that Bell Beaker-associated individuals from Northwestern Europe – who are reported to carry large amounts of steppe-related ancestry - share large levels of drift and significantly increased allele sharing (Z-score < -3) with WBR relatively to other present-day French samples (Fig. 4c and Fig. S4.2). *f4* statistics of the form *f4(Mbuti, Western Hunter-gatherer; WBR, other modern French)* were mostly not significant (except when the other present-day French was either NOU or GRA, see Table S2.5), which allow us to exclude the possibility that the genetic affinities between WBR and ancient Bell-Beaker-associated individuals could be caused by a significantly increased WHG ancestry uniquely in WBR.

Altogether, these results indicate that the largest contribution of steppe-related ancestry in present-day France is found across the regions located north to the Loire River, and that WBR shares similar levels of steppe ancestry only with other populations living along the northwestern shores of the European continent (e.g., Ireland, Scotland, Orkney Islands, Iceland and Norway).

## Discussion

By exploring the whole genomes of present-day individuals originating from Northwestern France in comparison to those from neighbouring French and European populations, we provide novel insights into the demographic events shaping the genetic makeup of the Northwestern edges of the European landmass.

First, we found that population structure in the northern part of France is accompanied with variation in genetic affinities with Northwestern versus Southwestern Europeans. In particular, observations based on supervised admixture analysis, *f3*-outgroup statistics and pairwise F_ST_ values consistently attest connections between Western Brittany, and Western Great Britain (*e.g.*, Wales and Cornwall) and Ireland (Fig. 3). These findings confirm previous reports of strong genetic affinities between Brittany and Ireland relative to the general population of Great Britain (44). They were further supported by haplotype-based methods, which are more powerful than allele frequency-based methods to capture differential contributions from closely related populations (30,51). Indeed, by inferring ancestry profiles from a set of surrogate sources with GLOBETROTTER, we highlighted the ancestry contribution from Irish populations into present-day Bretons (∼24%) and found relatively smaller contributions from Wales or Cornwall (∼9- and ∼3-fold, respectively). Importantly, we detected some Irish ancestry across all the French regions we surveyed, leading us to hypothesise a long history of shared ancestry likely on the basis of the genetic makeup of the Celtic (Iron-Age) Gaul. The ancestry contributions of Wales and Cornwall exclusively found in Brittany, albeit very limited, are likely associated with more recent migrations. The importance of ancient migrations between the British Isles and Northwestern France has been previously proposed based on ubiquitous haplotype sharing between samples from France and the British Isles, especially samples from Western Great Britain and Northern Ireland(5). Such results fit with the reported higher frequencies of mutations associated with cystic fibrosis, hemochromatosis and lactase persistence shared between Brittany and Ireland(44,52,53). In parallel, our observations reveal larger genetic affinities between populations from Northern France and present-day individuals from Germany (Fig. 3). As for present-day French populations located south to the river Loire, genetic proximity is observed with contemporary Spanish, and signals of shared ancestry with the Basques. Basque people have recently been described as a typical Iberian Iron Age population whose ancestry was not influenced by later admixture into Iberia(54). These results are compatible with a scenario where Atlantic France south of the Loire shares the Iron Age legacy of the Basques, while it diverges from the Basques likely due to higher levels of gene flow associated with later incoming migrations (*e.g.*, Germanic invasions) or simply through isolation by distance from other regions to the north and to the east.

Considering the relative ancestry contributions from ancient populations prior to 2000 BCE that spread across Europe, we provided evidence for differential distribution of steppe vs. Early Neolithic ancestry between the two sides of the Loire River, with Neolithic ancestry being more prevalent south of the river. These findings likely explain the shared ancestry between present-day Atlantic French located south of the Loire River (such as *Nouvelle-Aquitaine*) and the Neolithic-enriched Basque population. Similarly, the relatively high proportions of steppe ancestry found north to the Loire River and along the coast of the English Channel (Fig. 4b and Table S2.5) - and in particular in Brittany - might explain the genetic affinities observed with Ireland and the Western Great Britain (Fig. 3). Present-day Irish exhibit a strong signal of continuity with the geographically close Early Bronze Age individuals carrying high levels of steppe ancestry (∼39 ± 8%) and among whom was found the earliest presence of the hemochromatosis mutation(55). In Ireland and in the rest of the British Isles, the introduction of steppe ancestry, which led to a considerable turnover of the contemporaneous genetic makeup, has been recently linked to the migration of individuals associated with the Bell Beaker complex from Northern/Central Europe(18,55). The lack of human ancient DNA from northern France - dating from the period between Copper Age and Early Iron Age - has hampered our understanding of such genetic turnover across the northern coast of France. Given (i) the elevated levels of steppe ancestry found among present-day French located along the English Channel coast and (ii) the high degree of allele sharing observed between present-day individuals from Brittany and Bell-Beakers-associated individuals carrying high levels of steppe ancestry (Fig. 4), we hypothesise that a similar degree of turnover has reached northern France and Brittany after the arrival of steppe ancestry. Previously reported Late Iron Age individuals from the region of Normandy were found to carry considerable levels of mitochondrial steppe ancestry and strong affinities with Bronze Age samples from the British Isles(19), suggesting extensive gene flow across the English Channel during the Bronze Age. This hypothesis is supported by metalwork-based archaeological evidence dating from the Middle Bronze Age, which points to the existence of a British Channel metalworking core area(56). Indeed, metal-based relationships between Ireland and Brittany may have started earlier. In Ireland, Bell Beaker pottery is closely associated with the introduction of copper metallurgy ∼2400 BC. However, while Irish Bell Beaker culture shows strong influence from Britain, the introduction of metalwork appears to have occurred through Atlantic Europe(17). Metalwork identified in Brittany from the Chalcolithic and Early Bronze Age suggests exchanges of metals such as copper and tin between the regions of Brittany, Ireland(57) and Cornwall(17). Archaeological evidence exists for extensive connections between Brittany and Ireland/Western Britain dating as early as 4500 BCE and related to the phenomenon of Megalithic monument construction. However, genetic data presented in this study do not support a scenario where the close relationship between Brittany and Ireland/Western Britain can be simply explained by a larger sharing of alleles pre-dating the arrival of the steppe ancestry. Samples associated with the Megalithic period from the British Isles exhibit similar levels of allele sharing with all the present-day samples from France (Table S2.6), in contrast with increased allele sharing with Corded-Ware- and steppe-enriched Bell Beakers-associated samples uniquely found in Brittany (Fig. S4.2 and Table S2.7). Furthermore, we found no differences in genetic affinities between present-day French and the ancient samples related to the two Neolithic waves - the Mediterranean and the Danube waves - that could alternatively explain both the differences between north and south of the river Loire and the close relationship between Brittany and Ireland (Table S2.8).

In contrast with the pattern found for the distribution of Neolithic and steppe ancestries, we found that fractions of western hunter gatherer (WHG) ancestry do not differ across the two margins of the river but instead tend to decrease inland (Fig. 4b, Table S2.4). Larger WHG proportions in present-day coastal French populations might reflect a longer coexistence between late Mesolithic populations and newly incoming farmers due to the significant exploitation of marine resources by the former communities(9), as it seems to have been the case in Brittany. However, a denser sampling of present-day populations from central and eastern France is required to test this hypothesis. Together our results point to a scenario where the migration of people associated with the Bell Beaker complex culture from North/Central Europe considerably influenced the genetic makeup of Northwestern France during the Bronze Age by introducing steppe-related ancestry all along the French coast of the English Channel. However, whether this occurred through direct influence of the Bell Beaker incomers from the east or through extensive contacts with the British Isles is an important question that can only be addressed through the study of human ancient DNA from Northern France from the Copper to the Early Iron Age. Importantly, Brittany is the place where the genetic legacy associated with the introduction of steppe ancestry is currently the strongest. This likely indicates relative isolation from later continental migrations, which seem to have increased Neolithic ancestry eastwards. Our admixture modelling approach lends support to this view by showing a lack of genetic affinities between Central European and Brittany, as one- or two-way models involving such populations (e.g. Hungarians, Germans) do not fit the data for Western Brittany (Table 1). This contrasts with the results for all the remaining present-day French populations. The introduction of this Central European component, probably less rich in steppe ancestry, might reflect the influence of Germanic peoples in present-day France during the first millennium CE similarly to what has recently been reported in England(58). Nevertheless, we do not exclude the possibility that the shift towards a Central European component, which we observed across multiple analyses with the “merged-modern dataset” (Fig. 3), could be explained by earlier migrations.

We found Mediaeval samples from Western France to carry generally less steppe ancestry than their geographical close present-day populations (Fig. 4b). This is consistent with a model in which the introduction of steppe ancestry in Northern French ∼2000 BCE remained restricted to the coastal or near coastal regions for centuries. Nevertheless, we recall that Early Mediaeval samples display substantial genetic heterogeneity as two samples carry contrasting proportions of steppe- vs. Neolithic-ancestry (fra016 and fra017) and one sample (fra009) did not fit the genetic diversity of present-day France. Instead, this sample seems to originate from North Africa and provides evidence for long-distance migration between the northern part of France and northern Africa, as early as the Early Mediaeval period (∼5-6 century CE). Finally, we found a lack of genetic continuity between Mediaeval French and Iberian populations dating from the first millennium BCE (*P* < 0.05, results not shown). Signals of genetic continuity were only found with Iberian individuals archaeologically associated with Germanic invasions, suggesting that until late Antiquity and Early Mediaeval Period (3th-10th century CE) French and Iberians might have kept low levels of gene flow.

Focusing on the genetic structure of the present-day populations from Northwestern France, global clustering patterns based on fineSTRUCTURE overlap with the distribution of surnames as well as the linguistic boundaries between varieties of Breton, which are spoken in the westernmost part of the Peninsula of Brittany, and Gallo-Romance varieties, spoken across the eastern part of the peninsula and the whole neighbouring region of *Pays-de-la-Loire* (Fig. 1). The role of linguistic relationships in the worldwide patterns of structure of human populations has long been recognized(59). Within-country studies, such as the one we present here, have not only validated previous claims but also showed that such a relationship holds at local geographical scales, as it has recently been shown in Northern Spain and in Great Britain(3,5). While we found that Western Brittany forms a distinct cluster based on haplotype similarity and rare allele sharing, it is worth noting that genetic differentiation increases from the westernmost tip of the peninsula of Brittany southwards, and especially south of the Loire River. The differentiation between Western Brittany and regions south of the Loire River is larger than that found along the west-east axis (between the *département* of *Finistère* and that of *Sarthe*) north of the Loire River, despite similar geographic distances (∼350 km). The role of the Loire River as a geographical barrier to the movement of people has been recently proposed due to the overlap between genetic cluster boundaries, inferred across the whole country, and the natural occurrence of the river (31). In the Netherlands, rivers have also been shown to locally restrict gene flow(60), but the general role of water bodies in promoting or impeding human movements remains unclear. A good understanding of the role of the Loire River on the local patterns of migration requires associating the river’s features with genetic differentiation measures along its full extension, which, in our case, is not possible due to the absence of samples along its upstream drainage area. Altogether, the structure patterns observed within Northwestern France contrast with the simple vision that genetic differentiation increases with the geographical distance(43).

At finer scales, present-day data revealed extensive population structure across Brittany and *Pays-de-la-Loire*, despite the overall low levels of differentiation (F_ST_ = 0.00043). Within Brittany, at intermediate resolution levels, clustering patterns largely overlap with watercourses (see SOM for further details). Furthermore, low population differentiation and increased IBD sharing between the cluster “Vannes” and other clusters eastwards on one side, and between the westernmost clusters “Léon” and “Cornouaille” on the other side, suggest a dichotomy between northwest and southeast. Such division is in agreement with the accruing geo- linguistic evidence for a bipartition of Breton varieties into two groups - the Breton varieties spoken in the northwest and those spoken in the southeast of the peninsula of Brittany, whose distributions coincide with the ranges occupied by the Gaulish (Celtic) Ossimii and Veneti, respectively(61). Noteworthy, the cluster “Bretagne-centre” roughly overlaps with the border of the Ossimii and Veneti territories, suggesting relative genetic homogeneity likely due to extensive gene flow at this border. Such homogeneity is also recognizable at the level of linguistic features (Fig. S.1.11, Fig. S1.12, Fig. S5; see SOM for details). Finally, we also found genetic clusters with no apparent geographical barrier at their border, such as “Vannes” and “Guérande”. Overall, the genetic structure observed within Brittany appears consistent with a complex scenario of interaction between geographical, cultural and likely economic factors influencing population connectivity. Located south to the Loire River, the region of *Mauges -* which exhibits the highest density of clusters - is situated at the southeastern tip of the Armorican Massif and encompasses hedged farmlands crossed by steep valleys. We hypothesise that this landscape contributed to relative isolation and increased genetic drift and lasted until about 300 years ago, as estimated in the dataset (assuming a generation time of 29 years/generation, Fig. S1.15). Such a scenario could explain the increase in haplotype sharing for long, and likely recent, chromosomal segments (Fig. S1.9). Other population-specific features, such as recent ancestry caused by inbreeding, could equally explain such strong clustering patterns. However, levels of relatedness were not found significantly larger than in other clusters (Fig. S1.2-S1.4). This ultra-fine-scale structure with geographically restricted clusters resembles that observed in the region of Galicia, Spain(3) and emphasises the need for whole-genome sequencing studies on local populations in order to better understand the distribution of rare, and likely more deleterious, variants across space(1).

In conclusion, the patterns of genetic structure observed across Brittany and the downstream Loire basin mainly reflect the likely combined effect of linguistics and geographic features. This lends further support to the idea that local population differentiation exists, as it has been shown within other countries (3,5), and can be detected at geographical distances as small as a few tens of kilometres even in the absence of major geographical barriers. Within this structured genetic landscape, Brittany reveals a history of isolation from the rest of France together with a strong legacy of the important genetic changes (i.e. the introduction of steppe ancestry) that followed the arrival, in Northwestern Europe, of people associated with the Bell Beaker complex from north-central Europe ∼2500 BCE. A similar scenario has been proposed for the present-day Celtic populations in the western part of the British Isles - the Welsh and Irish - who display elevated haplotype sharing with Bronze Age samples(55) and strong genetic affinities with pre-Anglo-Saxon samples from Britain(62). Our results imply that: 1) a scenario in which Mediaeval population movements along the English Channel and the Atlantic façade, such as those related to incoming Western Britons in Brittany during the 4-6th century CE or the Viking incursions ∼8-9th century CE, are unlikely to be the main explanation for the close genetic between the northwestern edges of the European continent; and 2) a long-history of shared ancestry between Brittany and Ireland/Western Britain followed by their relative isolation explains the sharing of disease-related alleles such as those associated with hemochromatosis, cystic fibrosis and lactase persistence.

## Methods

### France administrative division overview

This study relies on a dense sampling of the geographical region of western France, which crosses multiple administrative subdivisions of the territory. Among these subdivisions are the 1) regions - the largest administrative units - which are subdivided into 2) *départements.* A *département* is subdivided into 3) *arrondissements*, which at the smallest scale are divided into 4) town centers. Given the specificity of the administrative system we kept the French name for the *département* and *arrondissement* along the manuscript.

### Cohort description

Project PREGO (“Population de référence du Grand Ouest”, www.vacarme-project.org) collected the DNA of 5,707 healthy persons originating from western France (*Pays de la Loire* and Brittany regions). Individuals were recruited during 295 blood drives organised by the French Blood Service (EFS in French) carried out between February 2014 and March 2017, with a mean of 19 donors per blood drive. Blood drives were spatially and temporally sampled in order to obtain a coverage as homogeneous as possible of the nine *départements* included in the study. Priority was given to blood drives taking place in rural areas. Participants should be native of western France. Individuals’ birthplace was assessed by the place of birth of the four grandparents. Only individuals whose four grandparents were born in western France and preferably within a radius of 30 km were included in this study. From the 3,234 individuals included in the present study, 25%, 50% and 75% have their grandparents born 3.25, 6.38 and 12.33 kms from each other, respectively.

Venous blood samples (6ml) were collected from recruited individuals by venipuncture into Vacutainer tubes. Participants filled out a questionnaire providing grandparents’, parents’ and their own birthplaces, residence, age, sex and information about previous participation in the study (of the individual itself or another member of the family). Neither phenotypic nor clinical data was collected in the present study. Declaration and ethical approval process was achieved in 2013 and involved the Ministry of Research, the Committee of protection of persons (*Comité de Protection des Personnes*, CPP in French), the Advisory Committee on Information Processing for Health Research (CC- TIRS in French), and the National Commission on Informatics and Liberty (CNIL in French). Participants signed a written informed consent for participation in the study, inclusion in bio-resource and personal data processing.

The FranceGenRef study aims to describe patterns of population diversity across metropolitan France at the beginning of the 20th century. Thus, individuals were sampled based on the birthplace of their grandparents, whose distance should not exceed 30 kilometres. FranceGenRef includes a total of 862 individuals satisfying the aforementioned criteria and sampled under the scope of three different studies: 50 blood donors sampled in the *département* of *Finistère* (FIN, Fig. 1b), 354 blood donors from the PREGO cohort (www.vacarme-project.org) with origin in the three other *départements* of the region of Brittany - *Côtes d’Armor* (COT), *Ille-et-Vilaine* (ILL) and *Morbihan* (MOR) - and in the five *départements* of the region of *Pays-de-la-Loire* - *Loire-Atlantique* (LOI), *Maine-et-Loire* (MAI), *Mayenne* (MAY), *Sarthe* (SAR) and *Vendée* (VEN) -, and finally 458 individuals from the GAZEL cohort (www.gazel.inserm.fr/en)(63,64), among which are individuals from five other regions of France: Normandie, Hauts-de-France, Grand East, Centre-Val de Loire and Nouvelle-Aquitaine. All individuals signed informed consent for genetic studies at the time they were enrolled for the blood collection. DNA samples from GAZEL samples were extracted at the CEPH Biobank on an automated system either Autopure (Qiagen) or Chemagic Prime (PerkinElmer) using respectively the salting out method or magnetic beads and were quantified using fluorimetry (Quant-iT DNA Assay kit, Broad Range, Thermo Fisher Scientific).

### Genotyping, whole-genome sequencing and quality control (QC)

#### SNP array genotype processing

Under the scope of PREGO, out of 5,707 collected samples, 3,385 were genotyped on the Axiom TM Precision Medicine Research Array (∼ 920,000 markers, ThermoFisher). Standard QC was performed using SNPolisher software (http://tools.thermofisher.com) and SNPs not passing the QC report were removed according to the manufacturer’s instructions. SNPs with a missing rate >5%, minor allele frequency <10% and not in Hardy-Weinberg Equilibrium (*p<10^-6^*) were excluded from the dataset resulting in a total of 210,171 SNPs. Relatedness was assessed via the PI_HAT statistics, which provides an estimation of the proportion of the genome shared by two pairs of individuals (*i.e.* identity-by-descent, IBD). PI_HAT statistics was computed using PLINK (vs.1.9). After removing related individuals with a PI_HAT >= 0.08, there were 3,234 samples left for analysis. Genotyping was conducted in three batches of 971, 1266, 997 individuals, respectively. To investigate potential batch effects, we employed linear regression using a batch indicator variable on the first five principal components. We found no significant association at the significance level of 0.05.

#### Whole-genome sequencing and variant calling

Whole-genome sequencing of 856 individuals was performed at CNRGH (Evry, France) using their standard workflow (www.cnrgh.fr). All the samples were sequenced at high coverage (average coverage 30x). Read processing was performed with GATK 3.8 following the “best practices” NGS pipeline recommendations of the Broad Institute (https://software.broadinstitute.org/gatk/best-practices). Reads were mapped on the GRCh37 reference genome using *bwa-mem,* duplicates were removed and reads were then realigned and recalibrated according to GATK best practices. Variant calling was performed with GenotypeGVCF. GVCF files were then merged using GATK CombineGVCF function, recalibrated and annotated with SnpEff (65) and the gnomAD database (https://gnomad.broadinstitute.org/). The resulting GVCF files were filtered out in order to only keep SNPs with high mapping quality (MQ>30). Furthermore, only SNPs in Hardy-Weinberg equilibrium (HWE, *p-value* = 10^-5^) and less then 10% of missing data were kept for analysis.

### Merging with publicly available datasets

#### Publicly available datasets of western Europeans

To investigate the relationship of modern individuals from the north part of France and other European populations we merged the WGS dataset with three available genome-wide datasets encompassing a large number of western European samples: 1) the EGAD00000000120 from the The International Multiple Sclerosis Genetics Consortium and the Wellcome Trust Case Control Consortium 2 (referred to hereafter as the MS dataset)(42), 2) the EGAD00010000124 from the Genetic Analysis of Psoriasis Consortium & the Wellcome Trust Case Control Consortium 2 (referred to hereafter as PS dataset)(41), and 3) the EGAD00010000632 from the Peopling of the British Isles (referred to hereafter as POBI,(5)). In the MS and PS datasets samples were genotyped on the Human670- QuadCustom SNPchip encompassing 580,030 autosomal sites and the datasets include 11,376 and 2,622 individuals, respectively. MS dataset includes samples from: Australia, Belgium, Denmark, Germany, Finland, France, Italy, New Zealand, Northern Ireland, Norway, Poland, Spain, Sweden, US and UK, whereas the PS dataset includes samples from UK and Ireland. In the POBI dataset, 2,912 individuals from the UK were genotyped on the Human1-2M-DuoCustom SNPchip encompassing 1,115,428 autosomal sites. For the three datasets, original genotype likelihood files (.gen) were converted into plink format files with gtool (vs. 0.7.5). Only individuals and sites passing quality criteria thresholds (provided with the datasets) in the original studies were kept. Genotypes were called using a probability cut-off of 0.90. Sites containing alleles in the negative strand were flipped according to the corresponding strand file https://www.well.ox.ac.uk/~wrayner/strand/ using PLINK vs.1.9. First, we checked whether the alleles were in the illumina TOP configuration, as required to flip based on the strand file. Sites not found in TOP configuration were removed from the datasets. We used the liftOver tool (https://genome.ucsc.edu/cgi-bin/hgLiftOver) to convert the physical coordinates to hg19 as they were originally in hg18 in the MS, PS and POBI datasets. We merged the samples from Belgium, Denmark, Germany, Finland, France, Italy, Northern Ireland, Norway, Poland, Spain, Sweden and UK with the Irish samples from the PS dataset. In a second step, we merged this dataset with a subset (to keep the sample size computationally tractable) of the POBI dataset comprising two samples from Wales (Dyfed and Gwynedd), the sample from Cornwall and two samples from Norfolk and Kent (eastern UK). Finally, we merged the dataset above with seventy samples from the available Human Genome Diversity Project dataset (https://www.hagsc.org/hgdp/files.html) genotyped on the Illumina 650Y SNPchip array and originated from Sardinia, the Basque Country, Orkney Islands and the Mbuti (to use as an outgroup). Chromosomal coordinates for the HGDP dataset were lifted over as described above. From this merged dataset we filtered out sites not in Hardy-Weinberg Equilibrium (HWE, *p-value* = 10^-5^), removed those falling into the HLA region (chr6:29,691,116- 33,054,976) and removed all the multiallelic sites. The final dataset, hereafter referred to as “merged modern dataset”, encompassed 9704 samples and 433,940 SNPs.

### Human Origins Array and Viking datasets

We merged the French WGS dataset with the Human Origins Array (HOA) dataset V42.4 (https://reich.hms.harvard.edu/allen-ancient-dna-resource-aadr-downloadable-genotypes-present-day-and-ancient-dna-data) encompassing 3,589 ancient and 6,472 present-day individuals genotyped for 593,124 autosomal SNPs. From the original dataset we extracted European samples (Austria, Belgium, Czech_Republic, Denmark, France, Germany, Great_Britain, Greece, Hungary, Iceland, Ireland, Italy, Luxembourg, Netherlands, Norway, Poland, Portugal, Russia, Spain, Sweden, Switzerland, Turkey) with PASS tag under the “Assessment” field in the annotation file. From this subset we removed samples whose individuals’ ID contain *Ignore* and with less than 50,000 genotypes called to avoid potential bias. This dataset was then merged with 405 ancient DNA samples belonging to Vikings from the study: “Population Genomics of the Viking world”(49) and previously called for HOA set of SNPs. This dataset is referred to hereafter as “merged ancient dataset”. Mbuti samples (HGDP-CEPH, Human Genome Diversity Project-Centre d’Etude du polymorphisme Humain - https://cephb.fr/en/hgdp_panel.php) from the HOA were also extracted to compute *f*-statistics (see below).

#### ChromoPainter and fineSTRUCTURE

PREGO’s SNP-array dataset (post-QC) was phased with SHAPEIT v2.r790 (66) using the genetic map build 37 provided with the software and no reference panel. Phased genotype files were converted into CHROMOPAINTER(45) format using the impute2 chromopainter2.pl script. Switch (global Ne) and emission rates (μ) were estimated with CHROMOPAINTER version 2 using data on chromosomes 1, 4, 10 and 15 from 330 individuals (around 10% of the total sample). The estimated parameters were then used to run CHROMOPAINTER on the full data. A Principal Component Analysis (PCA) on the coancestry matrix (chunkcounts output) was performed in R. Coancestry matrix estimates the proportion of the genome of each individual that is most closely related to every other individual in the matrix. In particular, chunkcounts matrix is based on the number of copied haplotype chunks. We assigned individuals to clusters using the model-based approach implemented in fineSTRUCTURE version 2.1.3. We ran fineSTRUCTURE version 2.1.3 on the coancestry matrix with 10,000,000 burn-in iterations and 1,000,000 MCMC iterations, from which every 10,000th iteration was recorded. We kept the default values for the other options. The tree was built using 100,000 tree comparisons and 10,000,000 additional optimization steps. MCMC convergence was assessed by comparing the individual assignment to clusters in a second independent chain. Confidence measures of cluster assignment and visualisation of coancestry matrix were performed with the help of the R library FinestructureLibrary provided with the software. The tree provided by fineSTRUCTURE software is based on posterior probabilities of the population configuration (ie. individual partition across clusters), which is highly dependent on the sample sizes and does not reflect differences between pairs of clusters(32,67). Therefore, as an alternative we computed a tree based on the total variation distance (TVD) as firstly described by Kerminen and colleagues(32). TVD measures the distance between the clusters and it is not affected by differences in sample sizes. TVD values are computed from the copying profiles of all individuals, estimated with CHROMOPAINTER, and the cluster assignment inferred with fineSTRUCTURE as described in(32). It results in a final symmetrical matrix with as many rows and columns as the number of clusters inferred (K), where each entry represents the average closest ancestry contribution in cluster k coming from other clusters. We computed the TVD matrix on the final fineSTRUCTURE partition (k=<154, finest level of FS- tree) and the TVD-tree was obtained using complete linkage hierarchical clustering.

We assessed the performance of the two tree building approaches on the PREGO dataset. The performance was measured by comparing cluster assignment confidence as computed in Leslie *et al.*(5). As expected, cluster assignment confidence decreases with increasing number of clusters (Fig. S1.6.). Moreover, we found that for the same k the FS-tree shows lower cluster assignment confidence than the TVD-based tree, confirming the better performance of the latter. To visualise the relationship between the clusters (Fig. 1d), we arbitrarily chose *k*=39 because it provides a large enough number of clusters to access fine-scale structure while keeping cluster assignment confidence >90% (Fig. S1.6). At this level of the tree, clusters containing 1-5 individuals were merged into the closest cluster with >=31 individuals or removed if the closest cluster included itself <4 individuals. This approach resulted in a TVD- based tree of 18 clusters performing better than FS tree, which achieves similar values of cluster assignment confidence for only 12 clusters. Finally, we tested whether inferred clusters capture significant differences in ancestry by following the approach of (5). To do so, we randomly reassigned the individuals to clusters, maintaining the cluster sizes, and computed the p-value of the population configurations produced by the *k*=18. The TVD-based tree was less likely than a random distribution of individuals across pairs of clusters. We performed 1000 permutations to obtain p-values, which were computed as the number of permutations where random assignment resulted in a higher value of TVD between any pair of clusters and the total number of permutations. For the level of *k*=18, p-values were < 0.001 for all pairs of clusters. The tree shown in Fig. 1d was built using the clustree and ggtree packages of R-statistical software.

### Assessing sampling effect on fineSTRUCTURE clustering

We investigated whether the sampling scheme carried out in this study, which densely selects individuals representing every location over four generations, increases the probability of having recently related individuals. This recent ancestry may affect clustering patterns. To check the effect of recent ancestry on the fineSTRUCTURE clusters we computed a relatedness measure based on identity by state (IBS) within clusters, for *k*=3, 18 and 78 (only for those clusters with n > 10 at the finest scale level *k*=154), and compared it with the overall distribution. With the exception of *k*=154, where six clusters have a median IBS-statistics larger than the 75% of overall IBD distribution, across the other values of *k*, most of the clusters do not show increased relatedness (Fig S1.2-S1.4) and therefore, we conclude that recent ancestry is not driving the results.

### Surname analysis

The list of surnames associated with birth registration data and occurring in two periods (period one: 1891-1915 and period two: 1816-1940) were retrieved from the French Institute for Statistics and Economic Studies, the INSEE (https://insee.fr). Surname lists are available at the town level. To analyse the distribution of surnames we proceed as follows. 1) For each town, we chose the surnames present in the two periods and registered at least four times, *ie.* given to four newborns. Such an approach removes very rare names often associated with spelling errors in the registration and rare emigrants from other regions of France or elsewhere. 2) summed the number of surname occurrences over the two periods at the *arrondissement* level. There are 35 *arrondissements* within the ten selected *départments*. 3) We computed the Arccos distance (68) between the 35 *arrondissements* and 4) constructed a consensus tree based on 1000 bootstrapped distance matrices obtained by the Neighbour Joining method. Finally, we built a map linking the *arrondissements* in a nested way according to the bootstrap values of the NJ tree. The threshold for grouping *arrondissements* was 90% bootstrap robustness while the threshold for higher order grouping is 85%. There are various possible distances that can be computed from surname data (eg. Nei, Hedrick, Lasker, Euclidian). However, using different distances gives similar results (data not shown).

Correlation between surname based Arccos distance and F_ST_ was performed using Mantel distance matrix correlation test using Spearman rank correlation. The physical distance between arrondissements was done using partial correlation(69). Surname distribution diversity index were entropy and Barrai index. Let N be the population size in a geographical area and S the number of different surnames and pi the probability of Surname i.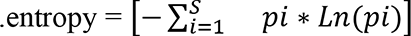

### PCA analysis and F_ST_ computation

PCAs was carried out using the smartpca software from the EIGENSOFT package version 6.1.4 (70). F_ST_ values were obtained by setting up the option F_ST_hiprecision: YES. Both analyses were performed after pruning for linkage disequilibrium using a sliding window of 50 SNPs in steps of 5 SNPs and keeping those SNPs with r^2^ < .50 (Fig. 3a and Fig. S2.2). The regions previously reported as exhibiting long-range linkage-disequilibrium (71) were excluded before performing the PCAs.

### Runs of homozygosity (ROH), Identity by descent (IBD) and IBDNe estimations

Individuals’ runs of homozygosity (ROH) and identity by descent (IBD) segments were calculated using RefinedIBD (version from 23rd December 2017) based on the PREGO dataset. Individuals’ ROH were computed by summing the total length in ROH segments per individual and averaged across individuals within *départments* and *arrondissements.* We computed IBD sharing by counting the number of IBD segments shared between pairs of individuals assigned to the 18 clusters inferred by fineSTRUCTURE. This was done independently for IBD segments of length: 1-2 cM, 2-7 cM and longer than 7 cM.

We estimated the trajectories of effective population size with IBDNe(33) (version released on 7th May 2018). IBD segments were identified with IBDseq (version r1206, with default settings). As suggested by Browning and Browning 2018 (72), we removed regions with excess of IBD to avoid potential bias, such as the Major Histocompatibility Complex (MHC) on chromosome 6 (chr6:26291527-33464061). We thus split chromosome 6 into two continuous parts. To evaluate the robustness of the effective population size trajectories across different IBD segment sizes, we varied the *mincm* parameter, which sets up the minimum length of IBD segments used by IBDNe. Value of *mincm* should be chosen considering SNP density on the SNP array.

### Rare variation analysis

We investigated allele sharing patterns using doubletons (alleles that are present in only two chromosomes or minor allele count MAC=2) and variants with MACs between 3 to 10 computed from the WGS dataset encompassing 620 individuals from Northwestern France. We first randomly selected 1 million sites from each dataset and allele sharing matrices were computed by summing all the variants shared between pairs of chromosomes of different individuals from Brittany and *Pays-de-la-Loire*. We plotted the results as a heatmap and performed hierarchical clustering (*hclust* function in R with method=”complete”) to identify clusters of highly correlated allele sharing. The resulting tree was cut at multiple levels (k=2- 10) and the proportion of individuals, within each *département*, assigned to the alternative clusters was plotted onto the map of Northwestern France. These analyses were performed using the R statistical package together with the following libraries: ComplexHeatmap, rgdal, sp, broom, ggplot2 and scatterpie.

### Supervised admixture analysis

The supervised clustering analysis was performed using ADMIXTURE vs1.3 software(73) on the WGS dataset including the 846 French samples used in the PCA. We assumed that modern French originate from three modern source populations: Spain, Germany and Ireland, which lie in the extremes of the distribution of modern French individuals in the PCA (Fig. 3a). Given that difference in sample sizes (see section *Publicly available datasets of western Europeans*) in the source populations we downsampled Germany and Ireland to 350 individuals. SNPs in linkage disequilibrium were removed by following the recommended practises in the software manual, ie. PLINK option *--indep-pairwise 50 10 0.1*. Only sites with minimum genotype rates <10% were analysed.

### Ancestry profiles with GLOBETROTTER

To further investigate the contribution of external populations to the genetic makeup of France we estimated ancestry contributions from their European neighbours using the statistical approach implemented in GLOBETROTTER software(45). This method identifies and describes recent (<4.5ky old) admixture events, by providing admixture times and proportions, explaining the haplotypic ancestry of a target population. Given that GLOBETROTTER infers admixture events from patterns of haplotype sharing it requires two input files: the “copy vectors” and “painting samples” generated by the CHROMOPAINTER. To obtain these files we ran CHROMOPAINTER following the author’s recommendations (section 8.2 “running ChromoPainterv2 with GLOBETROTTER”) on the phased “merged modern dataset”. The “copy vectors” files (*.chunklengths.out) were combined using the scripts provided with the software package. To detect signals of admixture we first ran GLOBETROTTER with the option *null.ind: 1* and then computed the number of bootstraps providing a date of admixture >= 1 and <= 400 generations. This step provides a *p-value* that can be interpreted as the evidence of “detectable admixture”. *P-values* were computed using 100 bootstraps. Confidence intervals for the inferred admixture times were estimated by running 100 additional bootstraps and with the option: *null.ind: 0*. We phased the “merged modern dataset” using SHAPEIT and no reference panel, in a similar manner to what was done with the PREGO dataset. To keep the analyses computationally tractable and due to imbalanced sample sizes, we randomly sampled 350 individuals from countries with large sample sizes such as Italy, Germany, Norway and Belgium. To represent the haplotypic diversity of the UK and the Scandinavian Peninsula we only kept the samples from PoBI and Norway, respectively. A total of 3,598 samples were used to estimate ancestry contributions.

### Admixture analysis: *f3-*, *f4*-statistics and *qpAdm* estimates

We computed the outgroup *f3-statistics* of the form *f3(Outgroup; Pop1, Pop2)*, as implemented in Admixtools(46), using the Mbuti population as an outgroup. Pop1 and Pop2 were either a pairwise combination of a modern French and any other European population or a Bell Beaker sample and a present-day western European. The outgroup *f3- statistics* quantifies the amount of shared history between pairs of populations in relation to an outgroup under a three-population tree. Provided that the outgroup is equally distant from Pop1 and Pop2, the outgroup *f3-statistics* represents the length of the branch from the outgroup to the internal node if no gene flow occurred between Pop1 and Pop2. To statistically assess the presence of admixture we computed *f4-statistics* using the same software package. Admixture was also computed in order to test whether modern and ancient Mediaeval French show signals of admixture *f3-statistics* by setting these populations as the *target.* Admixture *f3-statistics* was computed using the option *inbreed: YES* in the case of the ancient Mediaeval as recommended by the authors of the program.

*qpAdm*, implemented in the Admixtools, is a computer program that assesses the goodness of fit of admixture models, involving a user-defined set of commonly known as “left” and “right” populations, and estimates the proportion of admixture within a regression framework. The set of “left” populations encompasses the target population and the possible sources of admixture, whereas the “right” populations include a set of populations distantly related to the “left”. Importantly, the model assumes that the “right” populations are differentially related to the “left” and exchange of genes between “left” and “right” must not have occurred after the event of admixture, i.e. genetic drift shared between “left” and “right” derives from deep shared evolutionary history. The method implemented in *qpAdm* is built on a matrix of *f4-statistics* computed across all possible combinations of “left” and “right” populations (*f4*(target, source; R1, R2), where R1 and R2 is any possible pair of “right” populations). *P-values* > 0.05 together with admixture proportions varying between 0-1 are indicative of a good fitting model.

We performed the *qpAdm* analyses to estimate the contribution of the three major European migrations(50,74) using western hunter-gatherers (WHG), Neolithic early-farmers (EF) and Bronze Age steppe pastoralists (SP) as “left” populations together with the Mediaeval and modern French populations. We grouped ancient individuals into WHG, EF and SP based on the ancestry proportions inferred in Mathieson *et al.* (75). Only individuals with ancestry proportions >.90 were included into the three groups giving origin to sample sizes of 73, 42 and 17 for the EF, WHG and SP respectively. Admixture proportion estimates were computed using 25 randomly selected modern individuals for the three genetic clusters inferred by fineSTRUCTURE (WBR, EBP and SLO) and the remaining five regions of France (NOR, HAU, GRA, CEN and NOU). With respect to the French Mediaeval samples we inferred admixture proportions at the population (five Medieval French, the outlier was removed from this computation) and individual level. The set of *right* populations used in these analyses was the following: Mbuti, Han, AfontovaGora3, ElMiron, Karitiana, Kontenki14, MA1, Mota, Papuan, Ust_Ishim, Vestonice16, Villabruna, GoyetQ116-1, Morocco_Iberomaurusian.

In order to test for continuity of the genetic ancestry of Mediaeval and present-day French populations we tested one-way models. In the case of Mediaeval French, the single source is a neighbouring population (no post-Neolithic sample from France is available in the merged dataset) of the same period or of the preceding one. In the case of present-day French, the single source included the Mediaeval French and all the possible sources tested for the Mediaeval French. Furthermore, we also tested two-way models where the genetic ancestry of modern French is assumed to result from a mixture between the Mediaeval French and any other ancient population from the neighbouring regions from Mediaeval to present-day times. Given the larger availability of Iberian samples and the close geographical proximity with France we included in the aforementioned analyses all the Iberian samples from the 5th century BCE to the Mediaeval period (up to the 10th century CE). Only targets and source populations with a sample size larger than two were used in these analyses.

The early Neolithic period is characterised by the arrival of the first farmers in Western Europe and the Early Bronze Age is characterised by the arrival of populations related to or carrying a large proportion of their ancestry from pastoralist populations from the Eurasian steppes (50,74). Therefore, we added Anatolia_N and Russia_EBA_Yamnaya_Samara to the set of *right* populations: Mbuti, Han, AfontovaGora3, ElMiron, Karitiana, Kontenki14, MA1, Mota, Papuan, Ust_Ishim, Vestonice16, Villabruna, GoyetQ116-1, Morocco_Iberomaurusian, Anatolia_N and Russia_EBA_Yamnaya_Samara.

### aDNA Mediaeval samples

We generated whole-genome data for six ancient individuals from three different archaeological sites in western France, specifically from the region of *Pays-de-la-Loire*. Samples were dated using radiocarbon methods and estimates vary from the 4th to the 11th century CE. This interval corresponds to the early and High Mediaeval periods. Out of the six ancient individuals, four were sampled south of the Loire river while two were sampled north of the Loire. The archaeological excavations in Saint-Lupien, Rezé (south shore of the Loire) took place between 2005 and 2016 and were led by the team of Mikaël Rouzic under the request of the city of Rezé. The site located in Chaussé Saint-Pierre, Angers (north shore of the Loire), was excavated by the team of Martin Pithon, from the French Institute for Preventive Archaeological Research (INRAP) and the excavations occurred between July and August 2009. Based on the archaeological remains and radiocarbon dates, the site shows evidence of occupation from the beginning of the Roman Empire to modern times. The project was initialised under the request of the city of Angers given the plan for public construction affecting the archaeological site. The archaeological study of the site was authorised by the Regional Division for Cultural Affairs (Délégation Régional des affaires culturelles) and the INRAP. Finally, the excavations in the archaeological site in Chéméré (south shore of the Loire) started in the 60s but the two individuals sequenced in this study belong to a group of 181 individuals found in the last excavations in 2007. These excavations were led by an INRAP archaeological team under a project of preventive archaeology before construction of a pavilion. For more details on the description of the sites see Supplementary Material Online.

### aDNA library preparation and bioinformatic processing

One DNA extraction (following a modified version of the extraction method from(76–78) and one to two single indexed blunt end libraries(79) were prepared for each of the six ancient samples (fra001, fra004, fra008, fra009, fra016 and fra017). The DNA libraries were sequenced in two batches and over multiple lanes, first as a pilot run at the SciLife Sequencing Centre in Uppsala, Sweden, using Illumina HiSeq 2500 with paired-end 125 bp chemistry, and later in more depth at the CNRGH (Evry, France) using Illumina HiSeq X and with a paired-end 150 bp chemistry. Raw reads were trimmed with CutAdapt version 2.3(80) using the parameters -- quality-base 33, --quality-cutoff 15, -e 0.2, --trim-n and --minimum-length 15. Overlapping read pairs were then merged using FLASH version 1.2.11(81) and with parameters --min- overlap 11, --max-overlap 150 and --allow-outies. Merged fastq files were mapped to the human reference genome hs37d5 as single end reads using bwa-aln version 0.7.17(82) and parameters -l 16500, -n 0.01 and -o 2, as suggested for ancient DNA(83,84).

BAM files were merged to a per sample library level using the merge command in Samtools version 1.5 (85) before PCR duplicates were removed (reads with identical start and end positions were identified and collapsed) by a modified version of FilterUniqSAMPCons_cc.py, which ensures random assignment of bases in a 50/50 case. All reads longer than 35 base pairs and with less than 10% mismatches to the reference were kept, and a final merging step was performed for those samples with two sequenced libraries by merging the processed sample library BAMs to a final sample BAM.

Processed sample BAM files were then used to call pseudo-haploid genotypes using Samtools and the option *mpileup -R -B -q30 -Q30*. In order to merge with available datasets, the pseudo-haploid calling was carried out on the 593,124 autosomal sites on the Human Origins Array (Affymetrix). Contamination was estimated based on the X-chromosome and mitochondrial DNA using ANGSD(86) and schmutzi(87), respectively. Sample quality, inferred sex and contamination estimates are shown in the Table S2.1.

## Acknowledgments

This study received financial support from the Regional Council of Pays-de-la-Loire (VaCaRMe) and the Agence Nationale de la Recherche in France (ANR; FROGH, ANR-16- 599-CE12-0033). The PREGO bio-bank was built with the strong support from the Etablissement Français du Sang (EFS) and the Centre for Biological Resources (CRB) of CHU Nantes (BB-0033-00040). I.A. received funds from the People Programme (Marie Curie Actions) of the European Union’s Seventh Framework Programme (FP7/2007-2013) under REA grant agreement PCOFUND-GA-2013-609102, through the PRESTIGE programme (PRESTIGE-2017-4-0018) coordinated by Campus France. J.G. and R.R. were supported the Regional Council of Pays-de-la-Loire (GRIOTE) and the ANR (GenSud: ANR-14-CE10- 0001). The GAZEL Cohort Study was funded by EDF-GDF and INSERM, and received grants from the ‘Cohortes Santé TGIR Program’ of the ANR (ANR-08-BLAN-0028), Agence française de sécurité sanitaire de l’environnement et du travail (AFSSET; EST-2008/1/35), CAMIEG (Caisse d’assurance maladie des industries électrique et gazière) and CCAS (Caisse Centrale d’Activités Sociales du Personnel des Industries Électriques et Gazières). The extraction and WGS sequencing of blood samples was supported by the Laboratory of Excellence GENMED (ANR; grant no. ANR-10-LABX-0013, PIA: “Investissements d’Avenir’’ program). We also thank the CEPH-Biobank team for their involvement in DNA extractions for the GAZEL cohort. This work was supported by the France Génomique National infrastructure, funded as part of the PIA managed by the ANR (contract ANR-10-INBS-09). The authors also wish to thank the Fondation Genavie for its financial support. We are most grateful to the Genomics & Bioinformatics Core Facilities of Nantes (GenoA & BiRD, Biogenouest) and the Institut Français de Bioinformatique (IFB; ANR-11-INBS-0013) for their technical support. This work was supported by the *Institut National de Recherches Archéologiques Préventives* (Inrap), funded as part of the days allocated to research.

