## Supplemental file for "Genetic population structure across Brittany and the downstream Loire basin provides new insights on the demographic history of Western Europe"

### Supplementary Online Figures

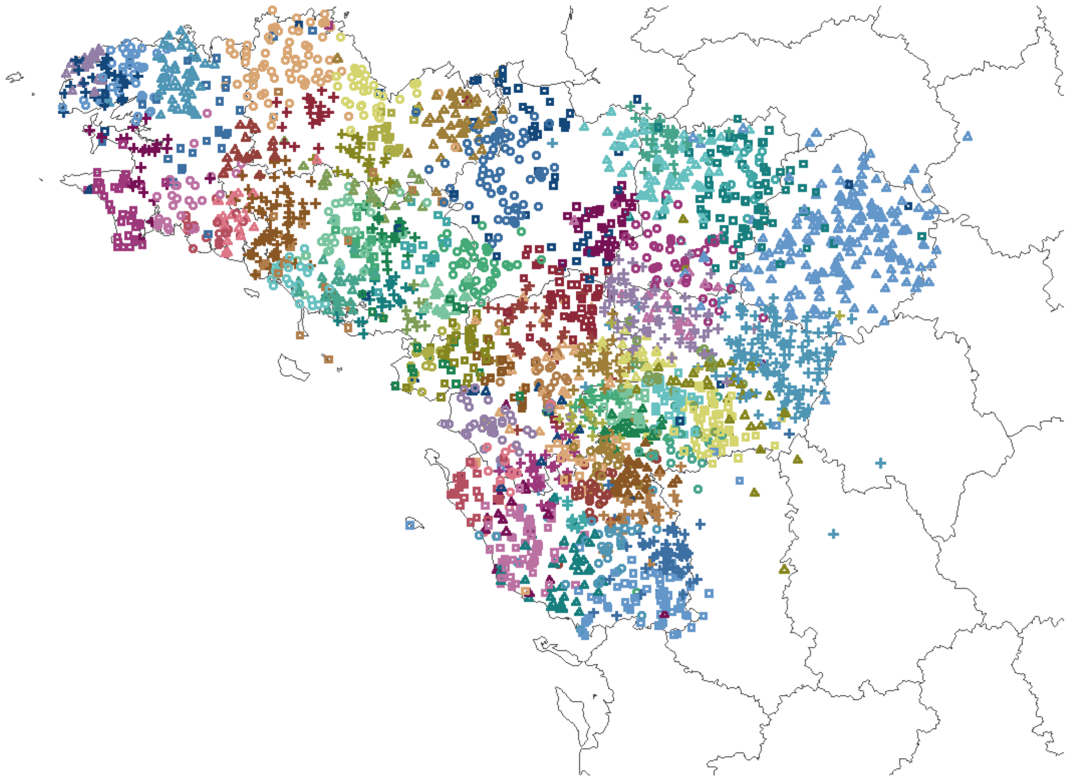

**Figure S1.1.** fineSTRUCTURE results for k=154. But only clusters with >10 individuals (78 clusters) are shown.

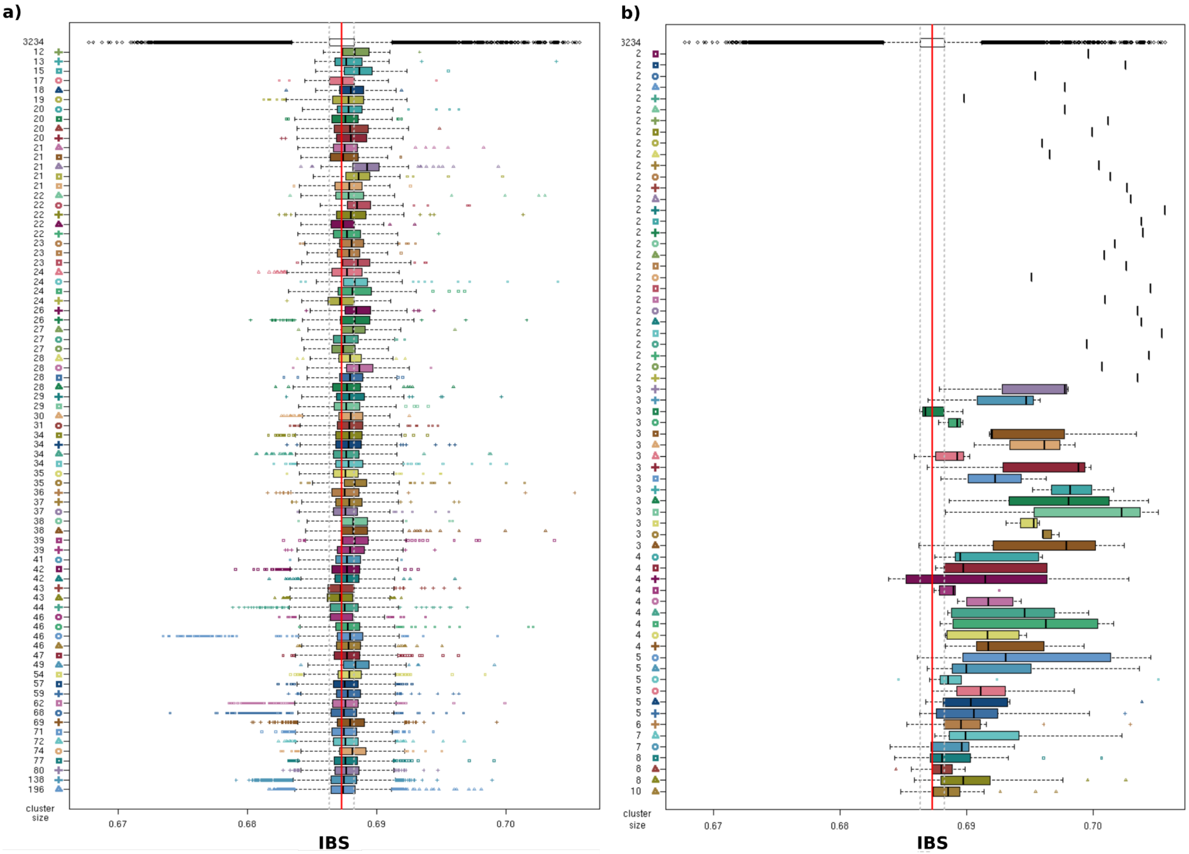


**Figure S1.2.** Identity-by-state (IBS) distances computed between pairs of individuals within the 154 clusters for the finest level of population structure. The white box plot represents the distribution of pairwise IBS values, computed in plink --*genome* option, across the whole dataset. Pairwise IBS values are shown in the x-axis while the y-axis shows the 78 clusters with n > 10 (**a**) and the remaining clusters with n <= 10 (**b**). Boxplots are coloured according to the cluster colours in Figure S1.1. The distributions of relatedness, here captured by the IBS distances, within clusters globally match that computed across the whole sample (median distribution within the 25th-75th percentiles). This occurs for most clusters with a n>10 (**a**). However, a departure from the whole distribution is observed for many of the clusters with a n < 10 (**b**). Given that fineSTRUCTURE detects groups of individuals with higher identity-by-descent, the smaller groups with increases IBS likely represent very recent shared ancestry (e.g., family relationships undetectable in the questionnaire, inbreeding).

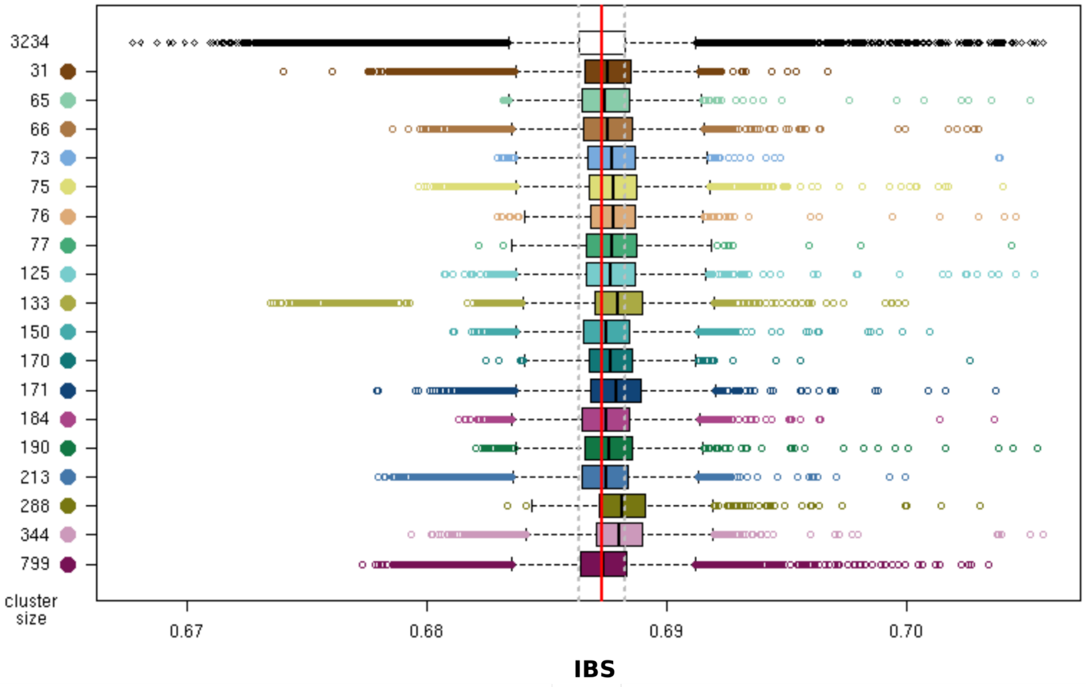


**Figure S1.3.** Identity-by-state (IBS) between pairs of individuals belonging to the 18 clusters inferred based on the TVD-based tree on fineSTRUCTURE clustering results. The distributions of relatedness within the 18 clusters (discussed in the main text) match that computed across the whole sample (white boxplot), indicating that individuals within clusters (n > 10) do not exhibit significantly more IBS than pairs of randomly selected individuals in the whole sample. Nevertheless, relatedness distributions slightly shift towards higher values as expected given that population structure is caused by shared ancestry. Such shared ancestry should be reflected over a large timescale and not only restricted to very recent periods (IBS ~ 1). Boxplots are coloured according to the cluster colors in Figure 1d (main text).

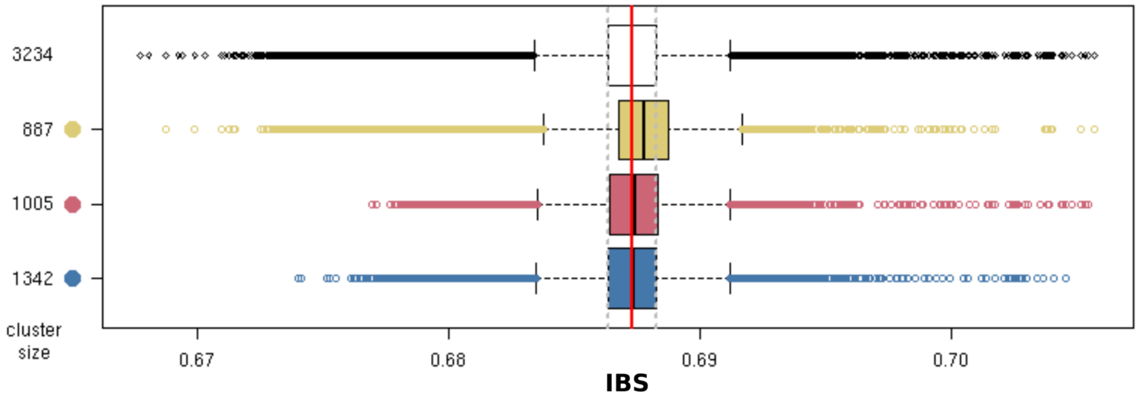


**Figure S1.4.** Identity-by-state (IBS) between pairs of individuals within the three clusters. The distributions of relatedness within the three clusters (Fig. 1a) match those computed across the whole sample (white boxplot), indicating that individuals within clusters (n > 10) do not exhibit significantly more IBS than pairs of randomly selected individuals in the whole sample. Overall, we conclude that fine-scale structure found in our study is not due to an overrepresentation of recently related samples.

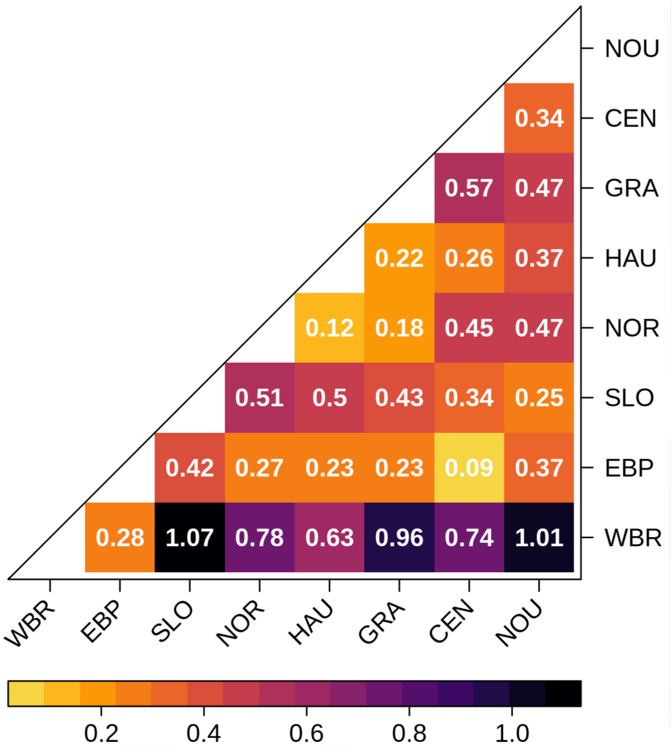


**Figure S1.5 -** Pairwise average Weir and Cockerham's $F_{ST}$s within France x 1000**.** $F_{ST}$ values were computed on the WGS dataset after pruning for LD and using 25 samples per population with the exception of Normandy whose n=19. Cluster names: Western Brittany (WBR), Eastern Brittany/*Pays-de-la-Loire* (EBP) and South Loire (SLO). Population acronyme: NOR, *Normandie*; HAU, Hauts-de-France; GRA, Grand Est; CEN, Centre-Val de Loire; NOU, Nouvelle-Aquitaine.

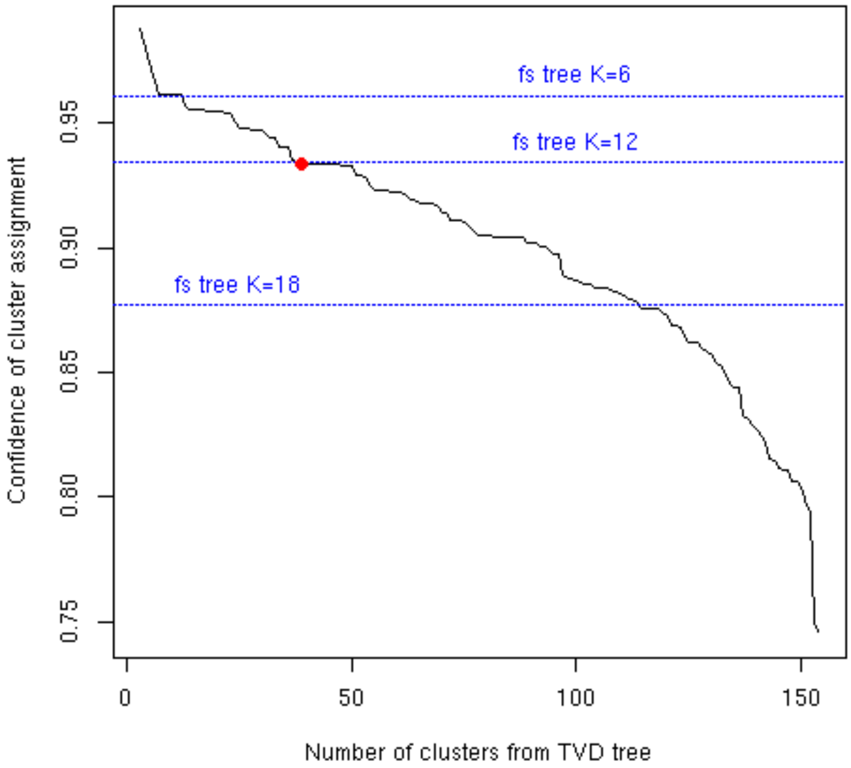


**Figure S1.6.** Confidence of cluster assignment from fineSTRUCTURE and TVD-based trees. FS-tree has lower confidence of cluster assignment than TVD-based tree for the same levels of number of clusters. For example, cluster assignment confidence retrieved for k=39 in TVD-based tree (red point) is similar for k=12 in FS-tree. See Material and Methods for details on the computation of cluster confidence (ChromoPainter and fineSTRUCTURE).

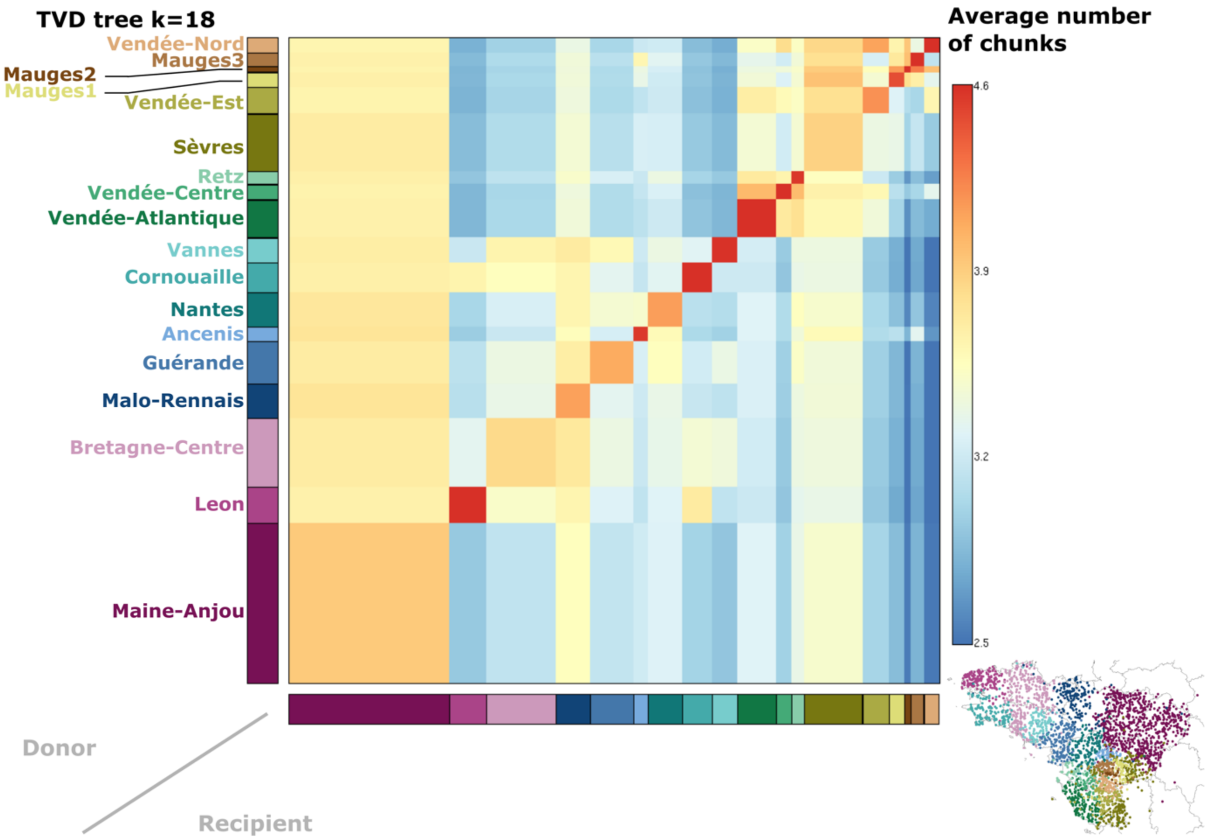


**Figure S1.7.** Population-level coancestry matrix for k=18. This matrix was based on the individual-level coancestry (chunkcount matrix) used in cluster inference. Along the columns is shown the average chunk counts a recipient cluster (bottom) received from the donor clusters (on the left) while across the rows is shown the average chunk count that donor clusters (on the left) have contributed to the recipient clusters (bottom). The diagonal represents the average chunk count between pairs of individuals assigned to the same cluster, i.e. the "drift component". In order to visualise the bulk of the variation, only values between 1-99 percentile were considered and those exceeding the range were coloured according to the closer 1 or 99 percentile value. Warmer colours indicate higher levels of ancestry sharing.

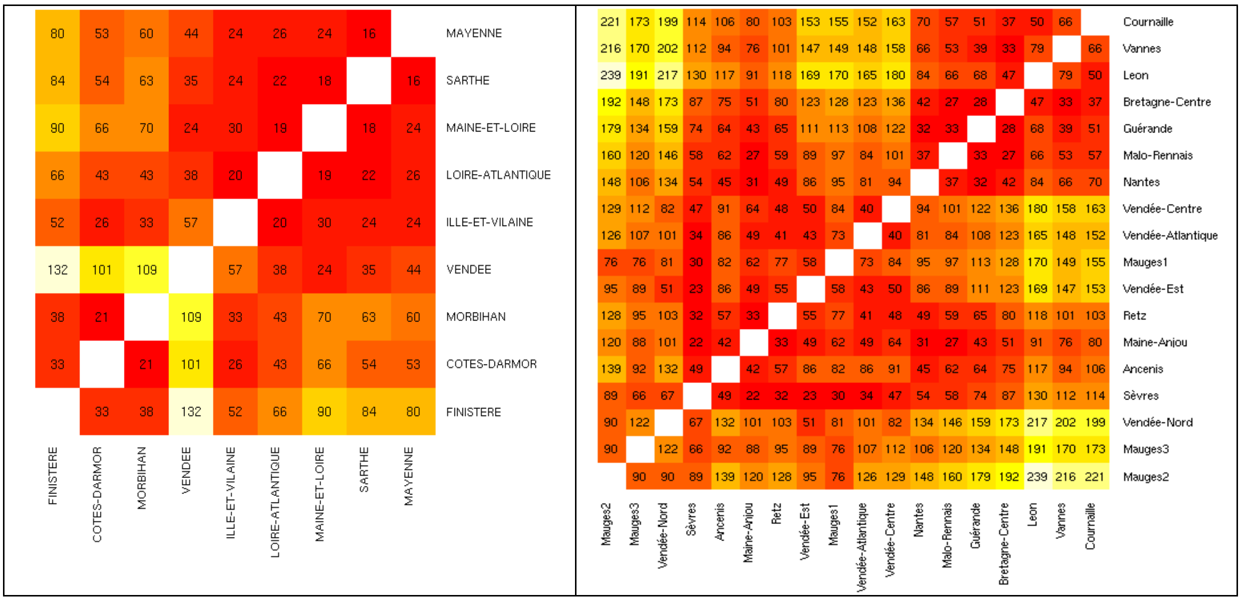


**Figure S1.8.** Pairwise F $_{ST}$ values multiplied by 100,000 between the *départements* of Brittany and *Pays-de-la-Loire* (lower panel) and the 18 clusters in Northwestern France (upper panel) identified in PREGO dataset with fineSTRUCTURE (Fig. 1d, main text).

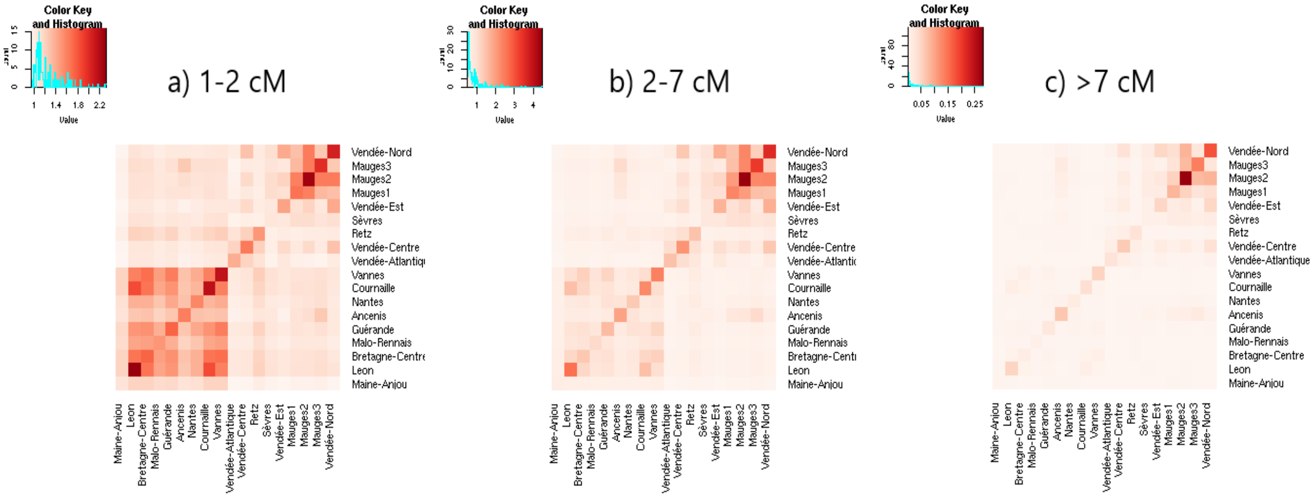


**Figure S1.9.** Identity-by-descent (IBD) sharing between pairs of individuals belonging to the 18 clusters identified with the TVD-based tree on fineSTRUCTURE (Fig. 1d main text) results for different chromosome segment sizes (a-c). Segment sizes are indicated over the heatmaps. Heatmap columns (from bottom to up) and rows (left to right) are ordered similarly to the coancestry matrices (see above).

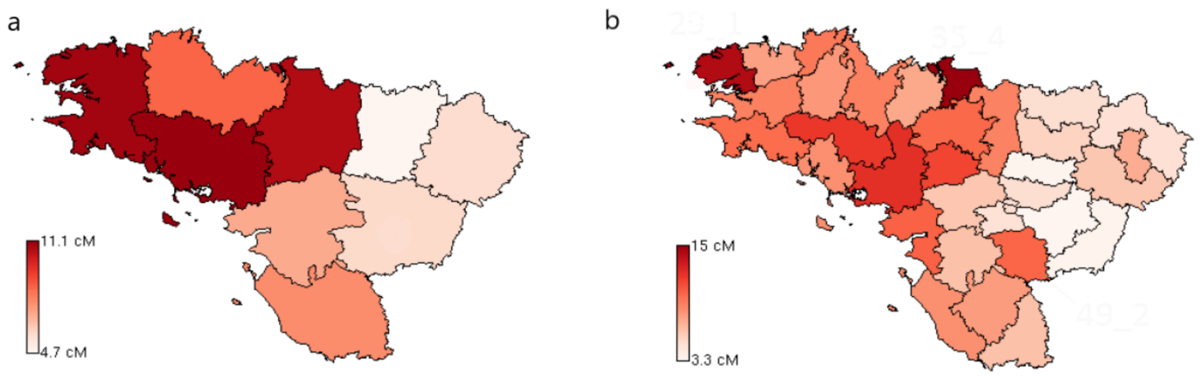


**Figure S1.10.** Heatmap displaying average length of runs of homozygosity (ROH) across individuals within **a)** *départements* and **b)** *arrondissements* of Northwestern France. The regions of Brittany and Mauges display large ROH indicating smaller effective population sizes.

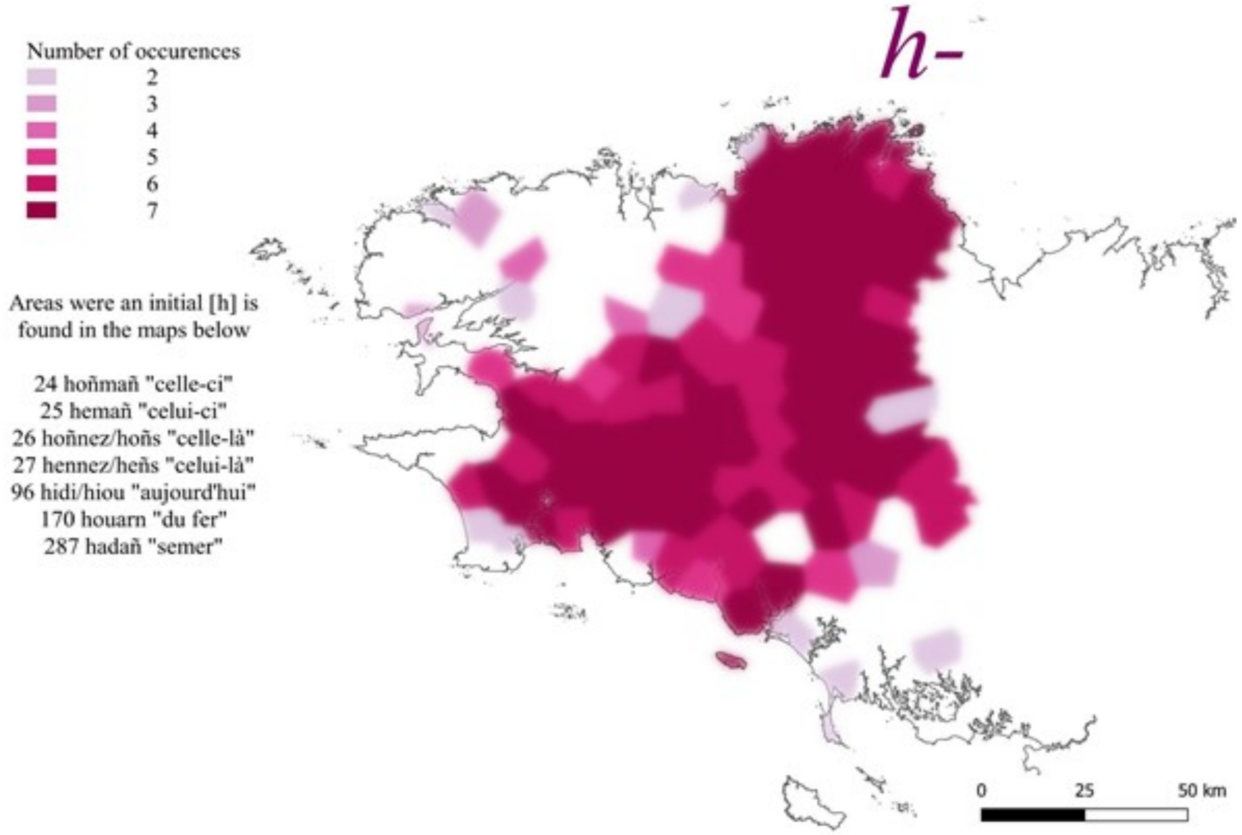

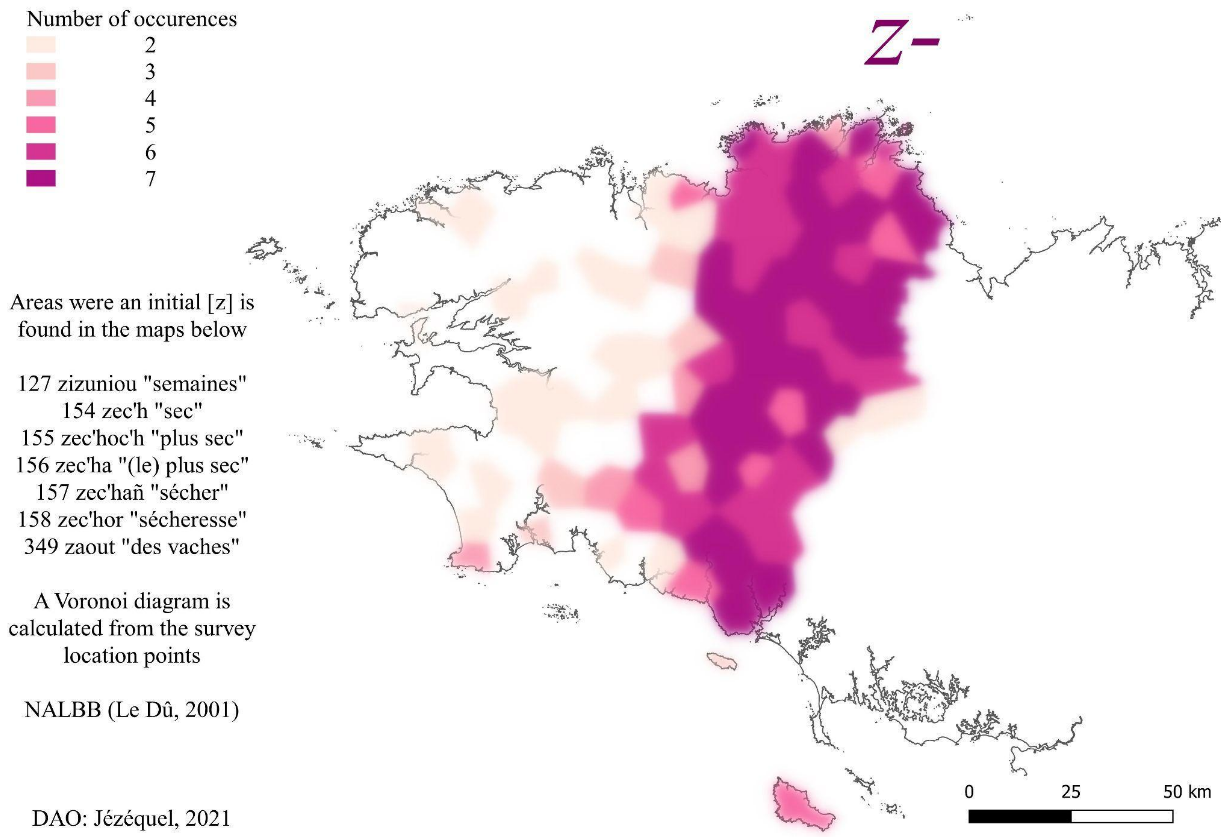


**Fig. S1.11.** Treatments of initial h- and z- in Lower-Brittany in correlation with the cluster “Bretagne-Centre”.

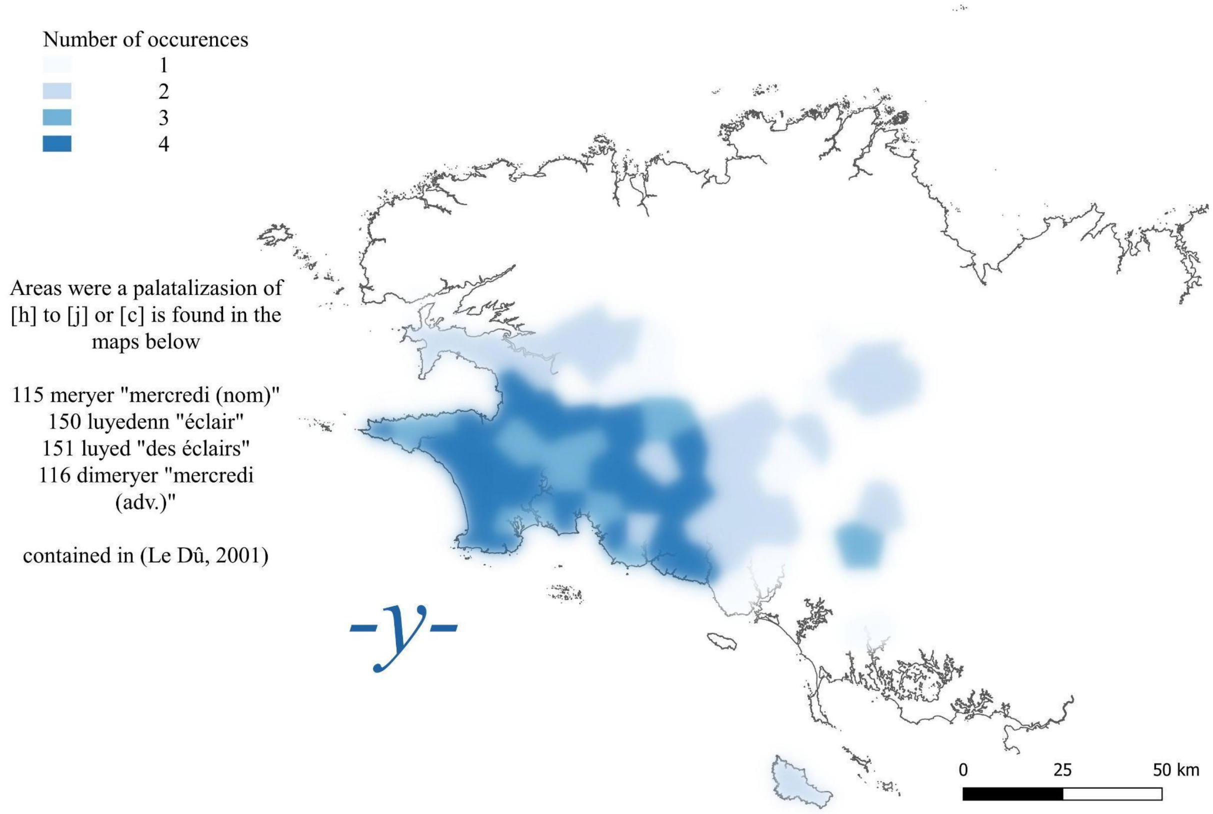


**Figure S1.12.** Palatalisation of the *h* in *y* in correlation with cluster "Cornouaille".

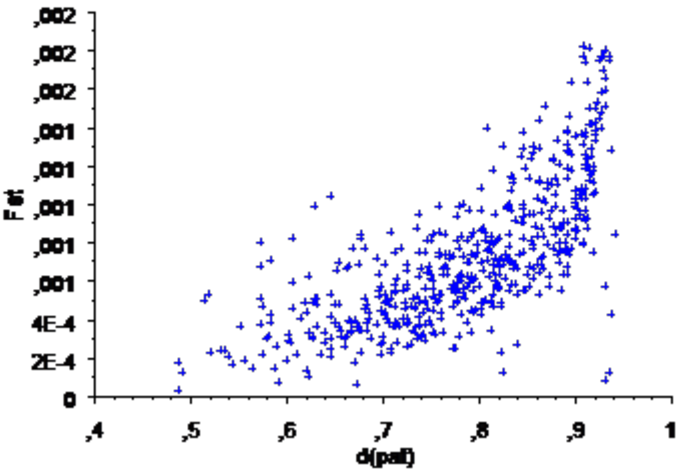


**Figure S1.13.** Correlation between surname distance and $F_{ST}$ (negative $F_{ST}$ removed).

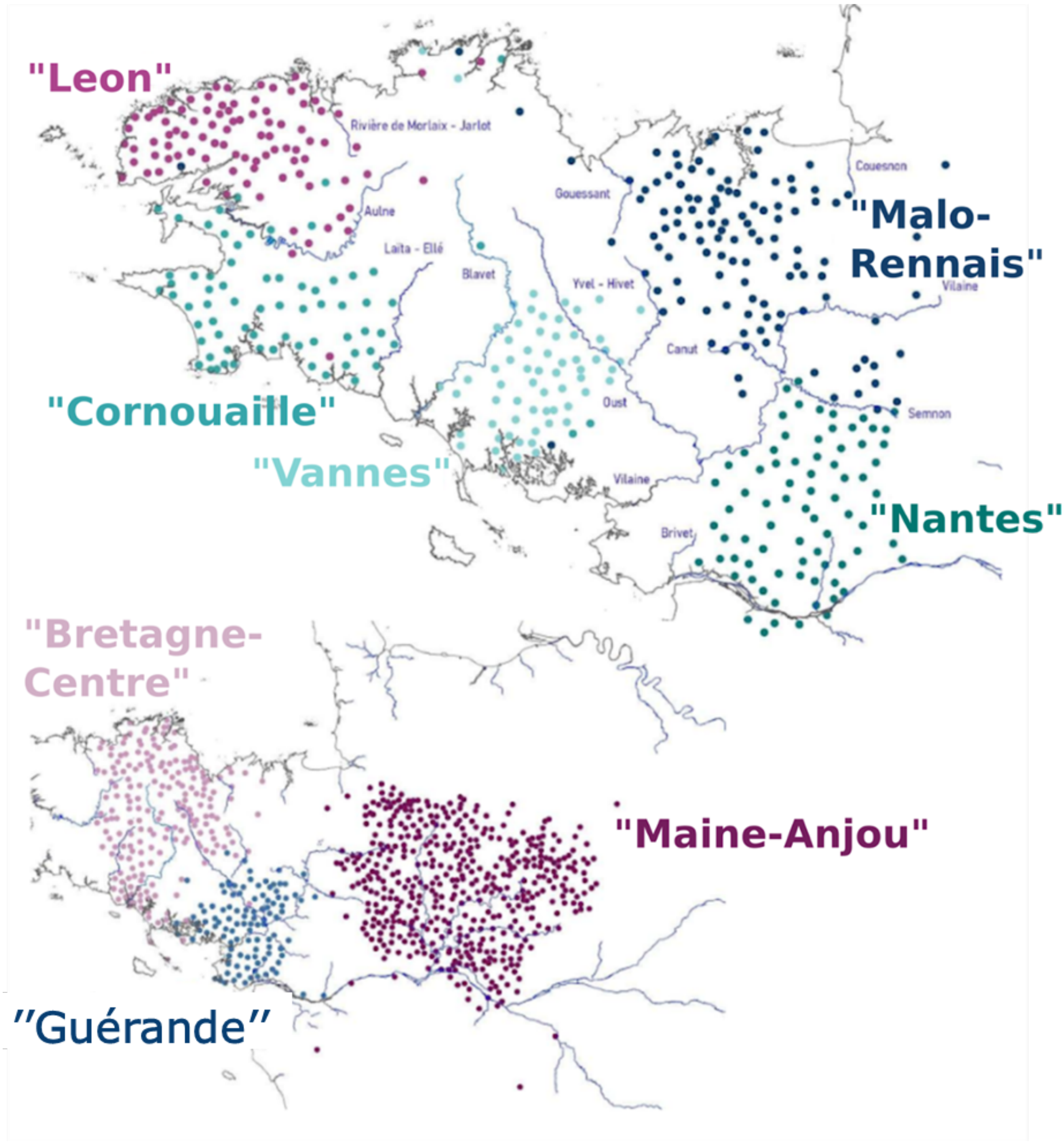
"Leon"
Rivière de Morlaix - Jarlot
Aulne
Laita - Ellé
Blavet
Gouessant
Couesnon
"Malo-Rennais"
Vilaine
Yvel - Hivet
Canut
Oust
Semnon
"Cornouaille"
"Vannes"
Vilaine
Brivet
"Nantes"
"Bretagne-Centre"
"Maine-Anjou"
"Guérande"

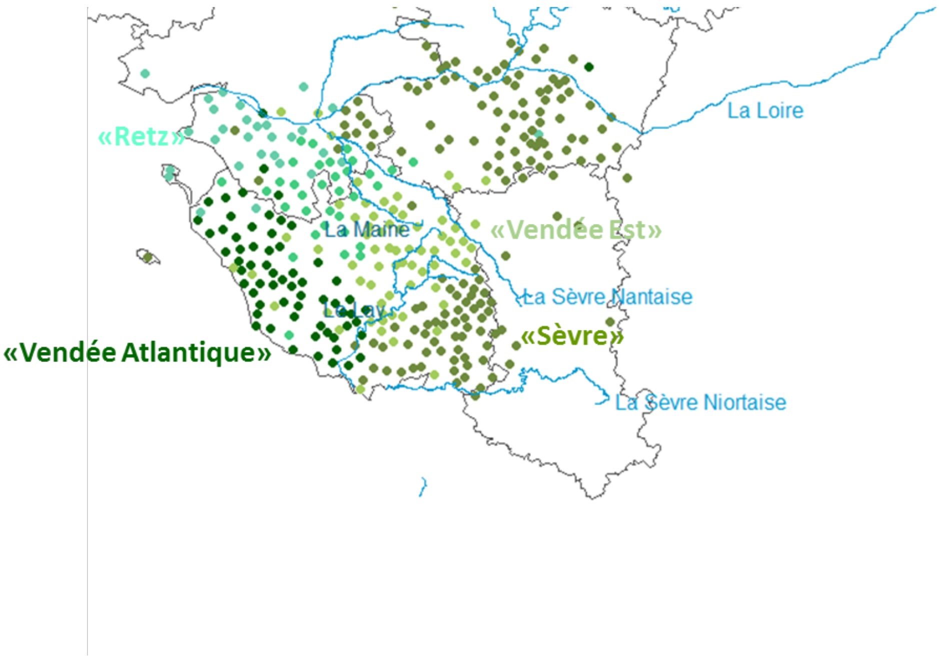


**Figure S1.14 -** Clusters inferred with fineSTRUCTURE (k=18 of the TVD-tree) and the distribution of local water bodies. The map is split in three parts and clusters in the Mauges region ("Mauges 1", "Mauges 2" and "Mauges 3") were not displayed to facilitate its visualisation.

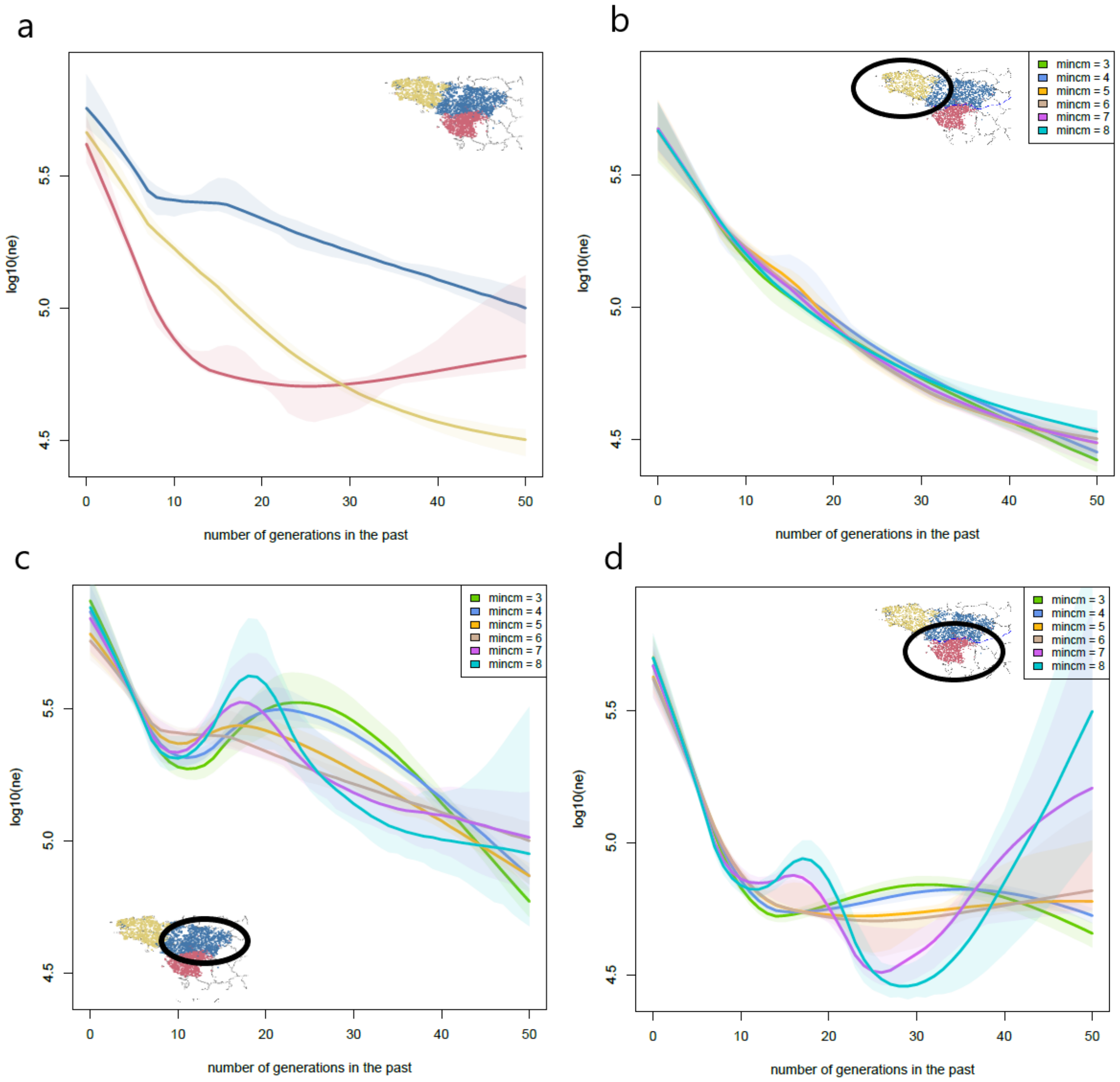


**Figure S1.15.** Effective population size trajectories for the three clusters inferred by fineSTRUCTURE (Fig. 1a, main text) obtained with IBDNe. **a)** IBDNe results for the three clusters together using minimum fragment length (*mincm*) of 6 cM. **b-d)** IBDNe results for each of the clusters shown in the inset across multiple *mincm* values.

<table>
<tr><th></th><th>MAC 2</th><th>MAC 3-10</th></tr>
<tr><td>K=2</td><td>
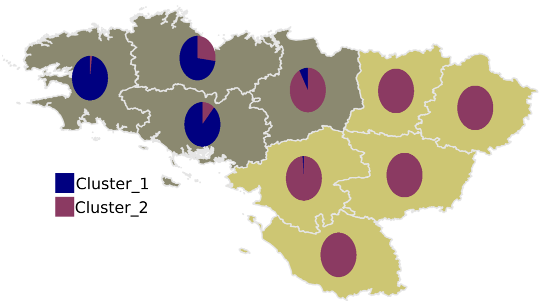

</td><td>
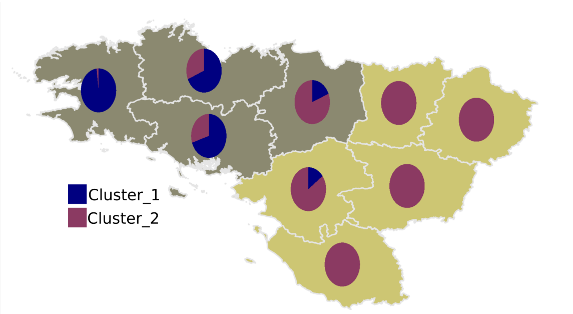

</td></tr>
<tr><td>K=4</td><td>
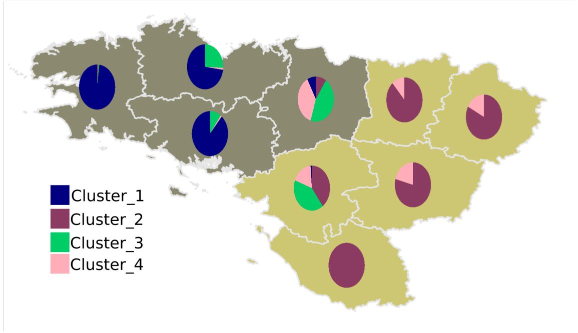

</td><td>
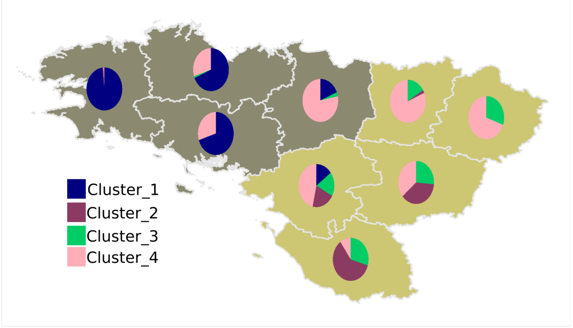

</td></tr>
<tr><td>K=5</td><td>
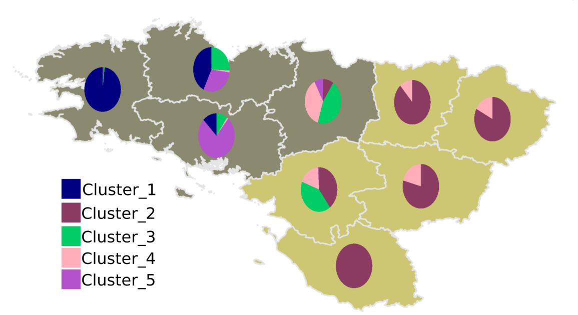

</td><td>
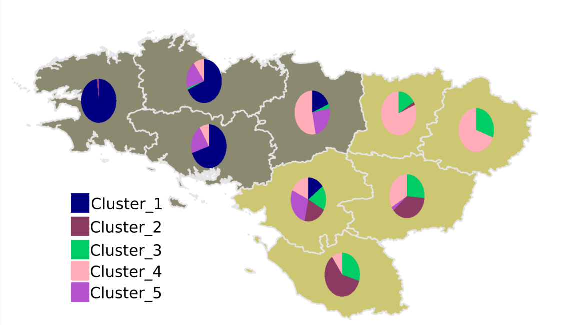

</td></tr>
<tr><td>K=6</td><td>
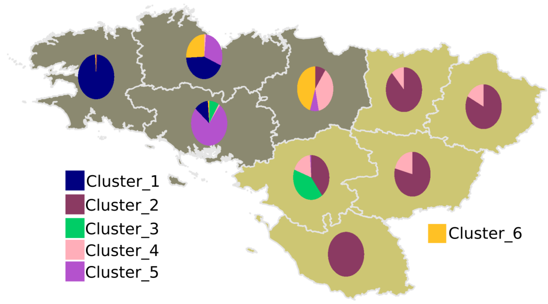

</td><td>
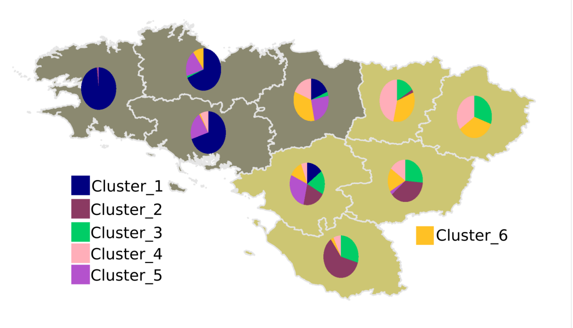

</td></tr>
<tr><td>K=7</td><td></td><td></td></tr>
</table>

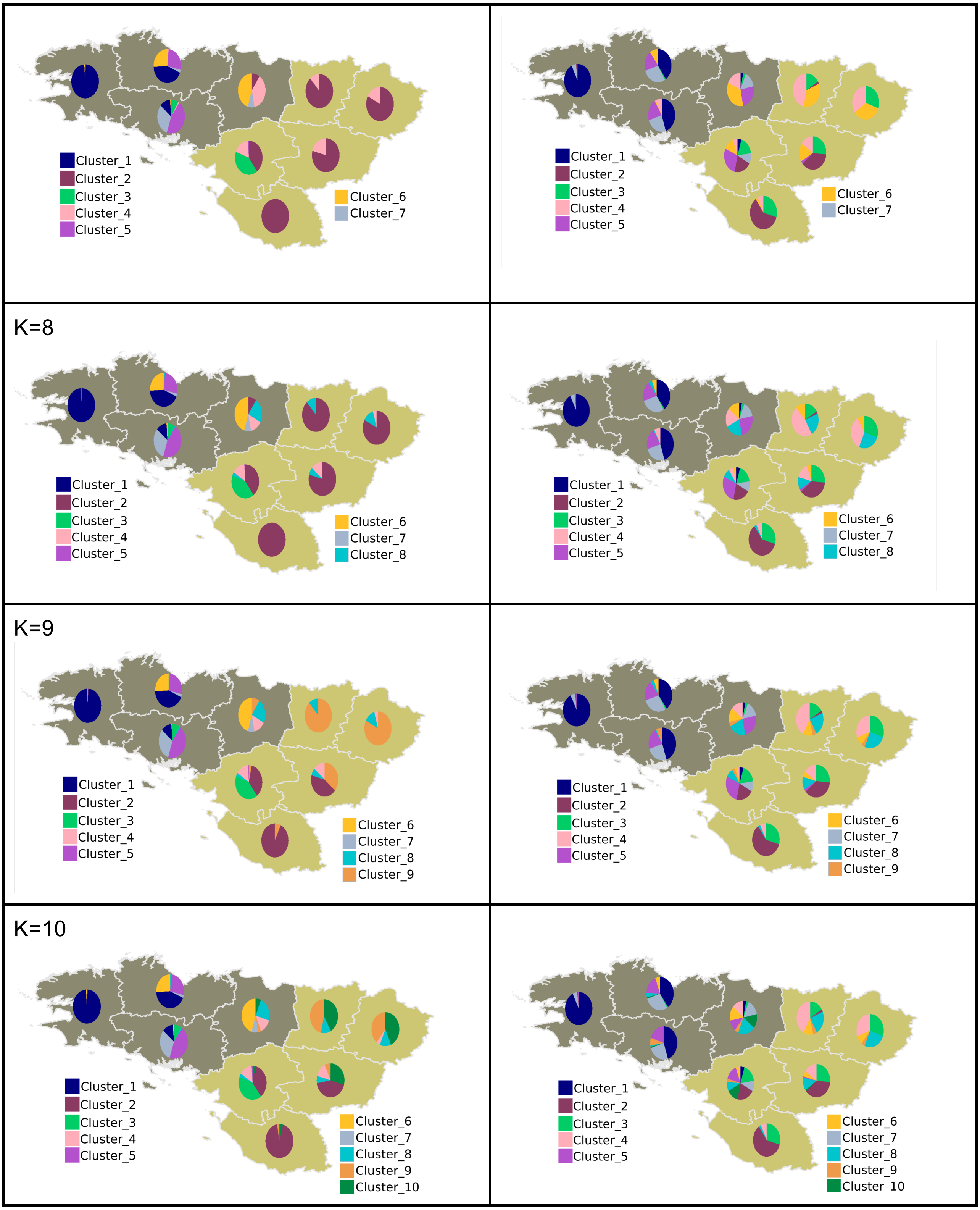


**Figure S2.1. -** Distribution of individuals based on the hierarchical clustering performed on the allele-sharing matrices. Rows show different *k* values used in the hierarchical clustering and columns show the results for different allele count classes. Allele sharing matrices were obtained by randomly selecting 1 million variable sites within each of the two allele count categories.

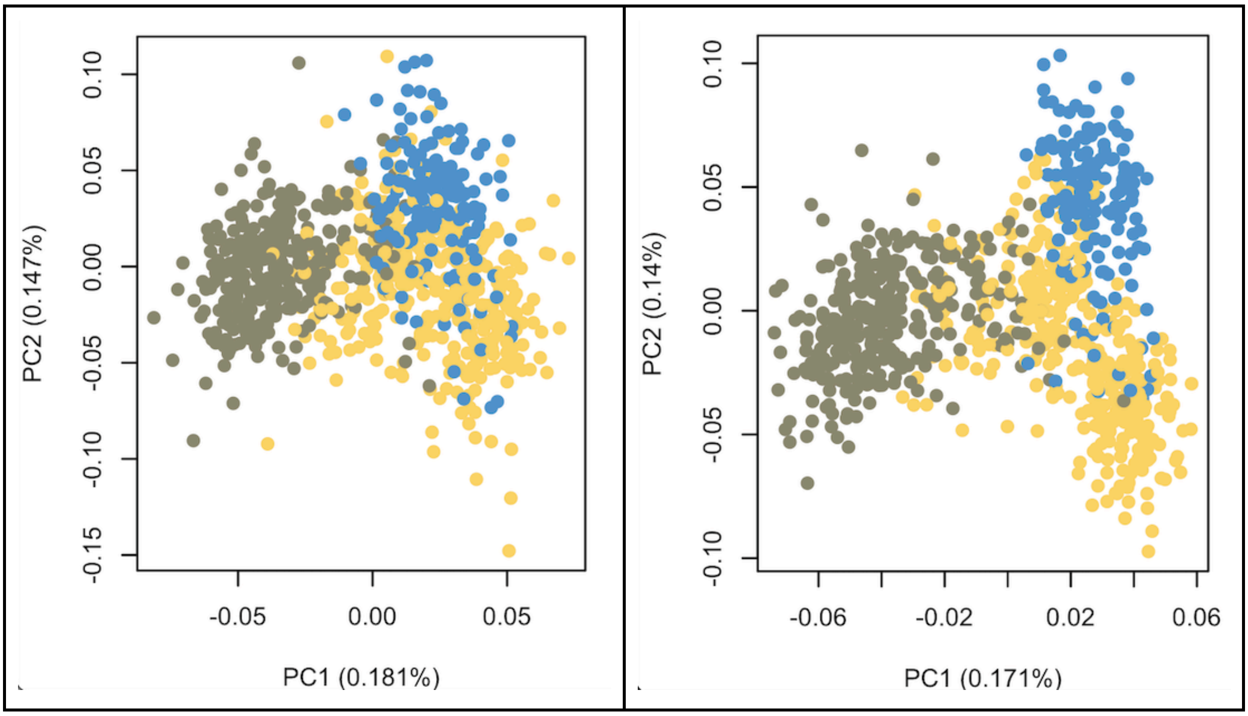


**Figure S2.2 -** PCA on the WGS dataset containing only French populations for which the all the four grand parents belong to the same *commune* (843 samples). On the left panel, PCA was performed on common sites, MAF > 10% (~443,933 sites). The right panel shows the PCA low-frequency variants (1% < MAF < 10%, ~863,141 sites). Samples from Brittany are coloured in dark green and samples from Northeastern and Southwestern France are coloured in blue and yellow, respectively. See Génin *et al*. (manuscript in preparation) for further details on the genetic structure within France.

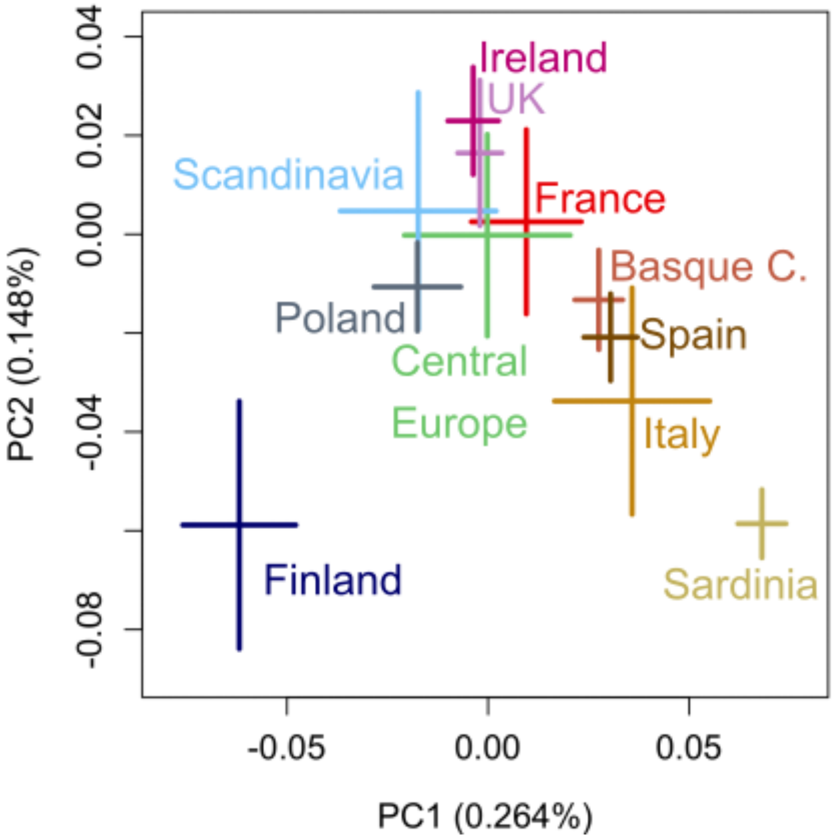


**Figure S3.1** The two first principal components of genetic variation in the "modern merged dataset", which includes 843 French WGSs (red) for which the birth places of the four grand-parents are situated in close proximity, and genome-wide data from 20 central and western European populations: Basque Country (HGDP), Belgium, Cornwall (UK, POBI), Denmark, Dyfed (UK, POBI), Finland, Germany, Gwynedd (UK, POBI), Ireland, Italy, Kent (UK, POBI), Norfolk (UK, POBI), Northern Ireland, Norway, Orkney Islands (HGDP), Poland, Sardinia (HGDP), Spain, Sweden, United Kingdom. For simpler visualisation samples from Norway, Sweden and Denmark were labelled as "Scandinavia", samples from Belgium and Germany were labelled as "Central Europe". All samples from Great Britain are represented by UK. Only samples from the FranceGenRef are shown in the plot. Crosses represent 2*standard deviations of the PC distribution for each group.

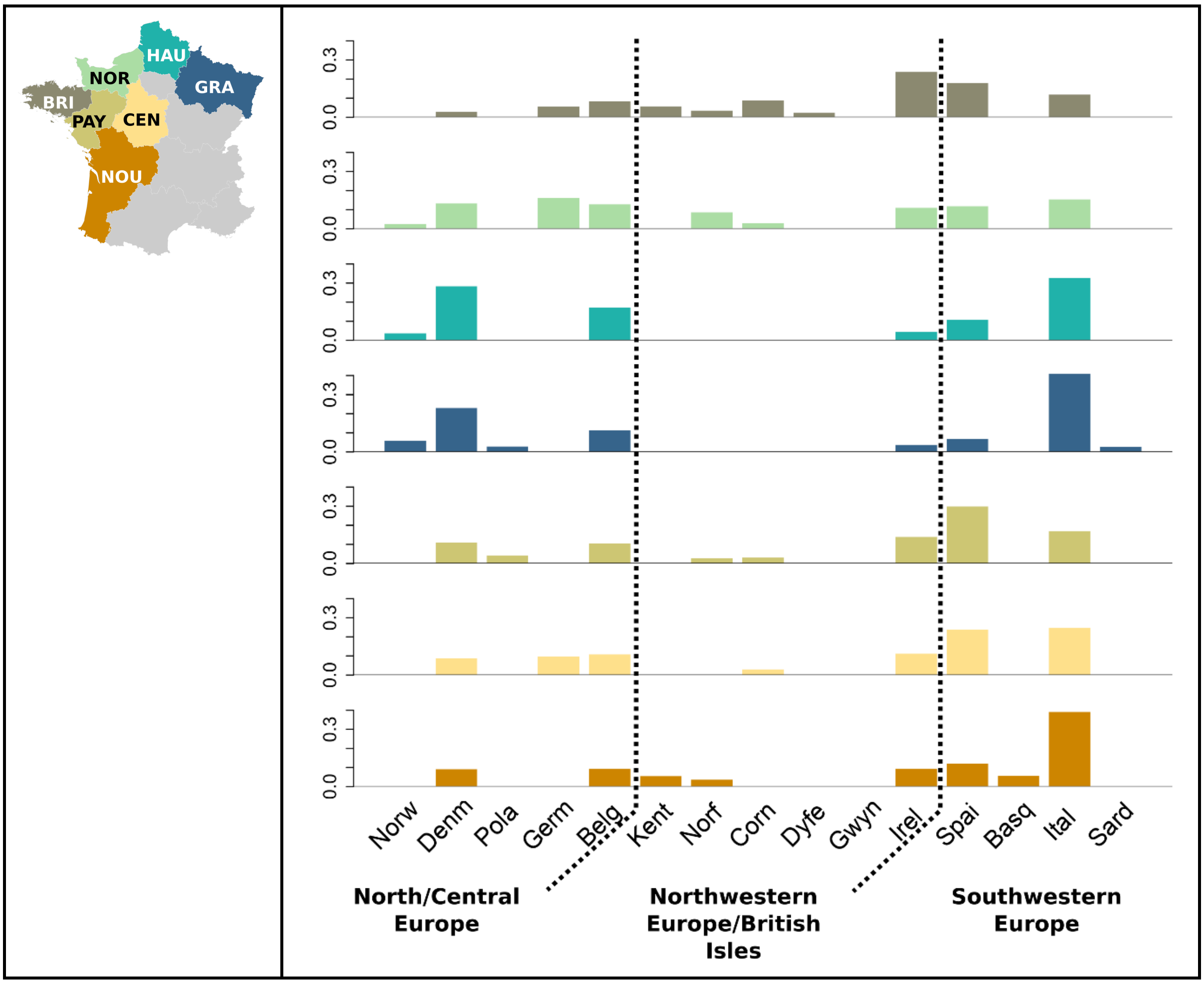


**Figure S3.2. -** Ancestry profiles obtained with GLOBETROTTER for each of the French regions shown on the left panel (rows). Ancestry contributions shown in the y-axis from each of the European samples in the x-axis. French regions are coloured according to the map on the left pane. Population acronyme: BRI, Brittany; PAY, Pays-de-la-Loire; NOR, *Normandie*; HAU, Hauts-de-France; GRA, Grand Est; CEN, Centre-Val de Loire; NOU, Nouvelle-Aquitaine.

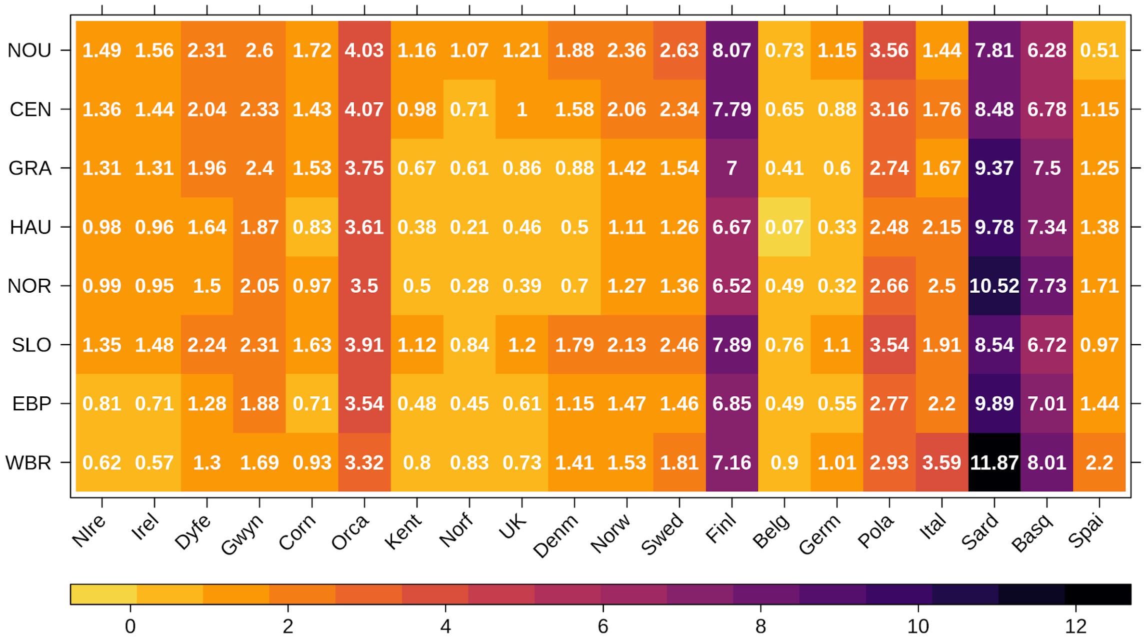


**Figure S3.3. -** Pairwise average Weir and Cockerham's $F_{ST}$s between French and other European populations x 1000. French clusters and populations are: WBR, Western Brittany; EBP, Eastern Brittany and Pays-de-la-Loire; SLO, South Loire; NOR, *Normandie*; HAU, Hauts-de-France; GRA, Grand Est; CEN, Centre-Val de Loire; NOU, Nouvelle-Aquitaine. Non-French population acronyms: Basque Country (Basq), Belgium (Belg), Cornwall (Corn), Denmark (Denm), Dyfed (Dyfe), Finland (Finl), Germany (Germ), Gwynedd (Gwyn), Ireland (Irel), Italy (Ital), Kent (Kent), Norfolk (Norf), Northern Ireland (NIrel), Norway (Norw), Orkney Islands (Orca), Poland (Pola), Sardinia (Sard), Spain (Spai), Sweden (Swed), United Kingdom (UK).

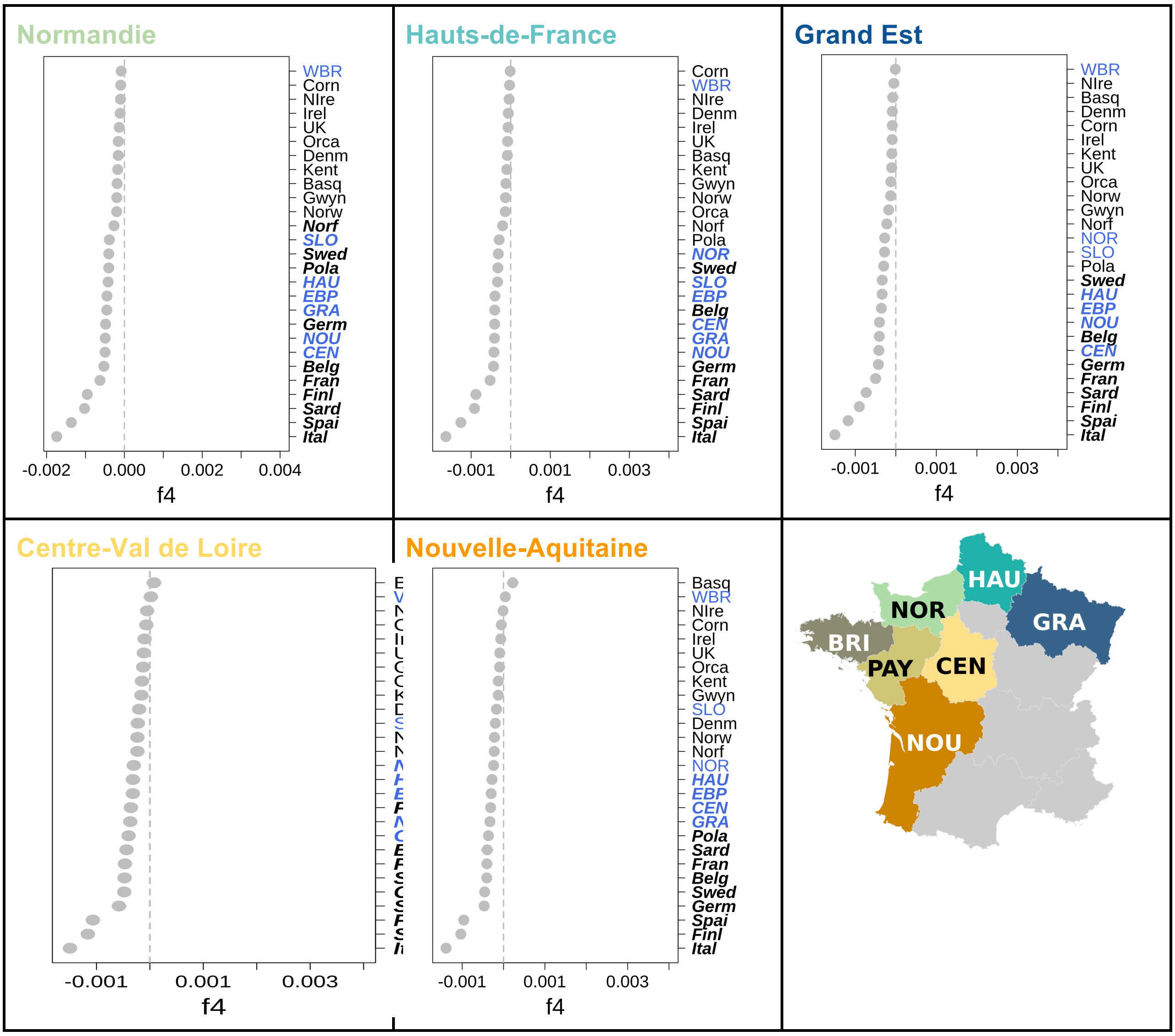


**Figure S3.4. -** $f_4$*-statistics* of the form $f_4$(Mbuti, French region; Dyfed, X), where the French region is indicated on top of the plot and X is each of the populations on the right side of the plot. $F_4$-statistics values are shown together with one standard deviation. French clusters and populations are: WBR, Western Brittany; EBP, Eastern Brittany and Pays-de-la-Loire; SLO, South Loire; NOR, *Normandie*; HAU, Hauts-de-France; GRA, Grand Est; CEN, Centre-Val de Loire; NOU, Nouvelle-Aquitaine.

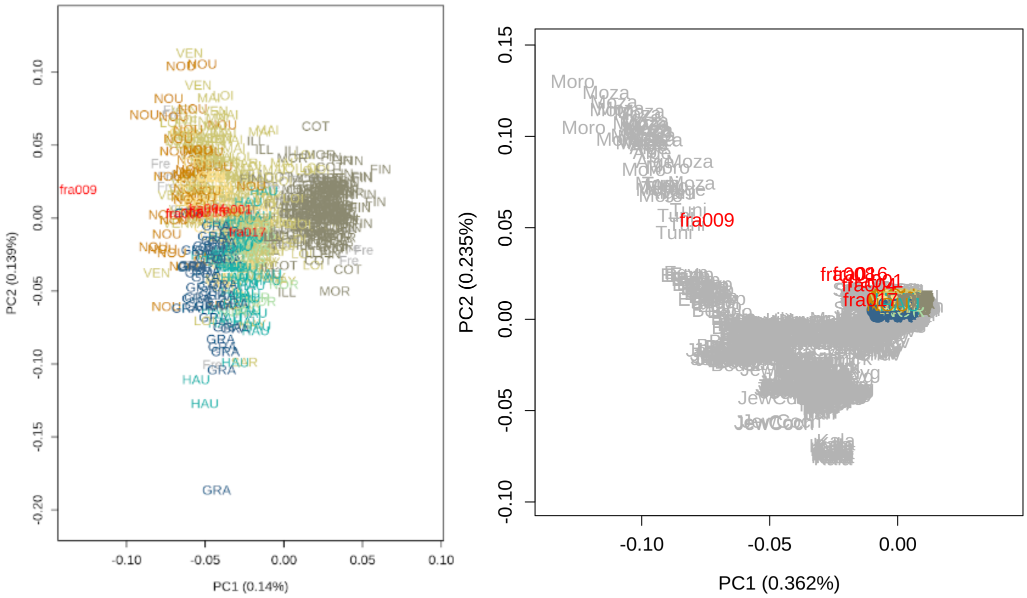


**Figure S4.1 -** Mediaeval French samples projected onto the two first principal components from present-day Europeans and north Africans. Modern European genotypes present in the Human Origins Array (Reich's Lab, vs.42.2 March 2020 release) are coloured in grey. Modern French samples analysed in this study are coloured according as in Fig. S3.3 (above) whereas ancient samples are represented in red. North African sample acronyms: Tuni - Tunisia; Moza - Mozabites; Moro-Morocco. The pca was performed with smartpca (lsqproject: YES).

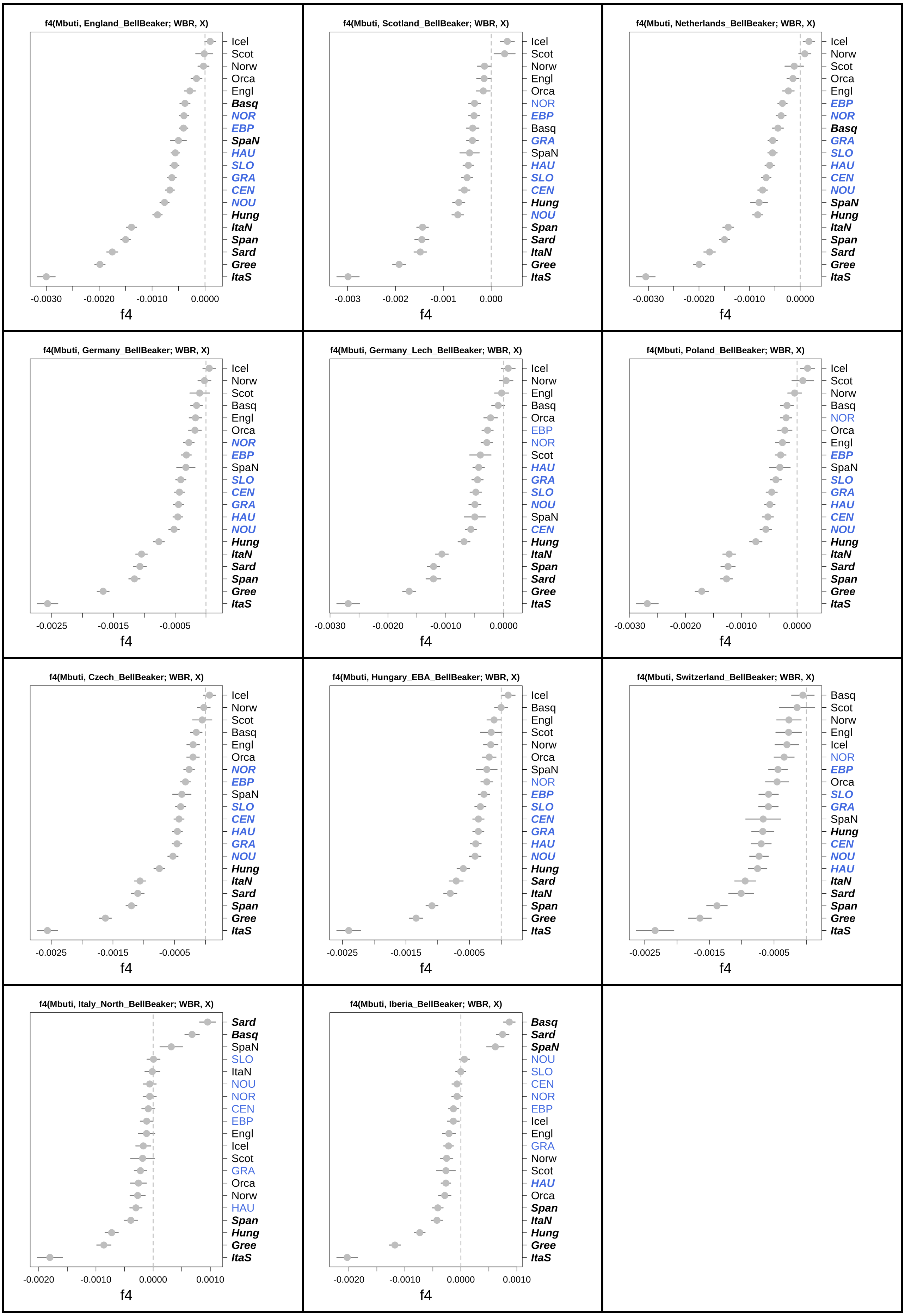

f4(Mbuti, England_BellBeaker; WBR, X)
Icel
Scot
Norw
Orca
Engl
Basq
NOR
EBP
SpaN
HAU
SLO
GRA
CEN
NOU
Hung
ItaN
Span
Sard
Gree
ItaS
-0.0030
-0.0020
-0.0010
0.0000
f4
f4(Mbuti, Scotland_BellBeaker; WBR, X)
Icel
Scot
Norw
Engl
Orca
NOR
EBP
Basq
GRA
SpaN
HAU
SLO
CEN
Hung
NOU
Span
Sard
ItaN
Gree
ItaS
-0.003
-0.002
-0.001
0.000
f4
f4(Mbuti, Netherlands_BellBeaker; WBR, X)
Icel
Norw
Scot
Orca
Engl
EBP
NOR
Basq
GRA
SLO
HAU
CEN
NOU
SpaN
Hung
ItaN
Span
Sard
Gree
ItaS
-0.0030
-0.0020
-0.0010
0.0000
f4
f4(Mbuti, Germany_BellBeaker; WBR, X)
Icel
Norw
Scot
Basq
Engl
Orca
NOR
EBP
SpaN
SLO
CEN
GRA
HAU
NOU
Hung
ItaN
Sard
Span
Gree
ItaS
-0.0025
-0.0015
-0.0005
f4
f4(Mbuti, Germany_Lech_BellBeaker; WBR, X)
Icel
Norw
Engl
Basq
Orca
EBP
NOR
Scot
HAU
GRA
SLO
NOU
SpaN
CEN
Hung
ItaN
Span
Sard
Gree
ItaS
-0.0030
-0.0020
-0.0010
0.0000
f4
f4(Mbuti, Poland_BellBeaker; WBR, X)
Icel
Scot
Norw
Basq
NOR
Orca
Engl
EBP
SpaN
SLO
GRA
HAU
CEN
NOU
Hung
ItaN
Sard
Span
Gree
ItaS
-0.0030
-0.0020
-0.0010
0.0000
f4
f4(Mbuti, Czech_BellBeaker; WBR, X)
Icel
Norw
Scot
Basq
Engl
Orca
NOR
EBP
SpaN
SLO
CEN
HAU
GRA
NOU
Hung
ItaN
Sard
Span
Gree
ItaS
-0.0025
-0.0015
-0.0005
f4
f4(Mbuti, Hungary_EBA_BellBeaker; WBR, X)
Icel
Basq
Engl
Scot
Norw
Orca
SpaN
NOR
EBP
SLO
CEN
GRA
HAU
NOU
Hung
Sard
ItaN
Span
Gree
ItaS
-0.0025
-0.0015
-0.0005
f4
f4(Mbuti, Switzerland_BellBeaker; WBR, X)
Basq
Scot
Norw
Engl
Icel
NOR
EBP
Orca
SLO
GRA
SpaN
Hung
CEN
NOU
HAU
ItaN
Sard
Span
Gree
ItaS
-0.0025
-0.0015
-0.0005
f4
f4(Mbuti, Italy_North_BellBeaker; WBR, X)
Sard
Basq
SpaN
SLO
ItaN
NOU
NOR
CEN
EBP
Engl
Icel
Scot
GRA
Orca
Norw
HAU
Span
Hung
Gree
ItaS
-0.0020
-0.0010
0.0000
0.0010
f4
f4(Mbuti, Iberia_BellBeaker; WBR, X)
Basq
Sard
SpaN
NOU
SLO
CEN
NOR
EBP
Icel
Engl
GRA
Norw
Scot
HAU
Orca
Span
ItaN
Hung
Gree
ItaS
-0.0020
-0.0010
0.0000
0.0010
f4

**Figure S4.2 -** $f_4$ of the form *$f_4$ (Mbuti, Bell Beaker population; WBR, X)*, where Bell Beaker population is indicated on top of the plot and X is each of the populations on the y-axis (right side of the plot). French populations are coloured in blue and comparisons with |Z| > 3 are indicated in bold on the y-axis.

### Supplementary Online Tables

**Table S1.1 - Outgroup f3-statistics of the form f3(Mbuti; Source 1, Source 2). Only the top 5 values are shown.**

| Source_1 | Source_2 | Target | f_3 | std.err | Z | # SNPs |
|---|---|---|---|---|---|---|
| WBR | Dyfed | Mbuti | 0.291993 | 0.002451 | 119.115 | 411777 |
| WBR | NIreland | Mbuti | 0.291794 | 0.002439 | 119.652 | 411783 |
| WBR | Ireland | Mbuti | 0.291773 | 0.002455 | 118.869 | 411782 |
| WBR | Cornwall | Mbuti | 0.291668 | 0.002441 | 119.492 | 411774 |
| WBR | Orcadian | Mbuti | 0.291541 | 0.002449 | 119.067 | 411701 |
| EBP | Dyfed | Mbuti | 0.290345 | 0.002441 | 118.933 | 411784 |
| EBP | WBR | Mbuti | 0.29021 | 0.002429 | 119.477 | 411773 |
| EBP | Cornwall | Mbuti | 0.290135 | 0.002427 | 119.561 | 411774 |
| EBP | Ireland | Mbuti | 0.290085 | 0.002441 | 118.846 | 411774 |
| EBP | NIreland | Mbuti | 0.290044 | 0.002434 | 119.146 | 411796 |
| NOR | Dyfed | Mbuti | 0.290544 | 0.002468 | 117.737 | 411765 |
| NOR | WBR | Mbuti | 0.290236 | 0.002455 | 118.236 | 411777 |
| NOR | Cornwall | Mbuti | 0.290189 | 0.002454 | 118.271 | 411746 |
| NOR | NIreland | Mbuti | 0.290166 | 0.002452 | 118.348 | 411780 |
| NOR | Ireland | Mbuti | 0.290135 | 0.002467 | 117.628 | 411773 |
| HAU | Dyfed | Mbuti | 0.290145 | 0.002448 | 118.539 | 411778 |
| HAU | Cornwall | Mbuti | 0.290092 | 0.002435 | 119.119 | 411788 |
| HAU | WBR | Mbuti | 0.290035 | 0.002434 | 119.151 | 411780 |
| HAU | NIreland | Mbuti | 0.289996 | 0.002439 | 118.891 | 411785 |
| HAU | Denmark | Mbuti | 0.289917 | 0.002423 | 119.653 | 411789 |
| GRA | Dyfed | Mbuti | 0.289879 | 0.00244 | 118.782 | 411773 |
| GRA | WBR | Mbuti | 0.289829 | 0.002442 | 118.706 | 411792 |
| GRA | NIreland | Mbuti | 0.289704 | 0.002436 | 118.931 | 411792 |
| GRA | Basque | Mbuti | 0.289597 | 0.002465 | 117.506 | 411738 |
| GRA | Denmark | Mbuti | 0.289556 | 0.002417 | 119.794 | 411782 |

| | | | | | | |
|---|---|---|---|---|---|---|
| CEN | Basque | Mbuti | 0.290093 | 0.002473 | 117.29 | 411734 |
| CEN | WBR | Mbuti | 0.289878 | 0.002457 | 117.985 | 411787 |
| CEN | Dyfed | Mbuti | 0.289809 | 0.002468 | 117.447 | 411781 |
| CEN | NIreland | Mbuti | 0.289612 | 0.002453 | 118.082 | 411785 |
| CEN | Cornwall | Mbuti | 0.289558 | 0.002453 | 118.049 | 411776 |
| SLO | Basque | Mbuti | 0.290673 | 0.002453 | 118.474 | 411707 |
| SLO | NIreland | Mbuti | 0.290111 | 0.002436 | 119.093 | 411783 |
| SLO | WBR | Mbuti | 0.290106 | 0.002428 | 119.466 | 411778 |
| SLO | Dyfed | Mbuti | 0.290056 | 0.002446 | 118.583 | 411777 |
| SLO | Cornwall | Mbuti | 0.289928 | 0.002429 | 119.383 | 411773 |
| NOU | Basque | Mbuti | 0.290438 | 0.002446 | 118.716 | 411713 |
| NOU | WBR | Mbuti | 0.289762 | 0.002418 | 119.857 | 411786 |
| NOU | Dyfed | Mbuti | 0.289602 | 0.002427 | 119.33 | 411778 |
| NOU | NIreland | Mbuti | 0.289553 | 0.00242 | 119.649 | 411774 |
| NOU | Cornwall | Mbuti | 0.289398 | 0.002422 | 119.48 | 411779 |

**Table S2.1 - Summary of the ancient DNA Medieval samples from France sequenced in this study**

| Samples ID | Date (CE) | Place | Mapped reads | Genome cov.* | Gender** | %endog. | X-contam. | mtDNA contam. | HOA sites | total sites HOA | mean cov. |
|---|---|---|---|---|---|---|---|---|---|---|---|
| fra001 | 340-535 | Saint Lupien Rezé | 271,093,228 | 0.48 | XX | 0.199312 | NA | 0.01 | 593124 | 247020 | 0.551608 |
| fra004 | 394-545 | Saint Lupien Rezé | 15,241,917 | 0.28 | XY | 0.057764 | 0.04367 | 0.02 | 593124 | 169051 | 0.344645 |
| fra008 | 943 - 1024 | Chaussé Saint Pierre - Angers | 5,983,204 | 0.08 | consistent with XY but not XX | 0.018284 | 0.22336 | 0.01 | 593124 | 64438 | 0.116075 |
| fra009 | 414-548 | Chaussé Saint Pierre - Angers | 26,781,466 | 0.36 | XY | 0.276266 | 0.03787 | 0.01 | 593124 | 199735 | 0.421043 |
| fra016 | 600-700 | Chémeré | 234,557,159 | 3.43 | consistent with XY but not XX | 0.480058 | 0.014 | 0.01 | 593124 | 574582 | 4.153501 |
| fra017 | 600-700 | Chémeré | 6,559,927 | 0.14 | XY | 0.020224 | 0.10887 | 0.01 | 593124 | 92736 | 0.170474 |

* after MapQ 30 filtering

** estimated biological sex from alignment data (1).

%endog. - % of endogenous DNA.

Contam. - contamination; Cov. - Coverage

| **Table S2.2 - $f_4$-statistics of the form *$f_4$(Mbuti (W); ancient EUR sample; sLoire_France_3-4cCE (Y), sLoire_France_6-7cCE (Z))*** | | | | | | | | |
|---|---|---|---|---|---|---|---|---|
| **Pop1** | **Pop2 (X) :** | **Pop3** | **Pop4** | **f4-stat** | **Z** | **BABA** | **ABBA** | **# SNPs** |
| Mbuti | Iberia_Medieval_published | Y | Z | 0.01 | 1.495 | 7351 | 7206 | 121038 |
| Mbuti | Scotland_MBA | Y | Z | 0.006 | 1.288 | 14059 | 13892 | 231058 |
| Mbuti | Italy_North_BellBeaker | Y | Z | 0.0059 | 1.154 | 13399 | 13242 | 219922 |
| Mbuti | Germany_EBA_Unetice | Y | Z | 0.005 | 1.11 | 15539 | 15383 | 255333 |
| Mbuti | Germany_BA.SG | Y | Z | 0.0141 | 0.984 | 1666 | 1620 | 27578 |
| Mbuti | England_MBA | Y | Z | 0.0036 | 0.952 | 15790 | 15677 | 259625 |
| Mbuti | Switzerland_BellBeaker | Y | Z | 0.0086 | 0.908 | 3806 | 3741 | 63565 |
| Mbuti | Iceland_Early_Christian.SG | Y | Z | 0.0048 | 0.763 | 8861 | 8776 | 151207 |
| Mbuti | Iceland_Pre_Christian.SG | Y | Z | 0.0026 | 0.694 | 16058 | 15974 | 267000 |
| Mbuti | France_BellBeaker | Y | Z | 0.0031 | 0.656 | 13879 | 13793 | 227996 |
| Mbuti | Germany_Lech_BellBeaker | Y | Z | 0.0026 | 0.589 | 15246 | 15167 | 251703 |
| Mbuti | Netherlands_BellBeaker | Y | Z | 0.0023 | 0.544 | 14636 | 14570 | 241118 |
| Mbuti | Germany_CordedWare | Y | Z | 0.002 | 0.481 | 16034 | 15969 | 264282 |
| Mbuti | Germany_EMedieval.SG | Y | Z | 0.0017 | 0.426 | 16197 | 16143 | 267044 |
| Mbuti | Vikings | Y | Z | 0.0015 | 0.42 | 16210 | 16162 | 267083 |
| Mbuti | Czech_BellBeaker | Y | Z | 0.0012 | 0.343 | 16023 | 15983 | 264009 |
| Mbuti | Italy_Medieval_EarlyModern_oCentralEuropean.SG | Y | Z | 0.0012 | 0.33 | 16142 | 16101 | 266252 |
| Mbuti | Iberia_IA | Y | Z | 0.0019 | 0.318 | 12802 | 12755 | 210115 |
| Mbuti | Italy_Imperial.SG | Y | Z | 0.0009 | 0.261 | 16009 | 15979 | 266257 |
| Mbuti | England_BellBeaker | Y | Z | 0.0009 | 0.259 | 16015 | 15984 | 263943 |
| Mbuti | England_LIA.SG | Y | Z | 0.0011 | 0.199 | 16154 | 16118 | 266869 |
| Mbuti | Germany_Lech_EBA | Y | Z | 0.0006 | 0.173 | 15896 | 15876 | 262151 |
| Mbuti | Italy_North_EarlyMedieval_Langobards | Y | Z | 0.0004 | 0.11 | 15936 | 15923 | 263943 |
| Mbuti | Iberia_Celtiberian | Y | Z | 0.0005 | 0.094 | 13661 | 13648 | 225336 |
| Mbuti | Germany_BellBeaker | Y | Z | 0.0003 | 0.082 | 16070 | 16060 | 265384 |
| Mbuti | Poland_BellBeaker | Y | Z | 0.0002 | 0.049 | 14394 | 14387 | 237819 |
| Mbuti | Iberia_EBA | Y | Z | 0.0002 | 0.02 | 7333 | 7331 | 121059 |
| Mbuti | England_Saxon.SG | Y | Z | 0.0001 | 0.014 | 16205 | 16203 | 267081 |
| Mbuti | Iberia_BA | Y | Z | -0.0001 | -0.033 | 15942 | 15946 | 262855 |

| | | | | | | | | |
|---|---|---|---|---|---|---|---|---|
| Mbuti | England_C_EBA | Y | Z | -0.0005 | -0.138 | 15623 | 15639 | 258037 |
| Mbuti | England_LBA | Y | Z | -0.0014 | -0.153 | 4239 | 4251 | 70203 |
| Mbuti | Germany_LBA_Halberstadt_published | Y | Z | -0.0015 | -0.249 | 14535 | 14579 | 240663 |
| Mbuti | Iberia_C_BA | Y | Z | -0.0018 | -0.251 | 8718 | 8749 | 144225 |
| Mbuti | England_Roman.SG | Y | Z | -0.0012 | -0.288 | 16127 | 16166 | 266546 |
| Mbuti | Ireland_BA.SG | Y | Z | -0.0016 | -0.337 | 16156 | 16206 | 267075 |
| Mbuti | Germany_Lech_MBA | Y | Z | -0.0027 | -0.382 | 6115 | 6147 | 102329 |
| Mbuti | Italy_North_Remedello_C.SG | Y | Z | -0.0032 | -0.461 | 8653 | 8708 | 148057 |
| Mbuti | Scotland_C_EBA | Y | Z | -0.0022 | -0.465 | 13781 | 13841 | 228477 |
| Mbuti | Iberia_MBA.SG | Y | Z | -0.0023 | -0.486 | 15137 | 15207 | 251643 |
| Mbuti | Iberia_BellBeaker | Y | Z | -0.0022 | -0.536 | 15765 | 15834 | 260421 |
| Mbuti | Scotland_LBA | Y | Z | -0.0026 | -0.577 | 14379 | 14455 | 237778 |
| Mbuti | Hungary_EBA_BellBeaker | Y | Z | -0.0024 | -0.614 | 15526 | 15602 | 256438 |
| Mbuti | Iberia_C | Y | Z | -0.0024 | -0.634 | 15959 | 16037 | 263490 |
| Mbuti | England_IA.SG | Y | Z | -0.0047 | -0.652 | 8691 | 8773 | 144659 |
| Mbuti | Italy_Medieval_EarlyModern.SG | Y | Z | -0.0024 | -0.685 | 15981 | 16059 | 266257 |
| Mbuti | Italy_C_BA.SG | Y | Z | -0.0034 | -0.748 | 15860 | 15969 | 262982 |
| Mbuti | Denmark_BA.SG | Y | Z | -0.0111 | -0.811 | 1702 | 1740 | 29356 |
| Mbuti | Italy_C.SG | Y | Z | -0.0044 | -0.894 | 16003 | 16143 | 265752 |
| Mbuti | Iberia_Roman | Y | Z | -0.0044 | -1.001 | 14989 | 15121 | 250637 |
| Mbuti | Scotland_BellBeaker | Y | Z | -0.0056 | -1.009 | 12120 | 12256 | 201217 |
| Mbuti | England_LIA_ERoman.SG | Y | Z | -0.0071 | -1.024 | 9452 | 9587 | 156401 |
| Mbuti | Germany_LRoman.SG | Y | Z | -0.0064 | -1.105 | 15085 | 15279 | 251566 |
| Mbuti | Italy_North_Remedello_EBA.SG | Y | Z | -0.0141 | -1.237 | 2755 | 2834 | 47322 |
| Mbuti | Denmark_LBA.SG | Y | Z | -0.0183 | -1.285 | 1329 | 1378 | 22809 |
| Mbuti | Wales_C.SG | Y | Z | -0.0122 | -1.695 | 7435 | 7620 | 124087 |
| Mbuti | France_BellBeaker_lowSteppe | Y | Z | -0.0119 | -1.919 | 10637 | 10894 | 177505 |

Note: Significant differences in allele sharing would result in |Z| > 3, which was not found for any of the above comparisons.

**Table S2.3 - qpAdm p-values for one-way models for modern and Mediaeval French**

| | WBR | EBP | SLO | NOR | HAU | GRA | CEN | NOU | Mediaeval French (300-1100 CE) |
|---|---|---|---|---|---|---|---|---|---|
| England_Roman | **0.9362** | **0.4472** | 0.0008 | **0.4504** | **0.3048** | 0.0004 | 0.0017 | 0.0000 | **0.1011** |
| England_Saxon | 0.0002 | 0.0000 | 0.0000 | 0.0000 | 0.0000 | 0.0000 | 0.0000 | 0.0000 | 0.0002 |
| Germany_EMedieval | **0.0744** | **0.1378** | 0.0000 | **0.0872** | 0.0054 | 0.0000 | 0.0000 | 0.0000 | **0.0756** |
| Iberia_Celtiberian | 0.0001 | 0.0000 | 0.0008 | 0.0002 | 0.0004 | 0.0000 | 0.0004 | 0.0028 | 0.0470 |
| Iberia_Medieval_published | 0.0000 | 0.0004 | 0.0366 | 0.0004 | 0.0009 | 0.0277 | 0.0219 | **0.0696** | **0.2434** |
| Iberia_Carolingian | **0.2219** | **0.4450** | **0.5731** | **0.5007** | **0.5409** | **0.4385** | **0.5462** | **0.6125** | **0.9324** |
| Iberia_Visigoth_Barcelona | **0.0689** | **0.1855** | **0.3636** | **0.1046** | **0.3221** | **0.3306** | **0.3678** | **0.4429** | **0.7020** |
| Iberia_Visigoth_Girona | 0.0363 | **0.2358** | **0.9075** | **0.3527** | **0.3833** | **0.7244** | **0.5947** | **0.7063** | **0.9538** |
| Iceland_Early_Christian | **0.3617** | **0.2317** | 0.0280 | **0.1455** | **0.1442** | 0.0442 | 0.0219 | 0.0009 | **0.2354** |
| Iceland_Pre_Christian | 0.0000 | 0.0000 | 0.0000 | 0.0000 | 0.0000 | 0.0000 | 0.0000 | 0.0000 | 0.0000 |
| Italy_Medieval_EarlyModern_oCentralEuropean | 0.0000 | 0.0002 | **0.4682** | 0.0003 | 0.0002 | **0.6033** | **0.3087** | **0.0695** | **0.8164** |
| Italy_Medieval_EarlyModern | 0.0000 | 0.0000 | 0.0000 | 0.0000 | 0.0000 | 0.0000 | 0.0000 | 0.0000 | 0.0000 |
| Italy_North_EarlyMedieval_Langobards | 0.0000 | 0.0000 | 0.0000 | 0.0000 | 0.0000 | 0.0000 | 0.0000 | 0.0000 | **0.4315** |
| Vikings | 0.0039 | 0.0000 | 0.0000 | 0.0000 | 0.0000 | 0.0000 | 0.0000 | 0.0000 | 0.0027 |

Modern population/cluster acronyms: Western Brittany (WBR), Eastern Brittany/*Pays-de-la-Loire* (EBP), south Loire (SLO), Normandy *(*NOR*)*; *Hauts-de-France* (HAU); *Grand Est (*GRA*)*; *Centre-Val de Loire (*CEN*)*; *Nouvelle-Aquitaine (*NOU*)*.

**Table S2.4. qpAdm results for the three-way model assuming Steppe pastoralists (SP), Early Farmers (EF) and Western Hunter-Gatherers (WHG) as sources populations of each modern and Mediaeval French populations (FRMedieval)**

| | SP | EF | WHG | | | |
|---|---|---|---|---|---|---|
| | Prop (std. error) | Prop (std. error) | Prop (std. error) | d.f | Chisq | p-value |
| WBR | 0.460 (0.021) | 0.436 (0.020) | 0.104 (0.010) | 10 | 77.265 | 1.72E-12 |
| EBP | 0.435 (0.021) | 0.476 (0.020) | 0.089 (0.009) | 10 | 69.372 | 5.86E-11 |
| SLO | 0.372 (0.021) | 0.545 (0.020) | 0.083 (0.010) | 10 | 63.73 | 7.09E-10 |
| NOR | 0.437 (0.021) | 0.480 (0.020) | 0.083 (0.010) | 10 | 51.303 | 1.54E-07 |
| HAU | 0.422 (0.021) | 0.483 (0.020) | 0.095 (0.010) | 10 | 67.78 | 1.19E-10 |
| GRA | 0.389 (0.020) | 0.548 (0.019) | 0.063 (0.009) | 10 | 64.295 | 5.53E-10 |
| CEN | 0.384 (0.020) | 0.538 (0.020) | 0.079 (0.010) | 10 | 70.677 | 3.28E-11 |
| NOU | 0.329 (0.019) | 0.578 (0.019) | 0.093 (0.009) | 10 | 59.818 | 3.92E-09 |
| FRMedieval | 0.326 (0.057) | 0.608 (0.054) | 0.067 (0.029) | 10 | 9.248 | 0.508768613 |

| **Table S2.5 - *f$_4$-statistics* of the form *f$_4$(Mbuti; ancient EUR sample; WBR, other modern French population)*** | | | | | | | | |
|---|---|---|---|---|---|---|---|---|
| Pop1 (W) | Pop2 (X) : | Pop3 (Y) | Pop4 (Z) | D-stat | Z | BABA | ABBA | # of SNPs |
| **Mbuti** | **Russia_EBA_Yamnaya_Samara** | **WBR** | **NOU** | **-0.0091** | **-11.381** | **27968** | **28481** | **464380** |
| **Mbuti** | **Russia_EBA_Yamnaya_Samara** | **WBR** | **CEN** | **-0.007** | **-8.586** | **28029** | **28422** | **464380** |
| **Mbuti** | **Russia_EBA_Yamnaya_Samara** | **WBR** | **SLO** | **-0.0069** | **-8.816** | **28020** | **28407** | **464380** |
| **Mbuti** | **Russia_EBA_Yamnaya_Samara** | **WBR** | **GRA** | **-0.0058** | **-7.646** | **28068** | **28398** | **464380** |
| **Mbuti** | **Russia_EBA_Yamnaya_Samara** | **WBR** | **HAU** | **-0.0053** | **-7.279** | **28085** | **28383** | **464380** |
| **Mbuti** | **Russia_EBA_Yamnaya_Samara** | **WBR** | **NOR** | **-0.0041** | **-4.895** | **28093** | **28326** | **464380** |
| **Mbuti** | **Russia_EBA_Yamnaya_Samara** | **WBR** | **EBR** | **-0.0041** | **-5.3** | **28108** | **28341** | **464380** |
| Mbuti | Germany_EN_LBK_published | WBR | HAU | -0.0004 | -0.47 | 28132 | 28152 | 463062 |
| Mbuti | Germany_EN_LBK_published | WBR | NOR | -0.0003 | -0.429 | 28103 | 28122 | 463062 |
| Mbuti | Germany_EN_LBK_published | WBR | EBR | 0.0003 | 0.37 | 28139 | 28123 | 463062 |
| Mbuti | Germany_EN_LBK_published | WBR | GRA | 0.0013 | 1.678 | 28189 | 28115 | 463062 |
| Mbuti | Germany_EN_LBK_published | WBR | CEN | 0.0021 | 2.66 | 28213 | 28096 | 463062 |
| **Mbuti** | **Germany_EN_LBK_published** | **WBR** | **NOU** | **0.0025** | **3.267** | **28231** | **28089** | **463062** |
| **Mbuti** | **Germany_EN_LBK_published** | **WBR** | **SLO** | **0.0026** | **3.384** | **28218** | **28070** | **463062** |
| Mbuti | Iberia_EN | WBR | HAU | -0.0009 | -1.089 | 27436 | 27484 | 452926 |
| Mbuti | Iberia_EN | WBR | EBR | -0.0001 | -0.179 | 27443 | 27451 | 452926 |
| Mbuti | Iberia_EN | WBR | NOR | 0.0006 | 0.721 | 27451 | 27417 | 452926 |
| Mbuti | Iberia_EN | WBR | GRA | 0.0013 | 1.654 | 27508 | 27434 | 452926 |
| Mbuti | Iberia_EN | WBR | CEN | 0.0015 | 1.831 | 27512 | 27428 | 452926 |
| Mbuti | Iberia_EN | WBR | SLO | 0.0021 | 2.638 | 27515 | 27402 | 452926 |
| **Mbuti** | **Iberia_EN** | **WBR** | **NOU** | **0.0032** | **3.944** | **27569** | **27392** | **452926** |
| **Mbuti** | **Luxembourg_Loschbour_published.DG** | **WBR** | **GRA** | **-0.0064** | **-5.314** | **28402** | **28769** | **466881** |
| **Mbuti** | **Luxembourg_Loschbour_published.DG** | **WBR** | **NOU** | **-0.0041** | **-3.444** | **28477** | **28713** | **466881** |
| Mbuti | Luxembourg_Loschbour_published.DG | WBR | CEN | -0.0034 | -2.883 | 28494 | 28689 | 466881 |
| Mbuti | Luxembourg_Loschbour_published.DG | WBR | HAU | -0.003 | -2.565 | 28504 | 28677 | 466881 |
| Mbuti | Luxembourg_Loschbour_published.DG | WBR | SLO | -0.0026 | -2.17 | 28513 | 28661 | 466881 |
| Mbuti | Luxembourg_Loschbour_published.DG | WBR | NOR | -0.0021 | -1.702 | 28508 | 28627 | 466881 |
| Mbuti | Luxembourg_Loschbour_published.DG | WBR | EBR | -0.0019 | -1.557 | 28533 | 28641 | 466881 |

Note: significant values (|Z| > 3) are indicated in bold; |Z| > 4 are indicated in yellow and |Z| > 6 are indicated in red. To test whether potentially increased WHG ancestry in WBR could lead to an over estimation of steppe-related ancestry in WBR relative to the other French populations, we computed $f_4$-statistics of the form f4(Mbuti, Loschbour; WBR, other modern French) using the Loschbour individual (7205±50 BP) to represent the WHG component. *f4* statistics of this form were not generally significant except when the other French population was either the NOU or the GRA. Therefore, the genetic affinities between WBR and ancestry Bell-Beaker-associated individuals carrying high-steppe ancestry does not seem to be caused by significantly increased WHG ancestry in WBR.

**Table S2.6 - *$F_4$-statistics* of the form *$f_4$(Mbuti; Megalithic samples; WBR, other modern French populations)***

| Pop1 (W) | Pop2 (X) : | Pop3 (Y) | Pop4 (Z) | D-stat | Z | BABA | ABBA | # of SNPs |
|---|---|---|---|---|---|---|---|---|
| Mbuti | Ireland_Megalithic.SG | WBR | English | -0.000207 | -1.648 | 28628 | 28726 | 471404 |
| Mbuti | Ireland_Megalithic.SG | WBR | HAU | -0.000153 | -1.59 | 28687 | 28759 | 471404 |
| Mbuti | Ireland_Megalithic.SG | WBR | GRA | -0.000106 | -1.092 | 28705 | 28755 | 471404 |
| Mbuti | Ireland_Megalithic.SG | WBR | NOR | -0.000054 | -0.527 | 28681 | 28707 | 471404 |
| Mbuti | Ireland_Megalithic.SG | WBR | EBP | -0.000031 | -0.325 | 28707 | 28722 | 471404 |
| Mbuti | Ireland_Megalithic.SG | WBR | SLO | 0.00009 | 0.916 | 28745 | 28703 | 471404 |
| Mbuti | Ireland_Megalithic.SG | WBR | CEN | 0.000095 | 0.96 | 28759 | 28714 | 471404 |
| Mbuti | Ireland_Megalithic.SG | WBR | NOU | 0.000196 | 2.023 | 28788 | 28696 | 471404 |
| Mbuti | Scotland_Megalithic.SG | WBR | HAU | -0.000144 | -1.007 | 9333 | 9355 | 154275 |
| Mbuti | Scotland_Megalithic.SG | WBR | GRA | -0.000104 | -0.72 | 9340 | 9356 | 154275 |
| Mbuti | Scotland_Megalithic.SG | WBR | English | -0.000133 | -0.684 | 9317 | 9338 | 154275 |
| Mbuti | Scotland_Megalithic.SG | WBR | NOR | -0.000088 | -0.577 | 9328 | 9342 | 154275 |
| Mbuti | Scotland_Megalithic.SG | WBR | EBP | -0.000079 | -0.555 | 9332 | 9344 | 154275 |
| Mbuti | Scotland_Megalithic.SG | WBR | CEN | -0.000014 | -0.091 | 9350 | 9352 | 154275 |
| Mbuti | Scotland_Megalithic.SG | WBR | SLO | 0.000103 | 0.695 | 9350 | 9334 | 154275 |
| Mbuti | Scotland_Megalithic.SG | WBR | NOU | 0.000316 | 2.098 | 9376 | 9327 | 154275 |

Note: Significant differences in allele sharing would result in $|Z| > 3$, which was not found for any of the above comparisons.

| Table S2.7 - $F_4$-statistics of the form *$f_4$(Mbuti; ancient EUR sample; WBR, other modern French population)* | | | | | | | | |
|---|---|---|---|---|---|---|---|---|
| **Pop1 (W)** | **Pop2 (X) :** | **Pop3 (Y)** | **Pop4 (Z)** | **D-stat** | **Z** | **BABA** | **ABBA** | **# of SNPs** |
| Mbuti | Germany_CordedWare | WBR | GRA | -0.00640 | -8.235 | 28300 | 28666 | 466547 |
| Mbuti | Germany_CordedWare | WBR | NOR | -0.00480 | -5.766 | 28320 | 28592 | 466547 |
| Mbuti | Germany_CordedWare | WBR | CEN | -0.00730 | -9.113 | 28269 | 28687 | 466547 |
| Mbuti | Germany_CordedWare | WBR | EBR | -0.00460 | -5.987 | 28343 | 28604 | 466547 |
| Mbuti | Germany_CordedWare | WBR | SLO | -0.00650 | -8.191 | 28285 | 28653 | 466547 |
| Mbuti | Germany_CordedWare | WBR | NOU | -0.00880 | -10.708 | 28224 | 28724 | 466547 |
| Mbuti | Germany_CordedWare | WBR | HAU | -0.00620 | -8.223 | 28302 | 28658 | 466547 |

| Table S2.8- $f_4$-statistics of the form *$f_4$(Mbuti; modern French population; Iberian Early Neolithic, Early Neolithic LBK )* | | | | | | | | |
|---|---|---|---|---|---|---|---|---|
| **Pop1 (W)** | **Pop2 (X) :** | **Pop3 (Y)** | **Pop4 (Z)** | **D-stat** | **Z** | **BABA** | **ABBA** | **# of SNPs** |
| Mbuti | EBP | Iberia_EN | Germany_EN_LBK_published | -0.000084 | -0.43 | 26520 | 26558 | 447696 |
| Mbuti | WBR | Iberia_EN | Germany_EN_LBK_published | -0.000141 | -0.716 | 26504 | 26567 | 447696 |
| Mbuti | NOR | Iberia_EN | Germany_EN_LBK_published | -0.000251 | -1.257 | 26470 | 26583 | 447696 |
| Mbuti | HAU | Iberia_EN | Germany_EN_LBK_published | -0.000063 | -0.317 | 26501 | 26529 | 447696 |
| Mbuti | GRA | Iberia_EN | Germany_EN_LBK_published | -0.000136 | -0.692 | 26512 | 26572 | 447696 |
| Mbuti | SLO | Iberia_EN | Germany_EN_LBK_published | -0.000056 | -0.279 | 26529 | 26553 | 447696 |
| Mbuti | CEN | Iberia_EN | Germany_EN_LBK_published | -0.000076 | -0.388 | 26523 | 26557 | 447696 |
| Mbuti | NOU | Iberia_EN | Germany_EN_LBK_published | -0.000214 | -1.098 | 26493 | 26589 | 447696 |

Note: Significant differences in allele sharing would result in |Z| > 3, which was not found for any of the above comparison.

### Supplementary Text

#### Supplementary archaeological details

***Rezé (Loire-Atlantique) - Saint-Lupien area***

Rezé is located in the *département* of *Loire Atlantique* on the southern shore of the Loire river facing the city of Nantes to the north. It is located at ~55km from the Atlantic coast. The excavation was carried out by the team of Mikaël Rouzic between 2005 and 2016 and the archaeological description of the sites remains unpublished.

Site: MR 4825
Samples: SP 4903 (fra001)
Radiocarbon dating: 340-535 CE

The wall of the site MR 4825 was pitted on several layers in order to install the SP 4903 grave with no signs of excavation. The grave was filled with a compact and heterogeneous brown silt, very rich in TCA, lime, gravel and coal. The grave contained the individual SP4903 and two ceramic shards were found inside. The individual was an adult who died at an age of 20-59 years.

Site: US 20380; US 20381; US 20382; US 20383
Samples: SP 20380 (fra004)
Radiocarbon Dating: 394-545 CE

The site consists of a 2.40 m long and 0.72 m wide burial pit, oriented east/west. I was dug in the ancient road and lies on top of an ash and charcoal layer. The fill is a compact and heterogeneous medium brown sandy loam containing numerous inclusions of architectural terracotta fragments, small blocks, some mortar fragments, several shards of ancient ceramics, glass and faunal bones. Six nails and eight shanks of carpentry nails were taken from the entire thickness of the filling. The skeleton remnants found inside belonged to a male individual 20-29 years old at death.

**Chéméré *(Loire-Atlantique)***

Samples: SP 46 and SP 55 (fra016 and fra017)
Radiocarbon Dating: 600-700 CE

The Merovingian necropolis of Chéméré (*Loire Atlantique*, south Nantes) is located in an area peripheral to the present village. Its occupation is dated to the 6th-7th centuries by the furniture deposited in the tombs and by radiocarbon-based analyses. The total surface area of the burial space is estimated to be around 7700 m². Approximately 1500 m² of land containing 416 tombs have been cleared since the 1960s. The latest excavations in 2007 yielded a population sample of 181 individuals (155 adults and 26 children) recognised by anthropological analysis as a morphologically homogenous group. The two men 46 and 55 are from this group. They were buried in the same row, at the southern end of the cemetery.

***Angers (Maine-et-Loire) - Chaussé Saint Pierre***
Samples: SP 1046 and SP 1049A (fra008 and fra009)
Radiocarbon Dating: 414-1024 CE

The site of Chaussé Saint-Pierre is located right in the modern city-center of Angers. The city of Angers is located on the intersection of Maine and the Loire rivers and on the northern shore of the latter. It is situated approximately 100km upstream from Nantes. The excavations in this site, which were led by the team of Martin Pithon, from the INRAP of west France, took place between July and August 2009.
The large density of archaeological remains around the Chaussé Saint-Pierre suggests it was densely occupied from the first period of the Roman Empire (27 BCE-250 CE). Until the 4th century CE the area was occupied by commercial buildings and thermes, nevertheless by the 5th century archaeological remains associated with a necropolis indicate a shift in the pattern of occupation. The excavation of Rue Chaussée-Saint-Pierre focused on the north-eastern end of the hopper where the project was the deepest and where the remains were best preserved. The excavations investigated a stratigraphic sequence of almost 1 m thick between the modern-day earthworks and the natural terrain over an area of ~100 $m^2$. The analysis of this sequence shows five main phases of occupation which lie within a chronological interval between the beginning of the Roman Empire (20 BC to 15 AD) and the modern period. Funerary-associated occupation begins in phase three.

**SP 1049A (fra009)**

The burial SP 1049 is among the earliest among those belonging to the phase three of occupation and the dating ranges from the beginning of the 5th century to the first half of the 6th century. This tomb, which has been badly damaged by Medieval/modern construction, seems to be part of a homogeneous group of six burials very similar in nature. The dating of the tomb shows repeated burial and skeletal remains indicate the presence of a total of three individuals, two adults and one child. The child's skeleton and one skull not belonging to the child have been dated to 401-539 CE and 414-548 CE, respectively. The skeleton remains indicate the child (fra009) was buried at the age of 1-4 years old. The other skeletal remains seem to belong to a female and an individual of indeterminate gender aged of >25 and >30 years, respectively. The time proximity of the burials within the tomb are likely indicative of a voluntary grouping and may represent a family management of the deposits.

**SP 1046 (fra008)**

The tomb SP 1046 together with a set of the other tombs dating from the 13-15th century CE indicate the practice of reuse of the tombs. The burial site is composed of heterogeneous loose deposits of fine sandy sediment including charcoal, fragments of tiles/bricks, blocks of slate shale as well as a batch of ceramic sherds mixing of Antiquity and Medieval (High Middle Ages? and Classical Middle Ages) elements. A full skeleton (fra008), belonging to a male aged >30 years, and other skeletal remains likely to come from three different individuals were buried in this site. Radiocarbon dating indicates the skeleton dates to 943 - 1024 CE.

### Supplementary Discussion

#### 1. Geolinguistics and genetic clusters

We can find some correlations between dialectal areas and genetic clusters. However, these are not considered as biological relationships. If they correlate, it is on the basis of long-term inertia and socio-economic complex and cultural factors. Below the linguistic data come from the *Nouvel Atlas Linguistique de Basse-Bretagne* (Le Dû 2001) and the place names are extracted from the IGN BDtopo® database. A deeper understanding of such correlations requires a higher resolution with respect to the genetic data.

##### 1.1. Cluster “Bretagne-Centre”, cluster “Cornouaille” and cluster “Vannes” together

The cluster “Bretagne-Centre” can be compared to the dialectal area featured by the use of two initial consonants: the aspirated [h] instead of an unaspirated and the alveolar fricative [z] instead of [s], stretching respectively from the northern coast to the southern sea-side of the central area (Fig.S1.10). The river Blavet agrees with the south-eastern limit separating the cluster “Bretagne-Centre” from the cluster “Vannes”.
Fig. S1.10 ‘h-’ shows seven analysed NALBB maps considering the treatment of the initial [h]. Fig. S1.10 ‘z-’ represents the area where the initial [s] is voiced as [z]. The [z] area extends in a central crossing zone from the North to the South. It gradually disappears towards the southwest[1].
These two phonological features extend over an area between the southern coast of Morbihan and the north of the Côtes-d'Armor, excluding a north-western and a south-eastern area without the initial [h] and where the initial [s] is pronounced. These two facts correlate with the distribution of “Bretagne-Centre” and “Cornouaille” clusters. The ‘z-’ map seems to correlate better than the ‘h-’ one. The articulation [z] for the initial [s] is also observed on the other side of the Channel, in the south-western dialects of England: *zome* 'some', *Zomerzet* 'Somerset', *to zwaip* for *to* sweep' fr. 'balayer', *zwell* for 'swell', *zwate* / *zweet* 'sweet' fr. 'doux' (Tristram 1995, Markus 2021). At this point, it would be interesting to compare the data from the People of the British Isles project (2,3) which shows a differentiation between the Cornwall and Devon clusters.

##### 1.2. Cluster “Cornouaille”

If we would look for a coherence between the distribution of dialectal facts in Lower-Brittany and “Cornouailles” cluster, we could consider the palatalization of -*h*- [h] into -*y*- [j] given their broad overlap (Fig. S1.11). This phenomenon can be found in the following words: *meryed* “girls”, *luyadenn* “lightning”, *luyed* “lightnings” and *meryer / dimeryer* 'Wednesday (noun/adv)'.

##### 1.3. Clusters ‘Léon’, ‘Cornouaille’, ‘Bretagne-Centre’, ‘Vannes’ and ‘Guérande’ and the Celto-Romance limit in eastern Brittany

[1] A first analysis of the initial [z], according to NALBB data, was carried out by François Falc'hun, 1981, *Perspectives nouvelles sur l'histoire de la langue bretonne,* Union Générale d'Editions, fig. 67 p. 204.

All together the clusters “Léon”, “Cornouaille”, “Bretagne-Centre”, “Vannes” and “Guérande” correlate with the Celto-Romance limit attested between the 16th and the 18th centuries. The distribution of place-names Ker- attests the progressive withdrawal towards the west of this eastern limit of Celtic dialects (Fig. S5). According to Bernard Tanguy, the oriental limit of toponyms Ker- does not show the maximum extension of Breton spoken varieties but testifies the retreat of Armorican Gaulish dialects facing the gradual advance of the Gallo-Romance dialects in formation during the Early Middle Ages (4). The same toponymic element can be found on both sides of the Channel spelled *Ker-*, *Car-*, *Gear* in Cornwall (Joanne Pye 2019), and *Caer* in Wales (National Library of Wales 2021), Great Britain. So far, there is no evidence that the presence of *Ker-* place names in Brittany is due to a migration from Great Britain. If one considers the density of place-names as indicating a point of departure, it would indicate the opposite. At this stage it is more cautious to simply consider this toponymic element as a celtic linguistic and cultural mark.

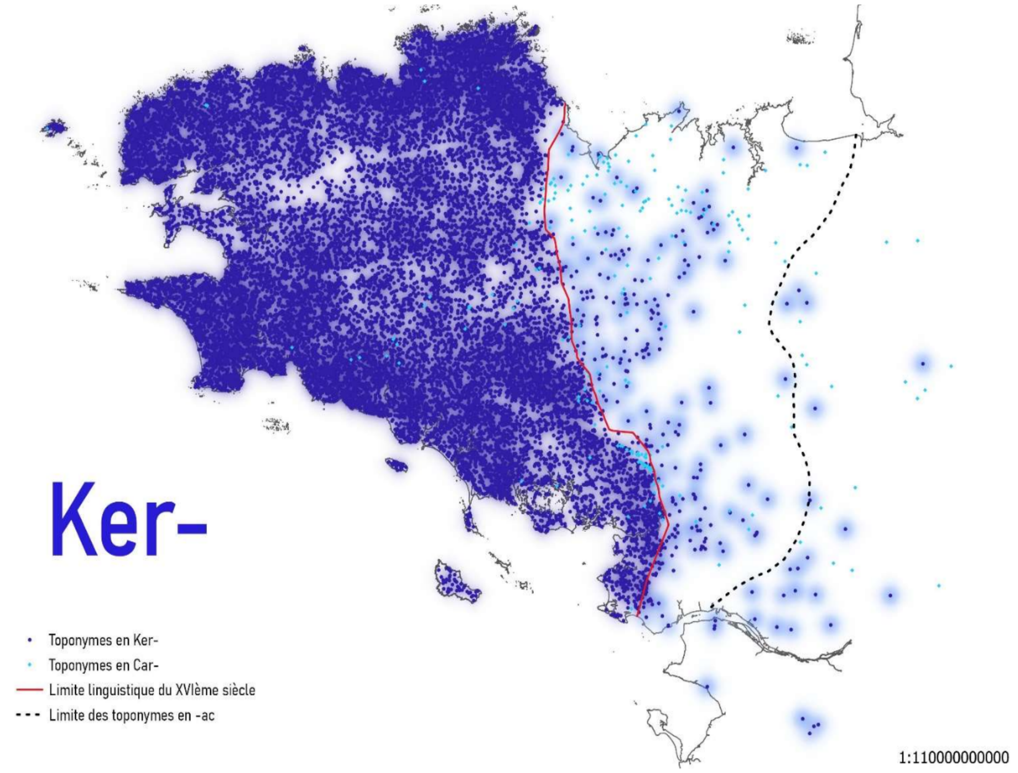


**Figure S5 - Distribution of *Ker-* “inhabited place”.** Data : BDtopo®, National Geographic Institute (IGN). *Ker-* (royal blue) and *Car-* (turquoise) toponyms.

#### 2. The role of rivers in the distribution of the 18 clusters

Rivers and streams could play a role in the diffusion of linguistic features. In France, the Loire initially formed an important border between the Oïl Gallo-Romance dialects in the North, and the Oc Gallo-Romance dialects in the South. Later, it became a vector for the diffusion of new traits coming from the North to the inner country (5). In Lower Brittany, on a smaller scale, the river Blavet seems to be at the origin of a bundle of isophones separating the different treatments of the interdental spirants (6). The linguistic frontier separating the Celtic and Gallo-Romance areas is not determined by a natural barrier. Nevertheless, the hydrography seems to mark the territories within these zones. By using a geographical information system and the file of the main rivers proposed by the IGN's BDcarthage®, we assessed their role in the organisation of the k=18 genetic clusters.

We first discuss the rivers which seem to correspond to the limits of the genetic clusters. Cluster “Léon” is bordered in the north-east by the River of Morlaix and the Jarlot. In the South, the same cluster is separated from cluster “Cornouaille” by the river Aulne. The Laïta, extended by the Ellé until the north of Le Faouët, seems to border the cluster “Cornouaille”. The Carnoët forest could have reinforced its natural limit in the South. Cluster “Vannes” is enclosed on its western side by the Blavet and on its north side by the Oust. Cluster “Nantes” is circumscribed by the Semnon in the north and the Vilaine in the west. Cluster “Malo-Rennais” is surrounded by the Gouessant, then by the Yvel and the Hivet in the west, by the Canut and the Semnon further south. Finally, the Loire delimits the clusters “Guérande”, “Nantes” and “Ancenis” in the north and the clusters “Retz”, “Mauges 3” and “Mauges 1” in the south. In South Loire Vendée, we observe a role in cluster landscape for rivers like “Le Lay” and “Sèvre Nantaise”, although it appears to be less clear. The “Maine-Anjou” cluster is crossed by several rivers and, when focusing at a finer, 78 clusters, scale, we also observe structuration along rivers like Mayenne, Vègre and Loir (data not shown).

The difference between navigable and non-navigable waterways does not seem to be decisive when considering the role of rivers by hampering gene flow. However, it has to be taken into account when studying their role as vectors. The geographical distribution of “Maine-Anjou" includes the rivers Loire, Sarthe, Mayenne and Oudon, all of them are navigable waterways according to the *Service d'administration nationale des données et référentiels sur l'eau* (Sandre and IGN 2017). The cluster “Guérande” seems to be contained by the Vilaine mainly, the Oust, the Arz and the Brivet in the south. When we consider river geography, the distribution of clusters seems to be more complex than a simple cluster/bishoprics correlation.

**Computer simulations to access the power to detect a short bottleneck with IBDNe**

Identity-by-descent (IBD) sharing in population samples can be used to estimate recent effective population size changes. IBDNe (7) assumes an idealised random mating population with a constant effective size that has similar random changes in allele frequencies over time. Shorter IBD segments represent coalescent events that occurred further back in time, while longer segments represent coalescent events that occurred in the past few generations. If the number of IBD segments is high, a larger number of coalescent events have occurred, indicating a high coalescence probability meaning a low effective size. Similarly, if the number of IBD segments is low, the effective size is high.

The effective population size trajectories inferred in this study for the three main genetic clusters inferred with fineSTRUCTURE indicate a slight and short population decline, starting around the years 1,230-1,350 CE (13 generations and assuming a generation time ~29 year/gen) and lasting for almost ~300 years (~10 generations). Given the assumption of random mating within samples and our knowledge of extensive fine-scale structure within Northwestern France, this motivated us to perform a simulation study to assess whether a bottleneck could be detected if the evolutionary history of the population involves other scenarios other than a bottleneck.

More specifically, we investigated: 1) The performance of IBDNe in detecting a recent bottleneck; 2) The effect of having admixture (punctual migration), sample structure and exponential growth in the estimation.

We used ARGON (8), a discrete time Wright-Fisher model simulator. Data was simulated for 50 diploid individuals (per population) according to a model containing 2 populations (see below). The data was simulated for each chromosome according to its respective size and we used the human recombination map (1000Genomes project). We ran the IBDNe software on our data only using IBD segments that were 4cM in size. Simulated scenarios are shown below (Fig. S6).

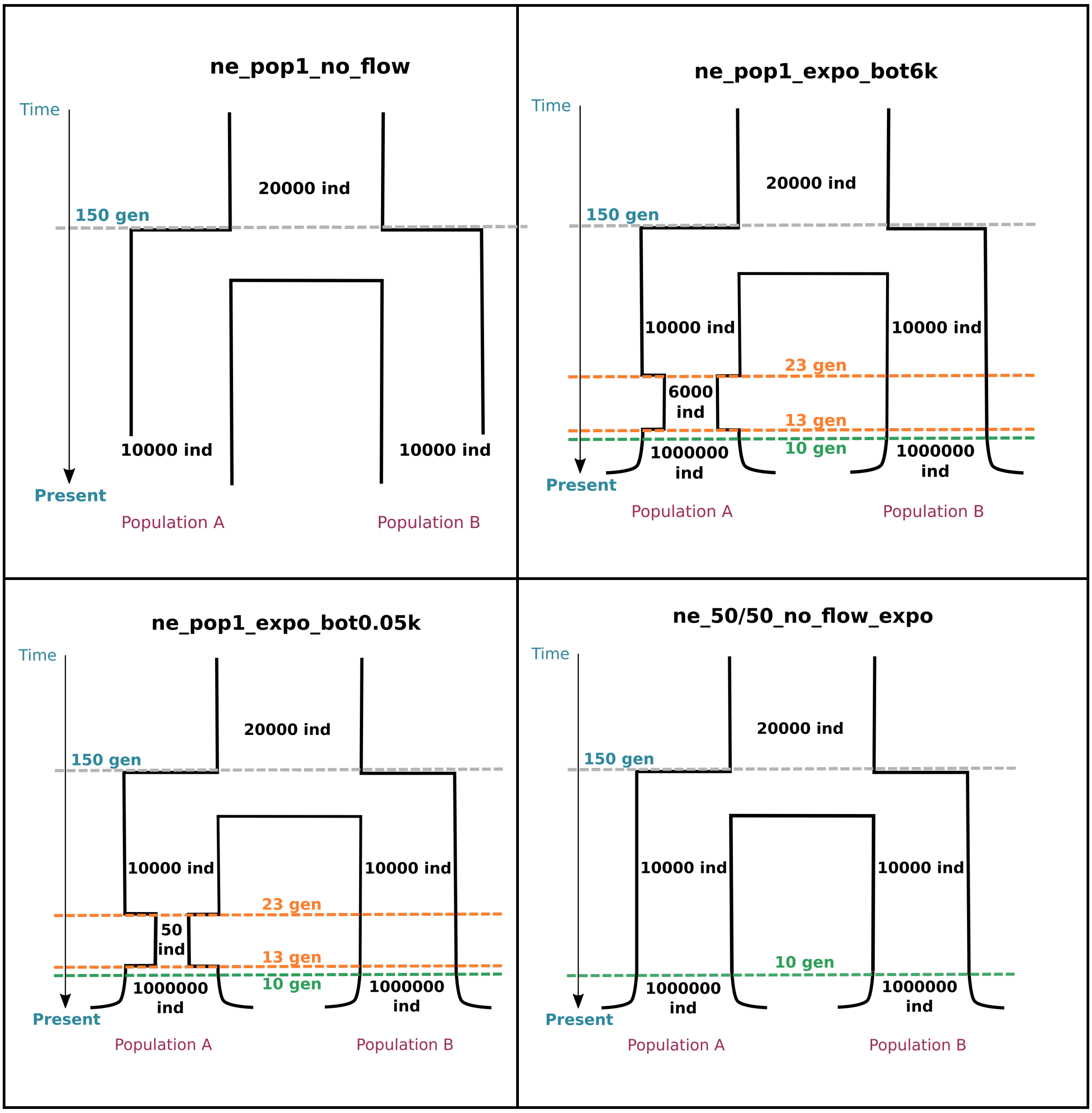

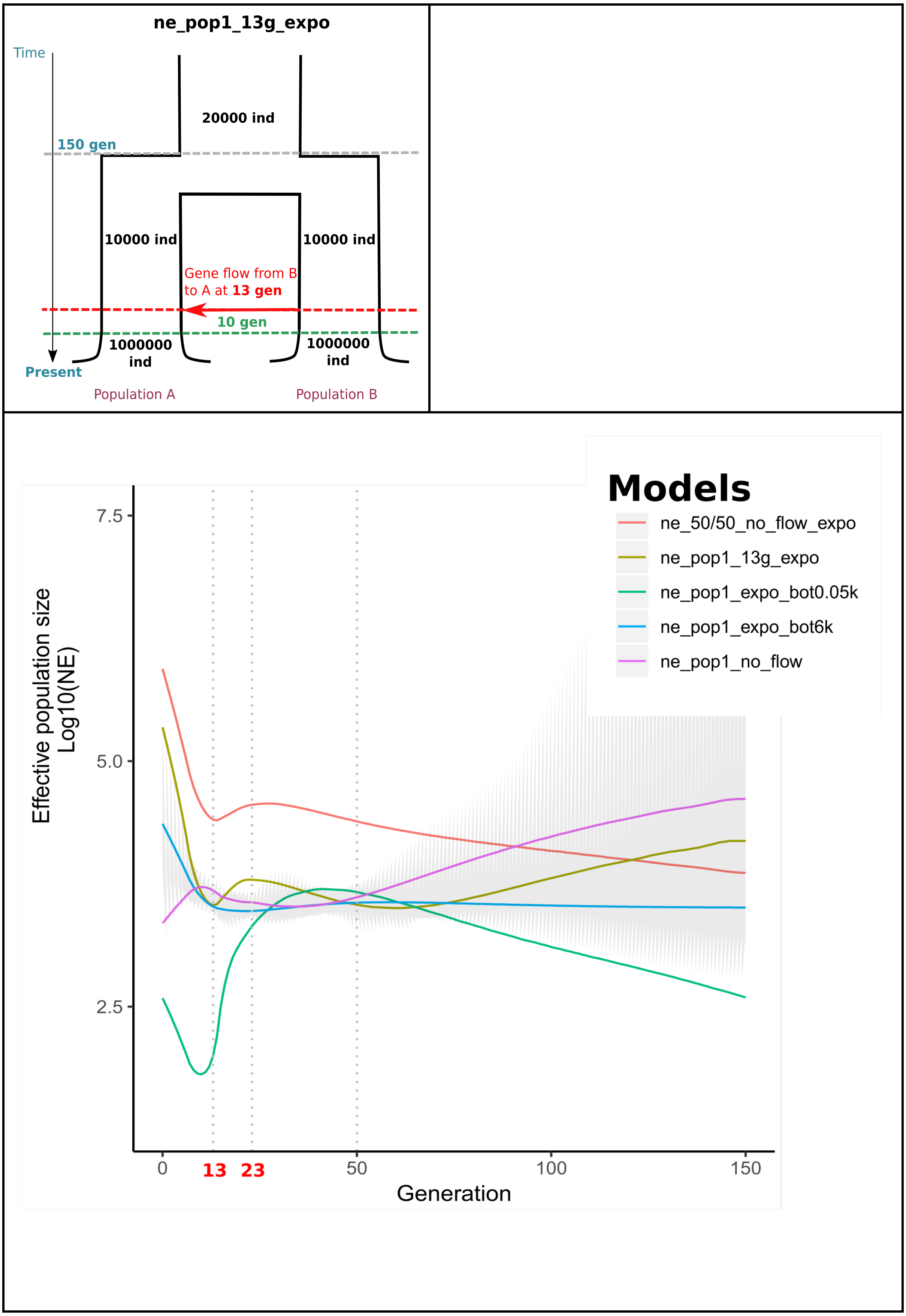


**Fig. S6 - Simulation study performed to assess the power of IBDNe to infer short-lasting bottlenecks**. We simulated models involving two populations that diverged 150

generations ago. Ne trajectories were inferred with IBDNe only from Population 1. The upper left panel represents the simplest model, in which after divergence Population 1 and 2 kept constant *Ne*. ne_pop1_expo_bot6k and ne_pop1_expo_bot0.05k models are variations of the simplest model, in which Population 1 undergoes a 10 generation long bottleneck after diverging from Population 2. Bottleneck starting times were set to 23 generations and the reduction in *Ne* were set to 6000/10000 and 50/10000, respectively. In model ne_50/50_no_flow_expo, Population 1 did not suffer a reduction in *Ne* but the sample analysed with IBDNe resulted from randomly sampling 50 individuals from both Population 1 and 2 in order to mimic population structure within the sample. Finally, the model ne_pop1_13g_flow_expo does not include a bottleneck but rather a one-generation event of gene flow. According to this model at generation 13 Population 1 has 10% of its lineages going to Population 2 (backwards in time). All models but the simplest one (upper left panel) include exponential growth in the last 10 generations as previously inferred in European populations (7,9).

We see that both the presence of stratification within the analysed sample represented here by the model ne_50/50_no_flow_expo (coral, Fig. S6) and the presence of admixture at generation 13 represented here by the model ne_pop1_13g_expo (kakhi, Fig. S6) induces a signal of a slight bottleneck similar to that found in the real dataset, with the drop in Ne and its recovery takes place in ~10 generations. In the case of an actual bottleneck, with a severe bottleneck represented here by the model ne_pop1_expo_bot0.05k (green colour, Fig. S6) and the bottleneck model (ne_pop1_expo_bot6k, turquoise colour, Fig. S6) with a reduction in size of ~⅓ of the original Ne, which is thought to have occurred during the black death period, we see that the actual start of the bottleneck is offset compared to the real timeframe (starting 23gen ago). Therefore, we caution that short-lasting bottleneck signals from IBDNe trajectories may represent population structure or recent gene flow rather than a real bottleneck, which are inferred to exhibit a more progressive decrease and recovery than the reality.
